# Examining Functional Linkages Between Conformational Dynamics, Protein Stability and Evolution of Cryptic Binding Pockets in the SARS-CoV-2 Omicron Spike Complexes with the ACE2 Host Receptor: Recombinant Omicron Variants Mediate Variability of Conserved Allosteric Sites and Binding Epitopes

**DOI:** 10.1101/2023.09.11.557205

**Authors:** Mohammed Alshahrani, Grace Gupta, Sian Xiao, Peng Tao, Gennady Verkhivker

**Affiliations:** Keck Center for Science and Engineering, Graduate Program in Computational and Data Sciences, Schmid, College of Science and Technology, Chapman University, Orange, CA 92866, United States of America; Department of Biomedical and Pharmaceutical Sciences, Chapman University School of Pharmacy, Irvine, CA 92618, United States of America; Department of Chemistry, Center for Research Computing, Center for Drug Discovery, Design, and Delivery (CD4), Southern Methodist University, Dallas, Texas, 75275, United States of America

## Abstract

In the current study, we explore coarse-grained simulations and atomistic molecular dynamics together with binding energetics scanning and cryptic pocket detection in a comparative examination of conformational landscapes and systematic characterization of allosteric binding sites in the SARS-CoV-2 Omicron BA.2, BA.2.75 and XBB.1 spike full-length trimer complexes with the host receptor ACE2. Microsecond simulations, Markov state models and mutational scanning of binding energies of the SARS-CoV-2 BA.2 and BA.2.75 receptor binding domain complexes revealed the increased thermodynamic stabilization of the BA.2.75 variant and significant dynamic differences between these Omicron variants. Molecular simulations of the SARS-CoV-2 Omicron spike full length trimer complexes with the ACE2 receptor complemented atomistic studies and enabled an in-depth analysis of mutational and binding effects on conformational dynamic and functional adaptability of the Omicron variants. Despite considerable structural similarities, Omicron variants BA.2, BA.2.75 and XBB.1 can induce unique conformational dynamic signatures and specific distributions of the conformational states. Using conformational ensembles of the SARS-CoV-2 Omicron spike trimer complexes with ACE2, we conducted a comprehensive cryptic pocket screening to examine the role of Omicron mutations and ACE2 binding on the distribution and functional mechanisms of the emerging allosteric binding sites. This analysis captured all experimentally known allosteric sites and discovered networks of inter-connected and functionally relevant allosteric sites that are governed by variant-sensitive conformational adaptability of the SARS-CoV-2 spike structures. The results detailed how ACE2 binding and Omicron mutations in the BA.2, BA.2.75 and XBB.1 spike complexes modulate the distribution of conserved and druggable allosteric pockets harboring functionally important regions. The results of are significant for understanding functional roles of druggable cryptic pockets that can be used for allostery-mediated therapeutic intervention targeting conformational states of the Omicron variants.

## Introduction

The enormous wealth of structural and biochemical studies have revealed mechanisms of the SARS-CoV-2 viral spike (S) glycoprotein consisting of the flexible amino (N)-terminal S1 subunit experiencing functional motions and structurally rigid carboxyl (C)-terminal S2 subunit. Conformational transformations of the SARS-CoV-2 S protein between the distinct closed and open states are driven by global movements of the S1 subunit consisting of an N-terminal domain (NTD), the receptor-binding domain (RBD), and two structurally conserved subdomains SD1 and SD2 which coordinate their dynamic changes to mediate functional responses of the S protein to binding with the host cell receptor ACE2 and antibodies [1–10]. The cryo-EM and X-ray structures of the SARS-CoV-2 S variants of concern (VOC’s) in various functional states and complexes with antibodies detailed molecular mechanisms, diversity of binding epitopes and allosteric communications between functional regions that underlie binding with different classes of neutralizing antibodies. The cryo-EM structures of the S Omicron BA.1 trimer in the open and closed forms revealed that the dominantly populated conformation is the closed state with all the RBDs buried, leading to ‘conformational masking’ that may prevent antibody binding and neutralization at sites of receptor binding [11–13]. The thermal stability of the S-BA.1, and S-BA.2 protein ectodomains was examined using differential scanning fluorimetry (DSF) assays revealing the reduced stability of the BA.1 RBD and increased stability of the BA.2 RBSD more stable than BA.1 but less stable than the Wu-Hu-1 [14–17]. The cryo-EM structures of the S Omicron protein ectodomain [18,19] showed that the S Omicron BA.1 proteins may preferentially adopt an open 1 RBD-up state predisposed for receptor binding. Moreover, the greater variability of the S-BA.2 trimers seen in the unbound and ACE2-bound forms [18,19] may represent the molecular basis for higher transmission and infection rates of the BA.2 Omicron sublineage compared to BA.1 variant. Cryo-EM structures of the S-BA.2 complexes with human ACE2 revealed a reorganization of the highly antigenic NTD regions resulting in the resistance to antibodies and suggesting that variant-induced modulation of conformational plasticity can underlie the antigenic drift in the Omicron variant landscape and a remarkable increase in the antibody escape pattern [20]. Structural and biophysical studies of the RBD-ACE2 complexes for the Omicron variants revealed that the binding affinity of the Omicron BA.2 with ACE2 is stronger than the affinities of the BA.3 and BA.1 subvariants [21]. Thermal shift assays showed that that the Omicron BA.2-RBD is more stable than that from BA.1, but the observed lower temperature for dissociation of the S-BA.2 trimer indicated that BA.2 is more dynamic than the S-BA.1 trimer [22]. The structural and functional analysis of the S-BA.2 protein trimer revealed three distinct states representing the closed 3-RBD-down conformation, a 1-RBD-up conformation and an RBD-intermediate conformation [23]. The cryo-EM structures of the S trimers for BA.1, BA.2, BA.3, and BA.4/BA.5 subvariants showed that S-BA.1 trimer is stabilized in an open conformation while S-BA.2 exhibiting two conformational states is the least stable among BA.1, BA.2, BA.3 and BA.4/BA.5 variants due to more dynamic and less compact inter-protomer arrangements [24]. The cryo-EM conformations of the BA.2.75 S trimer in the open and closed forms as well as structures of the open BA.2.75 S trimer complexes with ACE2 pointed to the increased structural heterogeneity of S1 regions in the BA.2.75 timer altering the interactions between the RBDs and yet exhibiting a more rigid and compact RBD structure [25]. This study also reported thermal stability of the Omicron variants at neutral pH, showing that the BA.2.75 S-trimer was the most stable, followed by BA.1, BA.2.12.1, BA.5 and BA.2 variants, while BA.2.75 also displayed 4-6-fold increased binding affinity to hACE2 compared with other Omicron variants [25]. Biophysical characterizations of the BA.2.75 variant unveiled the better balance between immune evasion and ACE2 binding, where the binding affinity to ACE2 is increased 9-fold as compared to the BA.2 variant [26–28].

The new recombinant variants such as BA.2.75.2, XBB.1 and XBB.1.5 display substantial growth advantages over previous Omicron variants, suggesting that the immune pressure promotes convergent evolution in which some RBD residues (R346, K356, K444, V445, G446, N450, L452, N460, F486, F490, R493 and S494) observed in at least five independent Omicron sublineages that exhibited a high growth advantage [29,30]. The cryo-EM structures of the XBB.1LJS ectodomain and the XBB.1 S-ACE2 complex revealed two dominant closed states and the RBD one-up state that becomes stabilized in the presence of the ACE2 receptor. These studies showed that despite thermodynamic stability of the XBB.1 S protein, there is considerable plasticity even in the packed closed form of the S trimer [31]. XBB.1.5 is equally immune evasive as XBB.1 but may have growth advantage by virtue of the higher ACE2 binding affinity owing to a single S486P mutation as F486S substitution in XBB.1 [32]. Subsequent functional studies confirmed the growth advantage and the increased transmissibility of the XBB.1.5 lineage due to the preserved strong neutralization resistance and the improved ACE2 binding affinity [33].

The cryo-EM and crystal structures of the S Omicron trimers and complexes with ACE2 and antibodies provided a staggering amount of accurate high-resolution structural information and unveiled diversity of conformational states sampled by Omicron variants. Nonetheless, the details of the intrinsic conformational dynamics and drivers of functional adaptability mechanisms are often hidden or obscured due to rapid interconversion of states and complexity of binding-induced modulation of flexibility seen in the experiments. Hydrogen/deuterium-exchange mass spectrometry (HDX-MS) is a powerful approach for monitoring local protein structural dynamics under native solution conditions whereby by tracking deuterium uptake kinetics for peptide segments throughout a given protein these approaches can inform of dynamics and residue-specific changes in the conformational dynamics induced by mutations or binding. HDX-MS studies uncovered an alternative open S trimer conformation that interconverts slowly with the known prefusion structures and can dynamically expose novel epitopes in the conserved region of the S2 trimer interface, unexpectedly suggesting emergence of hidden cryptic epitopes in a highly conserved region of the protein [34]. Another HDX-MS study identified changes in the S dynamics for VOCs, revealing that Omicron mutations may preferentially induce closed conformations with dynamic core helices in the S2 subunit exploited in the fusion stage and that the NTD acts as a hotspot of conformational divergence driving immune evasion [35]. Furthermore, this study suggested that not only the ACE2 binding promotes opening of the RBDs but can also allosterically enhance dynamics of the S2 core to prime spike prefusion conformations for the transition to the post-fusion form [35]. Comparative HDXMS analysis of the WT, D614G, Alpha, Delta, and Omicron BA.1 S variants tracked evolution of the intrinsic S dynamics by examining the progressively emerging VOC’s [60], showing that Omicron BA.1 can induce greater stabilization of the trimeric stalk interface in the S2 subunit concurrently with the increased NTD and RBD dynamics which may have direct implications for ACE2 binding, and proteolytic processing [36]. HDX-MS studies of the unbound S protein trimer also showed the highest relative exchange in the S2 subunit and helical segments of the stalk regions (central helix CH and heptad repeats HR1 and HR2), while a comparative analysis of the S protein and S-ACE2 complex suggested that ACE2 binding allosterically enhances dynamics at a distal S1/S2 cleavage site and can modulate dynamics of the stalk hinge region, thereby highlighting the role of stalk and proteolysis sites as dynamic allosteric centers in the prefusion state [37]. Another HDX-MS study of S-ACE2 binding confirmed that the ACE2 can induce the enhanced dynamics throughout the entire S protein, with allosteric changes being propagated to the S2 hinge region and to the central helical bundle of the S2 subunit [38]. Overall, these studies suggested that a considerable conformational plasticity can be preserved in both open and stable closed S trimers including more rigid S2 subunit, while Omicron mutations and ACE2 binding can allosterically modulate the S dynamics at distal functional regions.

Structural and functional studies also suggested that Omicron mutations can also impact the dynamics of binding epitopes and mediate formation of transient cryptic binding pockets [39–41]. These binding pockets become exposed or “unmasked” when the S protein undergoes spontaneous conformational changes or interacts with other molecules and antibodies. The cryo-EM structure of the S protein with linoleic acid (LA) complex revealed allosteric binding pocket in the S-RBD that is controlled through opening of a gating helix and can allosterically induce stabilization of the closed S trimer and promote decreased S-ACE2 binding [39,40]. Structural studies discovered a highly conserved cryptic epitope that buried deep inside the trimeric interface in the SD1 region of the S protein and is shared between the Omicron variants of SARS-CoV-2, including WT, BA.1, BQ.1.1, XBB, XBB.1.5 and the XBB.1.16 variants [42]. These highly conserved epitopes enable broad antibody recognition of Omicron variants suggesting S trimer disassembly as mechanism of the antibodies targeting trimeric interface epitopes. These studies further underscored the importance of identifying conserved cryptic epitopes as essential targets for the design of universal vaccines and broadly neutralizing antibodies [42]. The recent structural investigations unveiled another cryptic binding pocket located in the NTD region, also demonstrating that tetrapyrrole products of heme metabolism, biliverdin and bilirubin, can bind to this site with nanomolar affinity and inhibit neutralizing activity of the NTD antibodies recognizing these epitopes [43–45]. By leveraging HDX-MS and antibody engineering approaches, structural studies confirmed a cryptic hinge epitope located in the S2 region between the CH and HR1 segments that play a critical role in the spike conformational changes required for fusion of the viral envelope and target cell membrane [46]. This S2 hinge epitope is partially occluded by the S1 domain and access to this epitope may be controlled through RBD conformational switching.

Computer simulation studies provided important atomistic insights into understanding the dynamics of the SARS-CoV-2 S protein and the effects of Omicron mutations on conformational plasticity of the S protein states and their complexes with diverse binding partners. A number of computational studies employed advanced sampling techniques to gain insight into time scales of conformational changes in the S protein and attempted to characterize conformational landscapes of the S Omicron variants. Adaptive sampling simulations performed on a large scale for the viral proteome captured conformational heterogeneity of the S protein and predicted multiple cryptic epitopes, but the functional relevance and experimental validation of these predicted pockets were lacking [47]. The replica-exchange molecular dynamics (MD) simulations examined conformational landscapes of the full-length S protein trimers, discovering the transition pathways, hidden functional intermediates along open-closed transition pathways and previously unknown cryptic pockets that were consistent with FRET experiments [48]. Computer simulation mapping of the S protein pockets using benzene probes reproduced several experimentally discovered allosteric sites and identified a spectrum of novel cryptic and potentially druggable pockets [49], but this analysis was largely focused on the closed prefusion states of the Wu-Hu-1 S protein and the effects of various Omicron variants and ACE2 binding on evolution of cryptic sites remained unexplored. MD simulations of the unbound or ACE2-bound RBD conformations combined with pocket analysis and druggability prediction identified several promising druggable sites, including one located between the RBD monomers [50]. A network-based adaptation of the reversed allosteric communication approach identified allosteric hotspots and RBD binding pockets in the Omicron variant RBD-ACE2 complexes [51]. Integration of computational and experimental studies enabled discovery and validation of the cryptic allosteric site located between subdomains of the S protein, with several compounds targeting this site showing characteristic binding and anti-virus activities [52,53].

Collectively, the recent structural and computational studies suggested that Omicron mutations and ACE2 binding have significant effect on mediating conformational dynamics changes in the S protein including not only the binding interface regions but also allosterically induced variations and altered plasticity at the remote conserved S2 regions. In contrast to significant accumulation of mutations within the RBD, the S2 fusion subunit has remained highly conserved among variants with only several mutational sites targeted by Omicron variants. There is a steadily increasing interest in identifying novel druggable sites, particularly in the conserved S2 subunit as most recently developed broad-spectrum fusion inhibitors and candidate vaccines can target the functional regions in the S2 subunit [54,55]. These investigations also underscored the enormous diversity of potential cryptic pockets with some of them being present only transiently while others showing conservation across different S protein states.

Despite recent advances, our current structural and functional knowledge of the cryptic binding pockets and potential allosteric sites in the S protein is incomplete and lacks deep analysis of evolution, functional mechanisms and validity of the predicted cryptic sites. In addition, there is no clear understanding of the effects exerted by different Omicron variants, conformational states and ACE2 binding on evolution and redistribution of the cryptic pockets in the S protein. Finally, it remains unclear whether currently validated cryptic pockets are preserved in the S protein dynamic landscapes through evolutionary changes of Omicron variants and how functional roles of these binding sites is related to allosteric mechanisms and functional transitions of the S protein. To investigate and explore these outstanding issues, in the current study we perform a systematic comparative analysis of the conformational dynamics, allostery and cryptic binding pockets in the RBD-ACE2 complexes trimers and S trimer complexes with the ACE2 receptor. We first perform multiple microsecond MD simulations and Markov state model (MSM) analysis of the SARS-CoV-2 S RBD-ACE2 complexes for the Omicron BA.2, BA.2.75 and XBB.1 variants. Using a comparative MSM analysis we discover variant-specific changes of conformational mobility and confirm stability of the BA.2.75 RBD and increased mobility of the XBB.1 RBD. Using conformational ensembles of the SARS-CoV-2 Omicro S trimers, we conducted a systematic binding pocket screening and analysis of functional cryptic pockets in the BA.2, BA.2.75 and XBB.1 complexes with ACE2. The results of this study connect insights from conformational dynamics analysis, comparative mutational scanning of the S-ACE2 binding and the inter-protomer interactions with the evolution of cryptic binding sites. We show that our approach can capture and reproduce all experimentally known allosteric sites and discover networks of conserved cryptic pockets preserved in different conformational states of the Omicron variants and in the S-ACE2 complexes. The results of our study suggest that despite general rigidity of the S2 regions in comparison with more dynamic S1 subunit, there is still an appreciable level of conformational adaptability in the S2, resulting in significant number of dynamic cryptic pockets. The results detailed how mutational and conformational changes in the BA.2 and BA.2.75 spike trimers can modulate the distributions and mediate networks of inter-connected conserved and also variant-specific druggable allosteric pockets. The results of this study can be important for understanding mechanisms underlying functional roles of druggable cryptic pockets that can be used for both site-specific and allostery-inspired therapeutic intervention strategy targeting distinct conformational states of the SARS-CoV-2 Omicron variants.

## Materials and Methods

### All-Atom Molecular Dynamics Simulations

The crystal structures of the BA.2 RBD-ACE2 (pdb id 7XB0) [21], and BA.2.75 RBD-ACE2 complexes (pdb id 8ASY) [56] and the atomic coordinates for the structures of XBB.1LJS RBD bound to ACE2 (pdb id 8IOV) [31] as well as the cryo-EM structures of the BA.2 S trimer with two human ACE2 bound (pdb id 7XO7) [22], BA.2 S trimer with two three ACE2 bound (pdb id 7XO8) [22], BA.2.75 S trimer with one human ACE2 bound (pdb id 7YR2) [25] and XBB.1 S trimer with one human ACE2 bound [31] (Figure 1) were obtained from the Protein Data Bank [57]. During structure preparation stage, protein residues in the crystal structures were inspected for missing residues and protons. Hydrogen atoms and missing residues were initially added and assigned according to the WHATIF program web interface [58]. The missing loops in the studied structures of the SARS-CoV-2 S protein were reconstructed and optimized using template-based loop prediction approach ArchPRED [59]. The side chain rotamers were refined and optimized by SCWRL4 tool [60]. The protonation states for all the titratable residues of the ACE2 and RBD proteins were predicted at pH 7.0 using Propka 3.1 software and web server [61,62]. The protein structures were then optimized using atomic-level energy minimization with composite physics and knowledge-based force fields implemented in the 3Drefine method [63].

**Figure 1.**
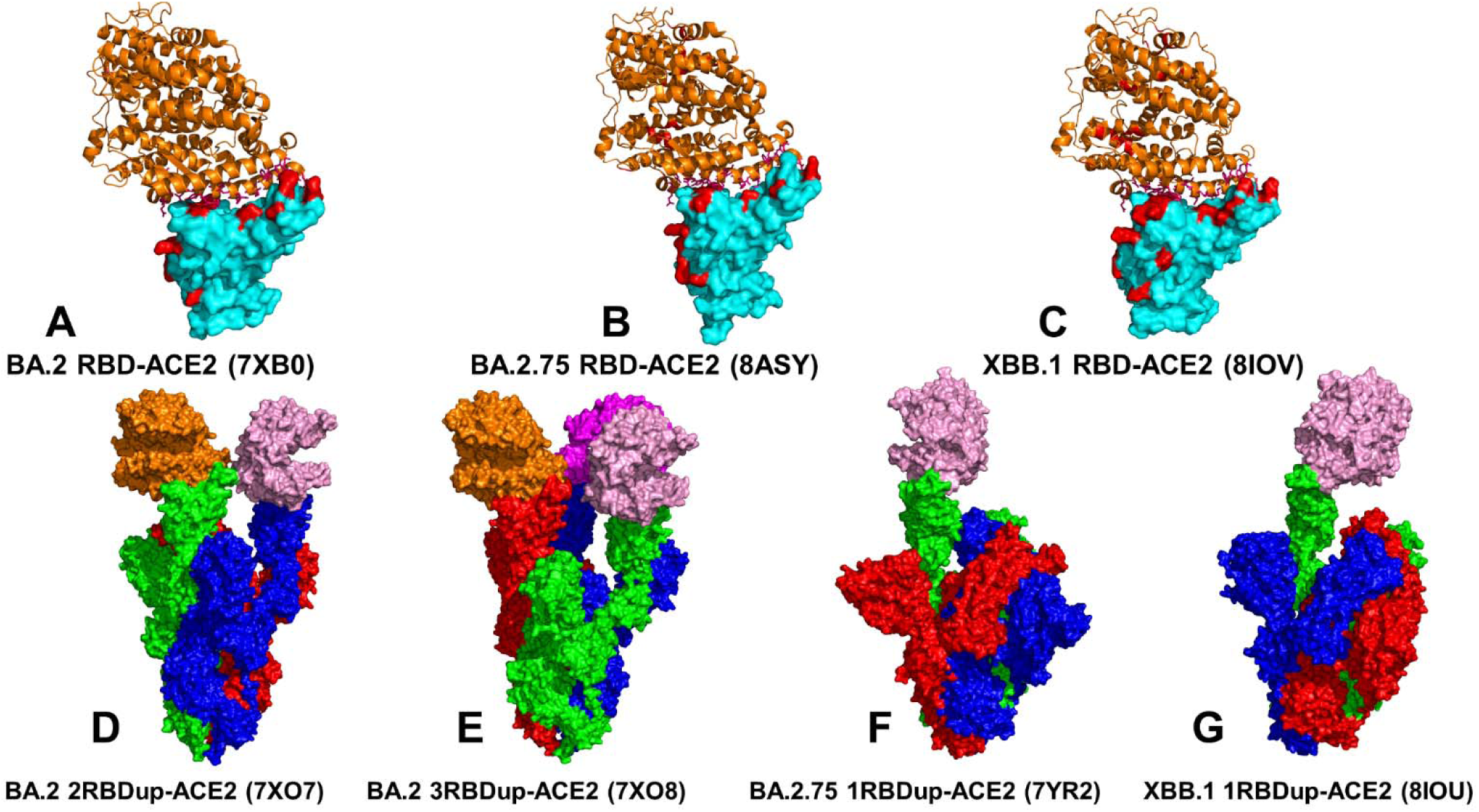
Structural organization of the SARS-CoV-2-RBD Omicron BA.2, BA.2.75, and XBB.1.complexes with human ACE2 enzyme and Omicron S trimer complexes with ACE2. (A) The crystal structure of the Omicron RBD BA.2-ACE2 complex (pdb id 7XB0). The RBD is shown in cyan-colored surface and the bound ACE2 enzyme is in orange ribbons. (B) The crystal structure of the Omicron RBD BA.2.75-ACE2 complex (pdb id 8ASY). The RBD is shown in cyan-colored surface and the bound ACE2 enzyme is in orange ribbons. (C) The cryo-EM structure of the Omicron RBD XBB.1-ACE2 complex. The RBD is shown in cyan-colored surface and the bound ACE2 enzyme is in orange ribbons. The Omicron RBD mutational sites are shown in red-colored surface. (D) The cryo-EM structure of the BA.2 S trimer with two human ACE2 bound (pdb id 7XO7). The protomers are shown in green, red and blue surface. The ACE2 molecules are shown in pink and orange surface. (E) The cryo-EM structure of the BA.2 S trimer with three human ACE2 bound (pdb id 7XO8). The protomers are shown in green, red and blue surface. The ACE2 molecules are shown in pink, orange and magenta surface respectively. (F) The cryo-EM structure of the BA.2.75 S trimer with one human ACE2 bound (pdb id 7YR2). The protomers are in green, red and blue surface. The ACE2 molecule is in pink surface. (G) The cryo-EM structure of the XBB.1 trimer with one human ACE2 bound (pdb id 8IOU). The protomers are in green, red and blue surface. The ACE2 molecule is in pink surface.

We employed CHARMM36M force field [64] with the TIP3P water model [65] to conduct microsecond all-atom MD simulations for structures of the BA.2, BA.2.2.75 and XBB.1 RBD-ACE2 complexes. The protein systems were solvated in 130 Å × 85 Å × 75 Å water boxes. In each system, sodium and chloride ions were added to maintain an ionic strength of 0.1 M. After energy minimization, the systems were first heated up from 100 to 300 K with a temperature increment of 20 K per 50 picoseconds (ps). Consequently, the systems were subjected to 1.5 nanoseconds (ns) isothermal−isobaric (NPT) equilibrations at 300 K (equilibrium run), followed by 1 microsecond (µs) canonical (NVT) simulations (production run) at 300 K. Snapshots of the production run were saved every 100 ps. In all simulations, the SHAKE constraint was used to constrain bonds associated with hydrogen atoms in the solvent molecules and the proteins [66]. The nonbonding interactions within 10 Å were calculated explicitly. The Lennard-Jones interactions were smoothed out to zero at 12 Å. The long-range electrostatic interactions were calculated using the particle mesh Ewald method [67] using a fourth order (cubic) interpolation and a cutoff of 10.0 Å. The simulations were conducted using OpenMM (version 7.6.0) software package [68]. For each system, MD simulations were conducted three times in parallel to obtain comprehensive sampling. Each individual simulation has 10,000 frames. For BA.2 RBD-ACE2, BA.2.75 RBD-ACE2 and XBB.1 RBD-ACE2 systems, MD simulations were conducted three times in parallel for each system to obtain comprehensive sampling. Each individual simulation of the studied Omicro n RBD-ACE2 complexes stored 10,000 frames, thus we collected 30,000 frames for each of the complexes.

While the production stage of MD simulations is often performed under the NPT ensemble to enable more accurate comparison with the experimental data, the NPT ensemble often needs significantly longer time to reach convergence because both temperature and volume fluctuate during the simulations. In this study we opted for an alternative and widely used approach in which the equilibrium simulations are carried out in the NPT ensemble to allow the total energy and volume of the system fluctuate to reach conditions suitable for the target pressure. In this case, the production simulations could be carried out in the NVT ensemble using the equilibrium volume sampled in the equilibration NPT simulations. This alternative protocol aims to take advantage of the NVT ensemble for better convergence than NPT ensemble given the same length of simulations. Although the binding association constants for complexes are often measured under constant (atmospheric) pressure and temperature (NPT ensemble), corresponding to Gibbs free energy, the respective experimentally measured macroscopic parameters obtained in the NVT ensemble, corresponding to Helmholtz free energy, are typically very similar.

A detailed comparison of MD simulations using different force fields and ensembles (including NAMD with the CHARMM36 force field and NVT for the production phase) produced models and equilibrium ensembles that are in similar agreement with experimental results [69]. Moreover, the agreement between MD simulations and experimental observables may not necessarily constitute a validation of the conformational ensemble produced by MD simulations as diverse conformational ensembles may produce averages that are consistent with the experiment [69]. Importantly, as the main objective of our study is the examination of the effects of different mutations on the SARS-CoV-2 S protein complexes, the employment of either NPT or NVT ensemble for the production simulations is not expected to produce appreciable differences and lead to different conclusions. We used MDTraj Python library [70] for the analysis, manipulation, and visualization of MD simulation trajectories. MDtraj provides a wide range of functionalities to extract structural and dynamic information from MD trajectories, including a fast RMSD procedure [71] that executes QCP algorithm [72] three times faster than the original implementation. The RMSD calculations were also performed using MDAnalysis Python toolkit (www.mdanalysis.org) utilizing the fast QCP algorithm and the optimal rotation matrix for superposition of simulation frames [73].

### Markov State Model Analysis of the Omicron RBD-ACE2 Complexes

The time-structure Independent Components Analysis (tICA) method identifies the slowest degrees of freedom and can preserve the kinetic information present in the MD trajectories by maximizing the auto-correlation function [74–77]. In contrast to principal component analysis (PCA), which finds coordinates of maximal variance, tICA finds coordinates of maximal autocorrelation at the given lag time. Therefore, the tICA approach is useful to find the *slow* components in a dataset and a robust method to transform MD simulation information before clustering data for the construction of a Markov model. Using a time-series of molecular coordinates provided by an *n*-dimensional MD trajectory **x**(*t*) = (*x*_1_(*t*), …, *x_n_*(*t*))*^T^*∈ □*^n^* with Cartesian coordinates (*x*_1_, …, *x_n_*) tICA can reduce the dimensionality of the trajectories and determine the slowest independent collective degrees of freedom onto which the projections of the initial dataset have the largest time-autocorrelation. The tICA approach identifies the slowest degrees of freedom by solving the generalized eigenvalue problem :

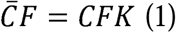

where *K* = *diag*(*k*_1_,…,*k_n_*) and *F* = (*f*_1_,…,*f_n_*) are the eigenvalue and eigenvector matrices, respectively; C and C are the covariance matrix and the time-lagged covariance matrix of the coordinate vector defined as follows :

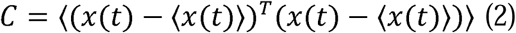

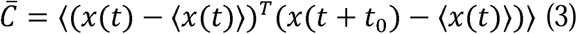

To obtain a symmetric time-lagged covariance matrix, 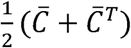 is calculated. The latter step assumes the time reversibility of the process, which is satisfied in MD simulations. The featurization and dimensionality reduction were performed using the PyEMMA package [78].

We employed Stochastic Markov state models (MSMs) [79–83] to characterize conformational landscapes of the Omicron RBD-ACE2 complexes and describe the transitions between functional protein states. In MSM, protein dynamics is modeled as a kinetic process consisting of a series of Markovian transitions between different conformational states at discrete time intervals. A specific time interval, referred to as lag time, needs to be determined to construct transition matrix. First, k-means clustering method is conducted on projected low-dimensional space and each simulation frame is assigned to a microstate. The transition counting is constructed based on a specific time interval lag time *τ*. Macrostates are kinetically clustered based on the Perron-cluster cluster analysis (PCCA++) [84] and considered to be kinetically separate equilibrium states. The transition matrix and transition probability were calculated to quantify the transition dynamics among macrostates. The corresponding transition probability from state *i* to state *j* is calculated as:

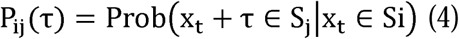

A proper lag time is required for MSM to be Markovian. The value of the lag time and the number of macrostates are selected based on the result of estimated relaxation timescale [85]. The implied timescales can be calculated using the eigenvalues (*λ*_i_) in the transition matrix as

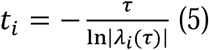

The number of protein metastable states associated with these slow relaxation timescales can be inferred based on the convergence of implied relaxation time scale. These metastable states effectively discretize the conformational landscape. The Markov state model building was conducted using PyEMMA package (v2.5.12) [78]. Based on the transition matrix we obtain implied timescales for transitioning between various regions of phase space and use this information determines the number of metastable states.

### Coarse-Grained Dynamics Simulations of the Omicron S trimer complexes with ACE2

Coarse-grained Brownian dynamics (CG-BD) simulations have been conducted for the cryo-EM structures of the BA.2 S trimer with two human ACE2 bound (pdb id 7XO7) [22], BA.2 S trimer with two three ACE2 bound (pdb id 7XO8) [22], BA.2.75 S trimer with one human ACE2 bound (pdb id 7YR2) [25] and XBB.1 S trimer with one human ACE2 bound [31] (Figure 1). We employed ProPHet (Probing Protein Heterogeneity) approach and program [86–88]. BD simulations employed a high resolution CG protein representation of the SARS-CoV-2 S Omicron trimer structures that can distinguish different residues. In this model, each amino acid is represented by one pseudo-atom at the Cα position, and two pseudo-atoms for large residues. The interactions between the pseudo-atoms are treated according to the standard elastic network model (ENM) in which the pseudo-atoms within the cut-off parameter, *R*_c_ = 9 Å are joined by Gaussian springs with the identical spring constants of _γ_ = 0.42 N m^−1^ (0.6 kcal mol^−1^ Å^−2^. The simulations use an implicit solvent representation via the diffusion and random displacement terms and hydrodynamic interactions through the diffusion tensor using the Ermak-McCammon equation of motions and hydrodynamic interactions [89,90]. The stability of the SARS-CoV-2 S Omicron trimers was monitored in multiple simulations with different time steps and running times. We adopted Δ*t* = 5 fs as a time step for simulations and performed 100 independent BD simulations for each system using 500,000 BD steps at a temperature of 300 K. The CG-BD conformational ensembles were also subjected to all-atom reconstruction using PULCHRA method [91] and CG2AA tool [92] to produce atomistic models of simulation trajectories.

### Mutational Scanning Analysis of the RBD-ACE2 Binding Interactions and Inter-Protomer Trimer Interactions

Mutational scanning analysis of the binding epitope residues for the Omicron RBD-ACE2 and S trimer-ACE2 complexes was conducted using PoPMuSiC approach [93–95] that is based on statistical potentials which include contributions from the pairwise inter-residue distances, backbone torsion angles and solvent accessibilities, and considers the effect of the mutation on the strength of the interactions at the interface and on the overall stability of the complex. The binding free energy of protein-protein complex can be expressed as the difference in the folding free energy of the complex and folding free energies of the two protein binding partners:

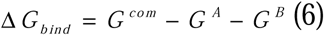

The change of the binding energy due to a mutation was calculated then as the following:

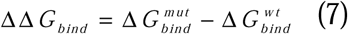

In this approach, the binding interface residues are defined based on the condition that the difference between a residue’s solvent accessibility in the complex and apo-protein is at least 5%. The solvent accessibility is defined as the ratio of the solvent-accessible surface in the considered structure relative to an extended tripeptide Gly-X-Gly. Each binding epitope residue was systematically mutated using all substitutions and corresponding protein stability and binding free energy changes were computed. We computed the ensemble-averaged binding free energy changes using equilibrium samples from simulation trajectories. The binding free energy changes were computed by averaging the results over 1,000 equilibrium samples that were evenly distributed along each of the three independent MD simulations for each of the studied systems. Hence, a total of 3,000 protein snapshots were employed in the computations of the mutation-induced binding free energies for each of the studied system.

In addition to mutational scanning of the RBD-ACE2 binding interfaces, the S protein residues involved in the inter-protomer contacts in the S trimer structures were also systematically mutated to evaluate the protein stability differences between BA.2, BA.2.75 and XBB.1 Omicron variants. If a free energy change between a mutant and the wild type (WT) proteins ΔΔG= ΔG (MT)-ΔG (WT) > 0, the mutation is destabilizing, while when ΔΔG <0 the respective mutation is stabilizing. We computed the average ΔΔG values using 1,000 samples from the CG-BD equilibrium ensembles.

### Machine Learning Detection of Cryptic Pockets

We used two different complementary approaches for identification of the cryptic binding pockets in the conformational ensembles of the Omicron RBD-ACE2 and S timer-ACE2 complex structures. A template-free P2Rank approach is among the most efficient and fast available ML tools for prediction of ligand binding sites that combines sequence and structural data to rank potential binding sites based on their likelihood of binding a specific ligand [96,97]. P2Rank uses support vector machine (SVM), random forests (RF), and artificial neural networks (ANNs) to learn the ligandability of a local chemical environment that is centered on points placed on the protein’s solvent-accessible surface [96,97]. P2Rank v2.4 with default parameters was deployed to identify pockets across all of the representative states from our simulations. By combining eXtreme gradient boosting (XGBoost) and graph convolutional neural networks (GCNNs) a robust approach for allosteric site identification and Prediction of Allosteric Sites Server (PASSer) was developed [98–100]. We also employed the PASSer Learning to Rank (LTR) model that is capable of ranking pockets in order of their likelihood to be allosteric sites [100]. Using P2Rank [96,97] and PASSer LTR [100] approaches, we identified binding pockets in the conformational ensembles and computed P2Rank-predicted residue pocket probability. The reported top binding pockets for each protein structure correspond to top-ranked consensus P2Rank/LTR predicted sites.

## Results

### Atomistic MD Simulations and Markov State Model Analysis of the Omicron BA.2, BA.2.75 and XBB.1 RBD-ACE2 Complexes

We first conducted analysis of the conformational dynamics and distribution of states in the Omicron BA.2, BA.2.75 and XBB.1 RBD-ACE2 complexes (Figure 1, Supplementary Materials, Figure S1) using microsecond atomistic MD simulations combined with subsequent dimensionality reduction and MSM analysis of the conformational landscapes. Here, we considerably expanded our recent simulations of the Omicron RBD-ACE2 complexes [101] by performing a systematic comparative analysis using multiple microsecond MD simulations based on the experimental structures for all studied Omicron RBD-ACE2 complexes (Figure 1) which included the recently released structure of the XBB.1 RBD-ACE2 complex [31]. By expanding to the microsecond time scale, a combination of MD simulations and MSM analysis provided evidence that despite highly similar RBD-ACE2 structures, the distributions of states for different Omicron variant complexes are considerably different. The RMSF profiles confirmed stability of the β-sheet RBD core region (residues 350-360, 375-380, 394-403) (Figure 2A), while revealing characteristic RMSF peaks corresponding to the flexible RBD regions (residues 360-373 and residues 380-396) (Figure 2A). In general, the RMSF profiles showed that thermal displacements observe in atomistic simulations of the BA.2, BA.2.75 and XBB.1/XBB.1.5 RBDs are similar, but also highlighted greater fluctuations of the flexible RBD regions in the XBB.1 RBD-ACE2 complex (Figure 2A). The receptor-binding motif (RBM) is the main functional motif in RBD and is composed of the RBD residues 437-508 that form the binding interface between the RBD and ACE2 (Supplementary Materials, Figure S2). The recognition loop residues (residues 470-491) of the RBM form a dense interaction network with the ACE2 receptor that determines the RBD-ACE2 binding affinity. We observed some differences in the flexible RBD regions (residues 355-375, 380-400, 480-490), also pointing to the greater local mobility of the RBM residues in the XBB.1 RBD (Figure 2A). While the flexible RBD loop (residues 440-452) contains several convergent mutational sites present in the XBB.1 variant (K444, V445, G446, N450, L452), we observed no appreciable differences in its mobility between Omicron variants. It may be argued that moderately increased RBM mobility in XBB.1 may result from F486S and F490S substitutions that could weaken the RBM-ACE2 interfacial contacts and allow for greater flexibility in this region.

**Figure 2.**
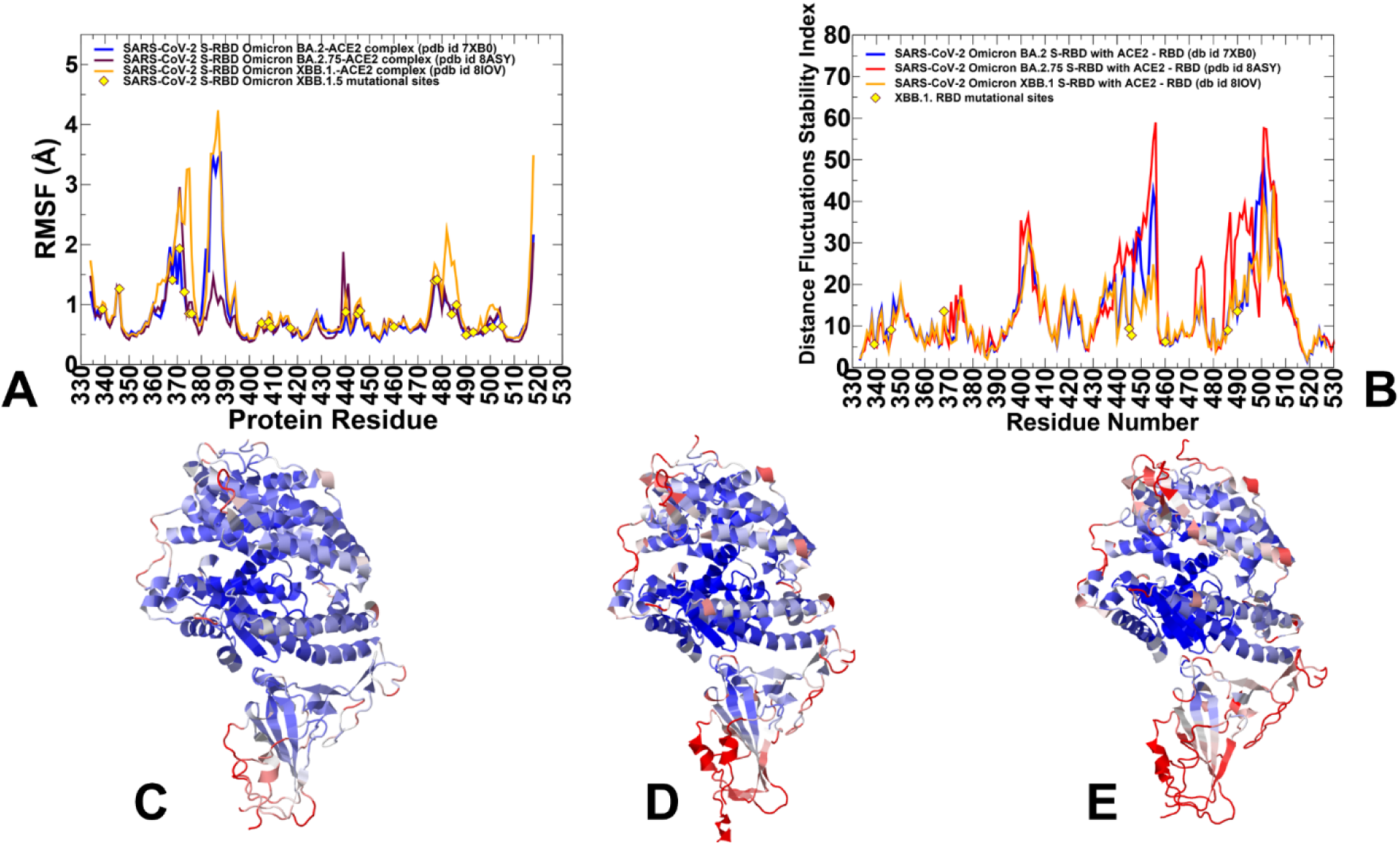
Conformational dynamics profiles of the Omicron RBD BA.2, BA.2.75 and XBB.1 RBD complexes with ACE2 obtained by averaging results from three independent all-atom 1µs MD simulations (A) The RMSF profiles for the RBD residues obtained from MD simulations of the BA.2 RBD-ACE2 complex, pdb id 7XB0 (in blue lines), BA.2.75 RBD-ACE2 complex, pdb id 8ASY (in maroon lines), and XBB.1. RBD-ACE2 complex, pdb id 8IOV (in orange lines). The positions of Omicron XBB.1 RBD mutational sites are highlighted in yellow-colored filled diamonds. (B) The distance fluctuations stability index profiles of the RBD residues obtained by averaging results from 3 independent all-atom 1µs MD simulations of the Omicron RBD-ACE2 complexes. The stability index profiles are shown for Omicron RBD BA.2 (in blue lines) BA.2.75 (in maroon lines), XBB.1 (in orange lines). The positions of the Omicron XBB.1 RBD mutational sites are highlighted in yellow-colored filled diamonds. Structural maps of the conformational profiles obtained from MD simulations of the Omicron RBD-ACE2 complexes. Conformational mobility maps are shown for the Omicron RBD BA.2-ACE2 complex (C), the Omicron RBD BA.2.75-ACE2 complex (D), and the Omicron RBD XBB.1-ACE2 complex (E). The structures are shown in ribbons with the rigidity-flexibility sliding scale colored from blue (most rigid) to red (most flexible).

The RBM tip is structurally ordered (“hook-like” fold) in both BA.2 and BA.2.75 conformational samples throughout the MD trajectory, while F486S mutation in the XBB.1 RBD-ACE2 complex may induce partly disordered configurations of the RBM tip (Figure 2A). The RMSF analysis of the ACE2 residues showed similar profiles across variants (Supplementary Materials, Figure S3). The stable ACE2 residues are in the core and the binding interface positions. The key ACE2 α-helix (residues 24-31) and a β-sheet (residue 350-356) show small RMSF values in all complexes (Supplementary Materials, Figure S3).

Another useful metric of the local conformational dynamics patterns is the distance fluctuation stability index that evaluates the extent of fluctuations in the mean distance between each residue and all other protein residues [86–88, 101]. The high values of distance fluctuation stability indexes are associated with structurally rigid residues as they display small fluctuations in their distances to all other residues, while small values of this index are characteristic of flexible sites [86–88]. The distributions predicted the local maxima for structurally stable regions in the RBD core (residues 400-420, 450-460) and to the RBD binding interface region (residues 484-505) (Figure 2B). The differences in the stability index profiles are mostly in the RBM region (residues 470-491) showing the greater stability for the BA.2.75 RBD, while smaller indexes are seen in the XBB.1 RBD profile in support of the greater mobility of this region (Figure 2B).

The conformational dynamics profiles projected onto the RBD-ACE2 structures showed a similar and strong stabilization of the RBD core, RBD binding interface, and the interfacial helices on ACE2 (Figure 2C-E). Although the conformational dynamics profiles revealed certain important differences between Omicron variants pointing to more flexible XBB.1 RBD and confirming stability of the BA.2.75 RBD, these metrics are poorly suited to distinguish salient features of the underlying landscapes and provide a quantitative characterization of the dynamic state equilibrium. To address these issues and identify the distribution of states we used tICA dimensionality reduction method by projecting the results of MD simulations onto low dimensional space. The low-dimensionality projection of the MD ensemble obtained from microsecond trajectories highlighted broader densities for BA.2 (Figure 3A,B) and XBB.1 RBDs (Figure 3D) while revealing narrow high-density region sampled in the BA.2.75 complex (Figure 3C) indicative of greater structural rigidity of the BA.2.75 RBD. Notably, the densities obtained from different simulations indicated wider dispersion of samples between the three different trajectories, while for BA.2.75 RBD-ACE2 complex all simulations converged to the crystallographic conformation that domines the thermodynamic equilibrium for this variant (Figure 3C). Hence, the low-dimensional projection analysis emphasized the key differences in the conformational dynamics of the Omicron variants, revealing the existence of multiple functional RBD conformations in the BA.2 and XBB.1 RBD-ACE2 complexes and supporting the increased RBD plasticity induced by these two variants. In contrast, the striking convergence of three independent microsecond simulations of the BA.2.75 RBD-ACE2 complex confirmed structural stabilization of the RBD for this Omicron variant. These findings corroborate with the experimental data showing a more rigid and compact BA.2.75 RBD structure as compared to other Omicron variants [25].

**Figure 3.**
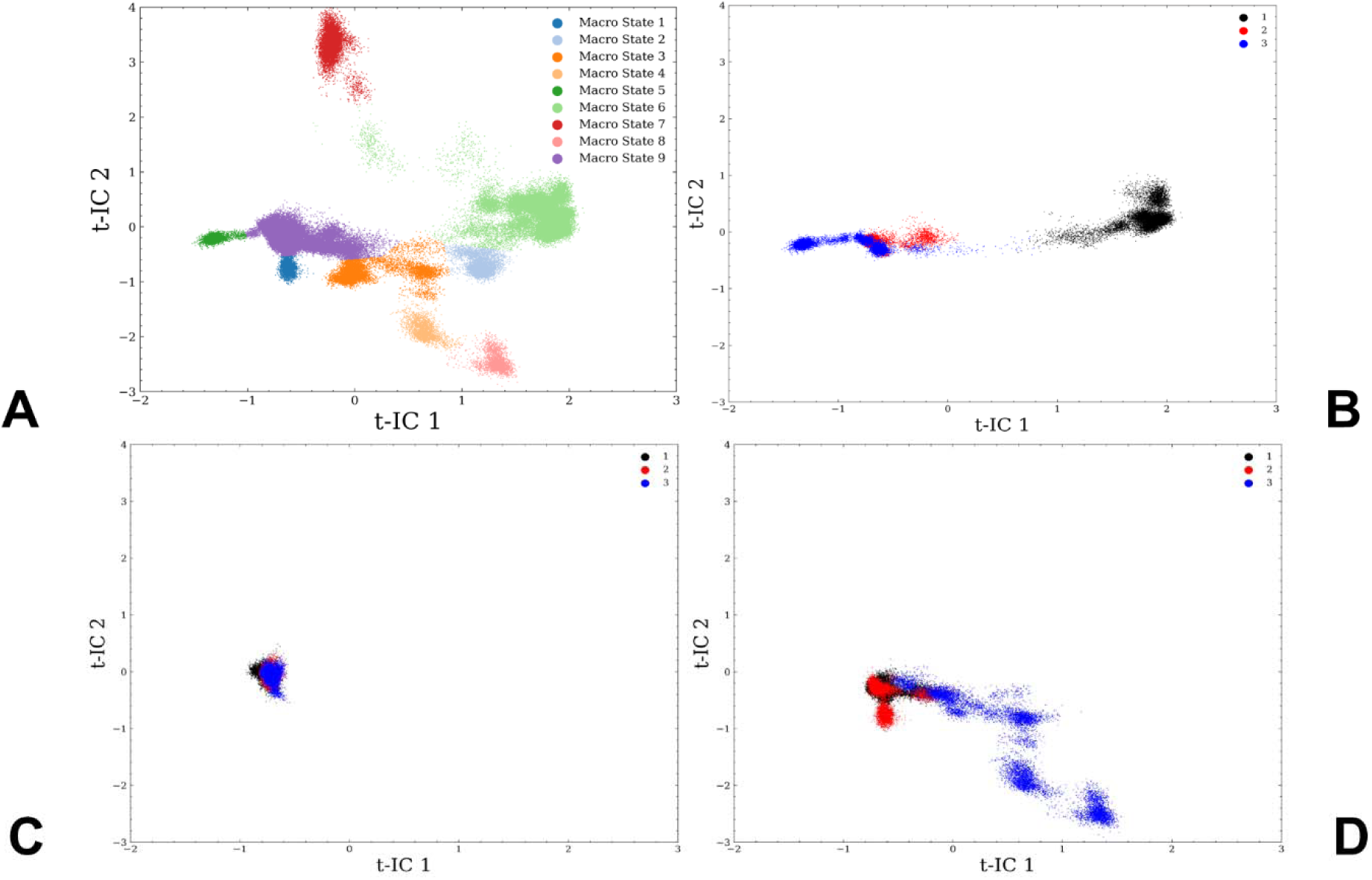
The t-ICA low-dimensional projection of the conformational space sampled in three independent microsecond MD simulations for the Omicron BA.2, BA.2.75 and XBB.1 RBD-ACE2 complexes. (A) The low-dimensional density representations with 2 components (t-IC) are shown for the identified macrostates 1-9. (B) The t-ICA low-dimensional projection of the conformational space for Omicron BA.2 RBD-ACE2 complex. (C) The t-ICA low-dimensional projection of the conformational space for Omicron BA.2.75 RBD-ACE2 complex. (D) The t-ICA low-dimensional projection of the conformational space for Omicron XBB.1 RBD-ACE2 complex. The densities are shown in black dots for MD trajectory 1, in red dots for MD trajectory 2 and in blue dots for MD trajectory 3.

Using low-dimensional characterization of the conformational ensembles, we proceeded with the MSM analysis and partitioning of the MSM macrostates (Figure 4). A total of nine macrostates emerged from the MSM analysis and the stationary distribution of states were evaluated for BA.2 (Figure 4A), BA.2.75 (Figure 4B) and XBB.1 RBD-ACE2 complexes (Figure 4C). For the BA.2 RBD-ACE2 ensemble, macrostates 2, 3, 5, 6 and 9 contribute to the distribution with the macrostate 9 dominating ∼ the conformational population (Figure 4A). This microstate corresponds to the crystallographic conformation of the complex, showing that other RBD conformations (macrostates 5, 6) showing variations in the flexible RBD loops could appreciably contribute to the equilibrium and binding with ACE2. A single peak distribution in the BA.2.75 RBD-ACE2 complex confirmed that this ensemble is overwhelmingly dominated by the native conformation (Figure 4B) suggestive of markedly rigidified RBD structure for this variant. In contrast, we detected a broader distribution of macrostates for the XBB.1 RBD-ACE2 complex (Figure 4C) where macrostates 1, 3, 4, 8 and 9 contribute to the equilibrium population. Nonetheless, the distribution is still dominated by the crystallographic state (macrostate 9) contributing 62%, and yet there are several additional macrostates present in the equilibrium which supported the notion of the greater XBB.1 RBD plasticity (Figure 4C).

**Figure 4.**
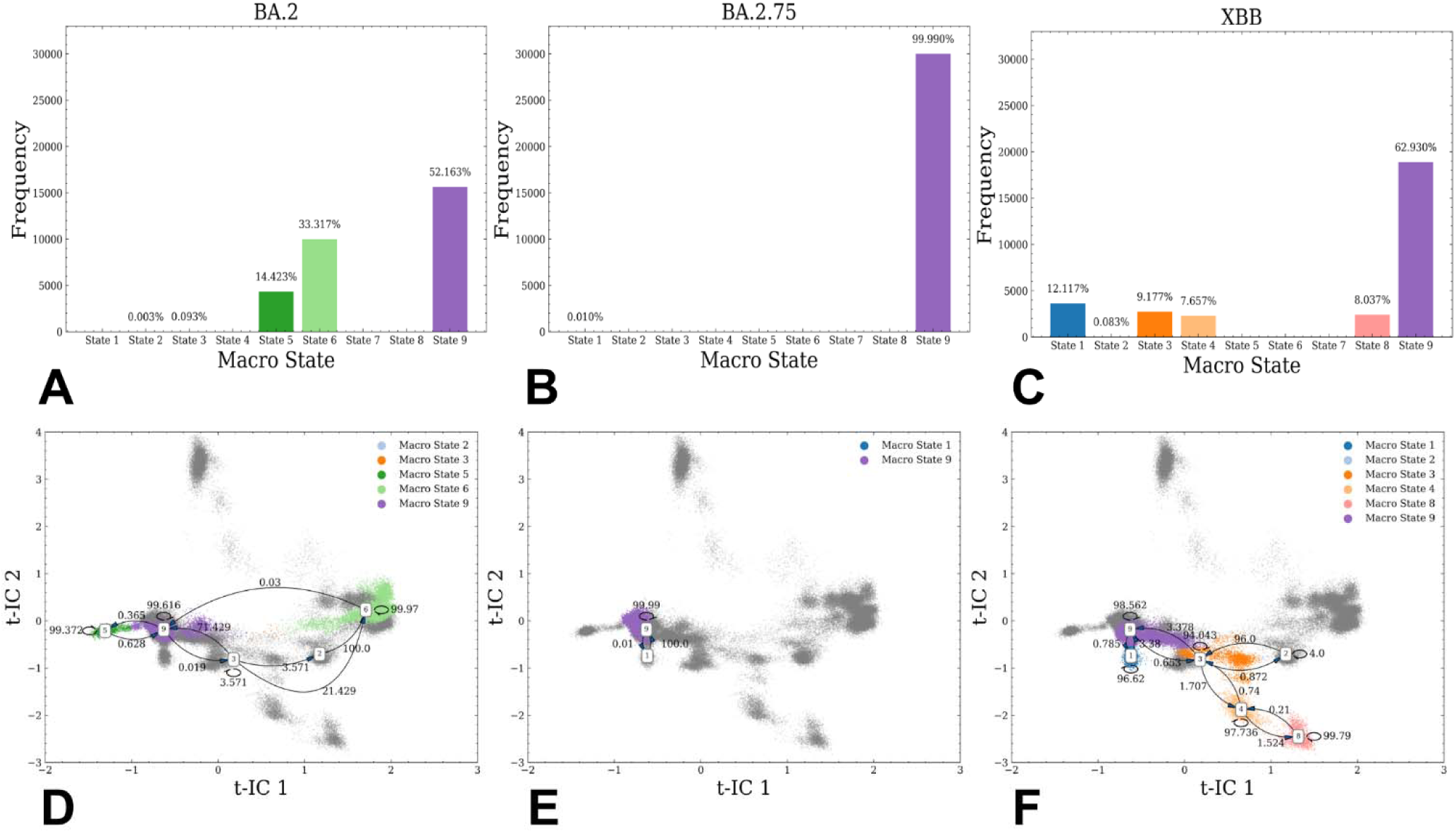
MSM analysis of the conformational landscape for the Omicron BA.2, BA.2.75 and XBB.1 RBD-ACE2 complexes. The frequency distribution of occupying different macrostates in BA.2 RBD-ACE2 (A), BA.2.75 RBD-ACE2 (B) and XBB.1 RBD-ACE2 (C). The transition probability maps among different macrostates with the 5 ns lag time are shown for BA.2 RBD-ACE2 (D), BA.2.75 RBD-ACE2 (E) and XBB.1 RBD-ACE2 (F).

These results revealed important differences in the conformational landscapes and distribution of functional conformations in the Omicron variants, still showing that the topology of the RBD fold and RBD-ACE2 binding interface are fully preserved across all variants. As a result, functional and binding differences between BA.2, BA.2.75 and XBB.1 variants can be determined through moderate variations in conformational flexibility that enable modulation and balancing of multiple fitness tradeoffs between immune evasion, ACE2 affinity and conformational adaptability which may potentially be an important driver behind evolution of Omicron variants.

The transition probabilities were determined among different macrostates for all systems, with the 5 ns lag time. The high percentage of self-conserved probability shows the stability of macrostates. The results showed significant and informative differences in the distributions and transitional probability maps between these structurally similar RBD-ACE2 complexes. The transitional probability map for the BA.2 RBD complex revealed that the macrostates 9 and 6 can interconvert representing two major dense areas (Figure 4D) indicating that the equilibrium population for this variant is determined by these RBD conformations. Structural inspection of the microstate 9 and 6 (Supplementary Materials, Figure S4) showed significant differences in the flexible regions, particularly in the RBM tip region that adopted an alternative orientation as compared to the crystallographic conformation. Hence, BA.2 RBD maintains a considerable plasticity in the complex with ACE2 suggesting that entropic component may contribute to the binding affinity of this complex.

Remarkably, the transition map for the BA.2.75 RBD-ACE2 complex showed only a single conformation that is virtually identical to the crystallographic state (Figure 4E). In sharp contrast, transition maps for XBB.1 (Figure 4F) revealed that dominant macrostate 9 can interconvert with macrostates 1 and 3 where microstate 3 is connected with other identified macrostates of the ensemble. Structural analysis of the macrostates showed a considerable remodeling of the flexible RBD region (residues 460-490) including different orientations of the RBM tip (Supplementary Materials, Figure S4). The MSM analysis also revealed that dominant macrostate 9 featured well-ordered and stable “hook-like” conformation of the RBM tip. This state becomes less favorable in XBB.1 (F486S) while the contribution of other macrostates with a partly disordered conformation of the RBM tip and more dynamic RBD loops increased.

XBB.1 subvariant is a descendant of BA.2 and recombinant of BA.2.10.1 and BA.2.75 sublineages, also featuring a group of specific RBD mutations (G339H, R346T, L368I, V445P, G446S, N460K, F486S, F490S and reversed R493Q) (Table 1). Many of these RBD mutations are known for their immune evasion functions, including R346T, G446S and F486S. Based on the structural comparison of the macrostates, it becomes apparent that a combination of F486S and F490S mutations in XBB.1 may induce the increased RBD mobility in regions important for antibody binding and thereby enhance immune evasion potential of this Omicron variant. The MSM analysis suggested that BA.2 and XBB.1 RBDs may exhibit a similar degree of conformational plasticity in the complex, while BA.2.75 RBD is characterized by pronounced structural rigidity featuring a single dominant microstate.

**Table 1.** Mutational landscape of the Omicron BA.2, BA.2.75 and XBB.1 Variants.

| **Omicron Variant** | **Mutational landscape** |
| --- | --- |
| BA.1 | A67, T95I, G339D, S371L, S373P, S375F, K417N, N440K, G446S, S477N, T478K, E484A, Q493R, G496S, Q498R, N501Y, Y505H, T547K, D614G, H655Y, N679K, P681H, N764K, D796Y, N856K, Q954H, N969K, L981F |
| BA.2 | T19I, G142D, V213G, G339D, S371F, S373P, S375F, T376A, D405N, R408S, K417N, N440K, S477N, T478K, E484A, Q493R, Q498R, N501Y, Y505H, D614G, H655Y, N679K, P681H, N764K, D796Y, Q954H, N969K |
| BA.2.75 | T19I, G142D, K147E, W152R, F157L, I210V, V213G, G257S, G339H, S371F, S373P, S375F, T376A, D405N, R408S, K417N, N440K, G446N, N460K, S477N, T478K, E484A, Q498R, N501Y, Y505H, D614G, H655Y, N679K, P681H, N764K, D796Y, Q954H, N969K |
| XBB.1 | T19I, V83A, G142D, Del144, H146Q, Q183E, V213E, G252V, G339H, R346T, L368I, S371F, S373P, S375F, T376A, D405N, R408S, K417N, N440K, V445P, G446S N460K, S477N, T478K, E484A, F486S, F490S, R493Q reversal, Q498R, N501Y, Y505H, D614G, H655Y, N679K, P681H, N764K, D796Y, Q954H, N969K |

### Coarse-Grained Simulations of the Omicron BA.2, BA.2.75 and XBB.1 S Trimer-ACE2 Complexes

Coarse-grained Brownian dynamics (CG-BD) simulations have been conducted for the cryo-EM structures of the BA.2 S trimer with two human ACE2 bound (pdb id 7XO7) [22], BA.2 S trimer with two three ACE2 bound (pdb id 7XO8) [22], BA.2.75 S trimer with one human ACE2 bound (pdb id 7YR2) [25] and XBB.1 S trimer with one human ACE2 bound [31]. The profiles showed the appreciable level of thermal fluctuations in the generally adaptable S1 subunit (residues 14-530) including NTD (residues 14-306) and moderate displacements of the RBD-up regions (residues 331-528) that are stabilized via binding interactions with ACE2 and a more rigid S2 subunit (residues 686-1200) (Figure 5). We observed markedly larger fluctuations in the NTD and RBD regions for the XBB.1 1RBD-up complex, showing significantly more flexible S1 subunit where both ACE2-bound RBD-up and two closed RBDs displayed an appreciable mobility (Figure 5D) as compared to more stable S1 regions in the BA.2 (Figure 5A,B) and BA.2.75 trimer complexes (Figure 5C). We also observed differences in the NTD mobility including NTD N-terminus (residues 14–20), N3 (residues 141–156), and N5 (residues 246–260) that form an antigenic supersite on the NTD. These regions showed the increased RMSF values for all variants, but conformational heterogeneity in the NTD was particularly apparent in the BA.2 and XBB.1 complexes (Figure 5A,B,D). Three major fluctuation peaks were observed in the Wuhan NTD, residues 62–83, 140–158, 177–189, and 239–260 corresponding to the loop regions N2, N3, N4 and N5, respectively. The RMSF values for the ACE2-interacting RBD residues 440-456, 470-491 and 491-505 were small in the BA.2 and BA.2.75 complexes (Figure 5A-C) but these fluctuations were larger in the XBB.1 complex (Figure 5D). Conformational flexibility of the NTD and RBD in the open BA.2.75 trimer complexes is reduced, displaying smaller displacements within RMSF < 2.0-2.2 Å (Figure 5C). Hence, ACE2 binding may induce stabilization of the BA.2.75 S trimer by affecting not only the interacting RBD regions but also allosterically promoting the enhanced inter-protomer packing and rigidification of the S2 subunit. The observed dynamics pattern in the BA.2.75 S trimer complex is consistent with the experimental findings that the BA.2.75 S-trimer was the most stable, followed by BA.1, BA.2.12.1, BA.5 and BA.2 variants, also featuring the increased RBD rigidity and enhanced ACE2 binding affinity compared with other Omicron variants [25].

**Figure 5.**
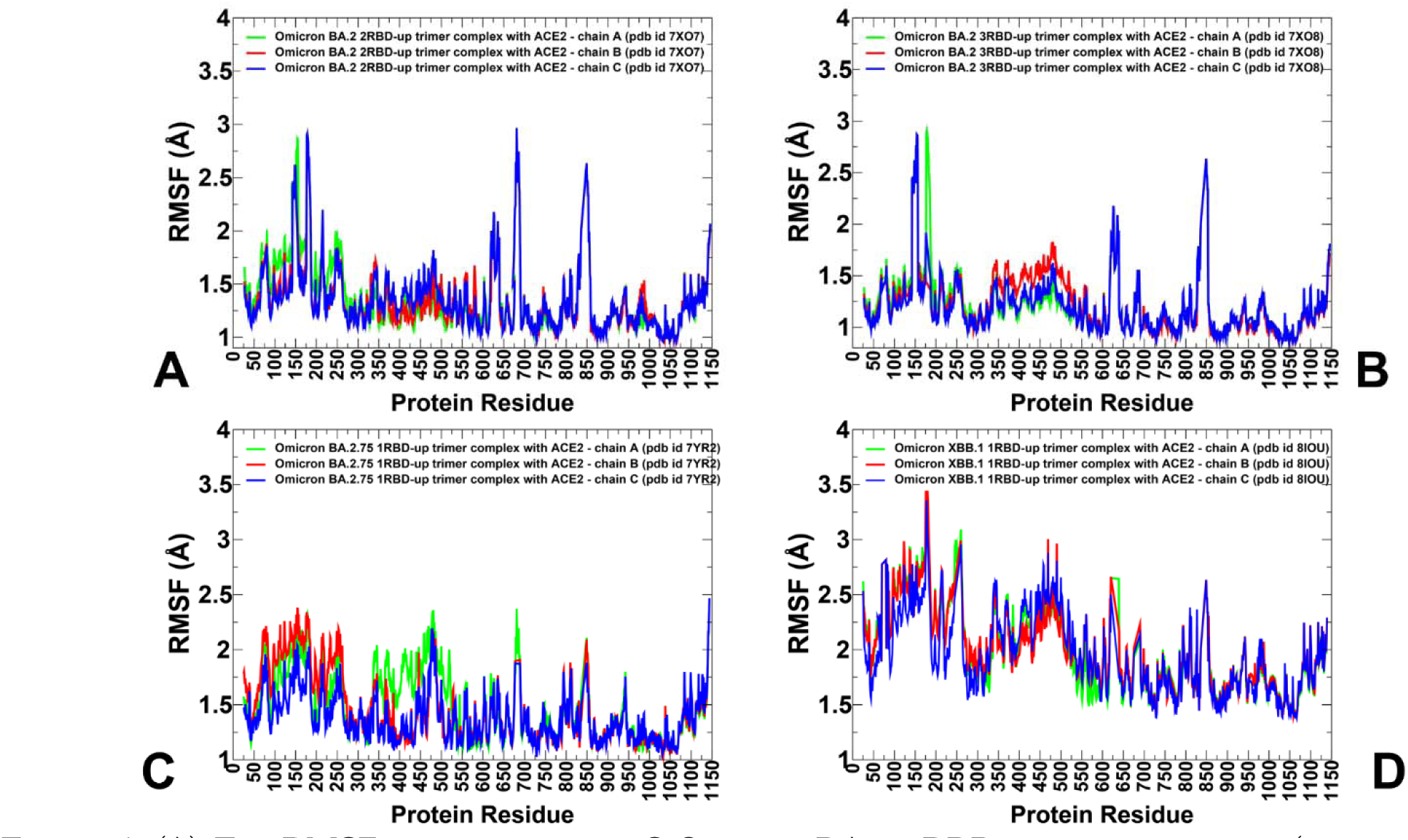
(A) The RMSF profiles for the S Omicron BA.2 2RBD-up trimer complex (pdb id 7XO7). (B) The RMSF profiles for the S Omicron BA.2 3RBD-up trimer complex (pdb id 7XO8). (C) The RMSF profiles for the S Omicron BA.2.75 1RBD-up trimer complex (pdb id 7YR2). (D) The RMSF profiles for the S Omicron XBB.1 1RBD-up trimer complex (pdb id 8IOU). The RMSF profiles are shown for protomer A in green lines, for protomer B in red lines, and for protomer C in blue lines.

In the Omicron BA.2 S trimer-ACE2 complexes, the RMSF profiles showed only relatively moderate fluctuations in the C-terminal domain 1, CTD1 region (residues 528-591) but more significant variations in the C-terminal domain 2, CTD2 (residues 592-686) (Figure 5A,B). Strikingly, we observed greater stability of both S1 and S2 regions in the BA.2.75 S trimer complex (Figure 5C) where NTD regions and RBD-up regions exhibited RMS displacements within 2.0-2.2 Å. The conformational dynamics profiles revealed larger deviations and greater flexibility for CTD2 (residues 592-686), particularly near the furin cleavage region (residues 670-690) (Figure5). The interesting patterns of conformational fluctuations were observed in the conserved and often considered tightly packed and rigid S2 subunit. The S2 regions fusion-peptide proximal region (FPPR) (residues 828-853), immediately downstream of the fusion peptide, the upstream helix (UH) (residues 736-781), central helix (CH) (residues 986-1035) are particularly rigid, UH (residues 736-781), heptad repeat HR1 (residues 910-985), central helix CH (residues 986-1035), CD, connector domain (residues 1035-1068), and heptad repeat HR2, heptad repeat 2; (residues 1069-1163) (Supplementary Materials, Figure S5). In all complexes with the exception of BA.2.75 complex, the RMSF displacements are significant near the S1/S2 cleavage site and in the FPPR region, while uniformly across all trimers only small deviations were seen in structurally stable UH and CH regions (Figure 5). Consistent with strong sequence conservation, the CH region is rigid in the all S trimer complexes. In particular, we found that ACE2 binding may promote stabilization of the trimeric interface stalk region of S2 (residues 899–913, 988–998, 1013–1021) (Figure 5). The stalk immobilization is particularly noticeable in the BA.2 and BA.2.7 complexes (Figure 5A-C). Only moderate displacements were seen in the HR1 region (residues 910-985) for BA.2 and BA..75 trimers but the flexibility in this region increases in the XBB.1 complex. In this context, it may be worth noting the results of HDX-MS experiments suggesting partial destabilization not only near the S1/S2 cleavage site but also in the HR1 region which may be functionally required to promote fusion stage. Our results showed considerable stability of the highly conserved hinge epitope (residues 980-1010) located in the S2 region between the CH and HR1 segments that is involved in regulation of conformational changes required for fusion of the viral envelope [46].

### Mutational Scanning of the Binding and Inter-Protomer Interactions in the Omicron BA.2, BA.2.75 and XBB.1 S Trimer-ACE2 Complexes

To provide a systematic residue-based mutational analysis of binding and stability in the Omicron S trimer-ACE2 complexes, we conducted mutational scanning of the RBD interface residues to evaluate ACE2 binding interactions and comprehensive scanning of the inter-protomer positions to estimate the packing interactions and differences in stability between Omicron trimers. Structure-based analysis of the Omicron RBD-ACE2 complexes showed considerable similarities as the differences in the number of intermolecular contacts were fairly small (Supplementary Materials, Tables S1-S3). We first constructed mutational heatmaps for the RBD binding interface residues (Figure 6). Unlike previous studies, these maps are based on scanning the RBD residues in the full length S-ACE2 complexes with a single or two ACE2 molecules present. In general, the results are in excellent agreement with the experimental deep mutagenesis scanning (DMS) data revealing a consistent group of predominantly hydrophobic energy hotspots (Figure 6). Indeed, common energetic hotspots Y453, L455, F456, F486, N487, Y489, Y501 and H505 also emerged as critical stability and binding hotspots in the experimental DMS studies [102,103]. Mutational heatmaps of the RBD-ACE2 binding interactions for all Omicron variants showed that the majority substitutions in these key interfacial positions can lead to loss in the stability and binding affinity with ACE2 (Figure 6).

**Figure 6.**
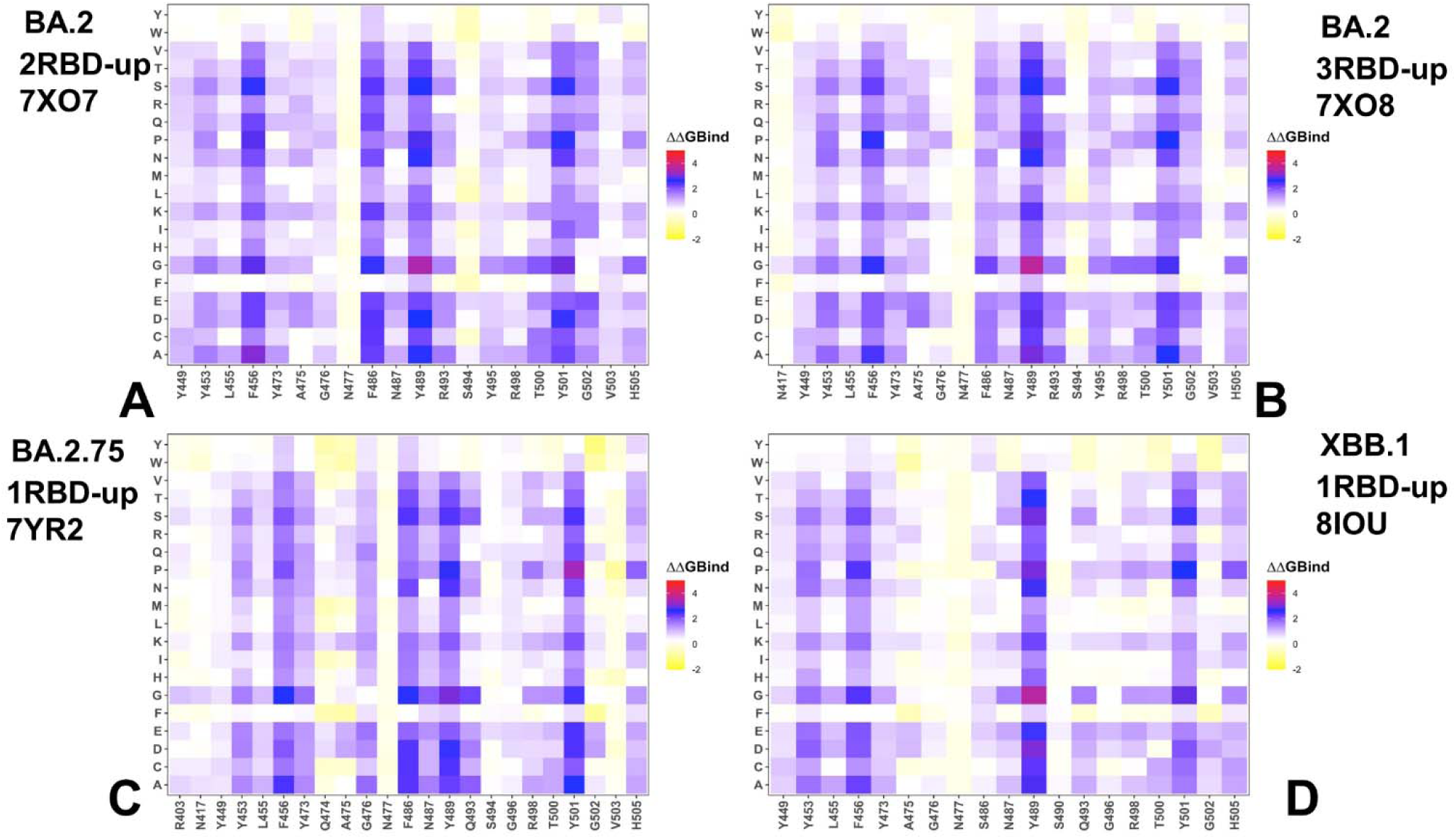
Ensemble-based mutational scanning of the RBD-ACE2 binding interfaces in the Omicron BA.2, BA.2.75 and XBB.1. S trimer-ACE2 complexes. The mutational scanning heatmaps are shown for the interfacial RBD residues in the S Omicron BA.2 2RBD-up trimer complex, pdb id 7XO7 (A), for the S Omicron BA.2 3RBD-up trimer complex, pdb id 7XO8 (B), for the S Omicron BA.2.75 1RBD-up trimer complex, pdb id 7YR2 (C) and for the S Omicron XBB.1 1RBD-up trimer complex, pdb id 8IOU (D). The heatmaps show the computed binding free energy changes for 20 single mutations of the interfacial positions. The standard errors of the mean for binding free energy changes using equally distributed 10,000 samples from the trajectories is 0.08-0.15 kcal/mol.

Hence, the conserved and stable hydrophobic RBD hotspots that are critical for RBD stability may be universally important for binding. Not surprisingly, Omicron mutations typically target more dynamic and vulnerable positions in the RBD to optimize virus fitness and balance the ACE2 binding affinity with immune escape potential. Interestingly, the mutational heatmaps for BA.2 trimer complexes with 2 ACE2 molecules (Figure 6A) and three bound ACE2 proteins (Figure 6B) showed considerable destabilization of the interfacial RBD residues where multiple positions (Y449, Y453, L455, F456, Y473, F486, N487, Y489, R493, R498, T500, and Y501) contributed significantly to the ACE2 binding. Compared to BA.1 variant BA.2 additionally contains S371F, T376A, D405N, and R408S mutations (Table 1). However, these RBD positions are outside of the binding interface and corresponding mutations may exert their effect through couplings with the interfacial sites. Overall, the structure of the S-BA.2 trimer complex with ACE2 revealed a more extensive interaction network at the binding interface and combined with stability of the BA.2 RBD could contribute to the high binding affinity of the BA.2 variant. BA.2.75 variant has additional mutations in the RBD (D339H, G446S, N460K, and R493Q). The mutational heatmap showed the increase in tolerance for positions R403, Y449, S493 and R498 where mutations result in moderate changes (Figure 6C). At the same time, the key hydrophobic sites Y453, L455, F456, F486, N487, Y489 as well as Y501 are preserved as binding hotspots in the BA.2.75 complex with a single ACE2 bound molecule. Previous studies suggested that R493Q reversal could restore the lost receptor-binding affinity due to N460K mutation [56,104]. In the BA.2 RBD/hACE2 complex, both R493 of RBD and K31 of human ACE2 are positively charged, which could decrease their binding, but a highly favorable salt bridge was observed between R493 of RBD and E35 of ACE2 (Figure 6C). Biophysical studies argued that R493Q substitution could improve the binding affinity towards human ACE2 [105]. At the same time, structural studies of the BA.2 trimer binding complexes with ACE2 used in our analysis suggested that Q493R forming a new salt bridge with E35 of human ACE2 and Q498R forming a new salt bridge with D38 of human ACE2 are key mutations resulting in an enhanced binding to human ACE2 [22]. This may explain the emergence of Q493R mutation in the first place in the BA.1 and BA.2 Omicron variants (Table 1). Our results showed that even though mutations of Q493 in BA.2.75 typically result in significant loss of binding affinity, Q493R substitution may not cause a dramatic decrease in binding (Figure 6C). Based on our findings the increased binding potential of BA.2.75 may be a cumulative effect resulting also from the increased stabilization of the RBD and S1-S2 regions. The reduced plasticity of the RBD and the inter-protomer/inter-domain regions may explain the increased sensitivity of BA.2.75 to neutralization [104].

Strikingly, the mutational heatmap for the XBB.1 trimer complex revealed smaller number of binding hotspots (Y453, F456, Y489 and Y501) while other positions including L455, Y473, A475, G476, N477, S486, S490 are more tolerant to substitutions that induce only small free energy changes (Figure 6D). These findings indicated that the increased flexibility of the XBB.1 trimer and binding interface combined with F486S and F490S mutations can result in decreased binding efficiency. Indeed, DMS studies have shown that F486P might enhance the affinity to ACE2 compared with F486S [103]. Moreover, the binding affinity of the BA.2.75 (K_D_ 1·8 nM) is much stronger than that of XBB.1 (K_D_ 19 nM) [32]. Mutational scanning analysis of the XBB.1 S-ACE2 binding is consistent with these experiments, revealing a more adaptable and mutation-tolerant interface which may also explain a significant immune escape potential of this variant traded in exchange of reduced ACE2 binding.

In addition, we also performed a systematic mutational scanning of the inter-protomer interfacial residues in the Omicron S trimer complexes to evaluate differences in the trimer packing and identify important structural stability hotspots (Figure 7). Strikingly, mutational analysis revealed clear and important differences between the variants. We found that the inter-protomer interaction contacts in the BA.2 trimer are generally sensitive to mutations, revealing strong destabilization changes in the S1 and S2 regions, including peaks aligned with NTD (residues F135, W152, F168, Y170, G199, Y200, P230, G257 and CTD1 interfaces (residues 542-547) (Figure 7A).

**Figure 7.**
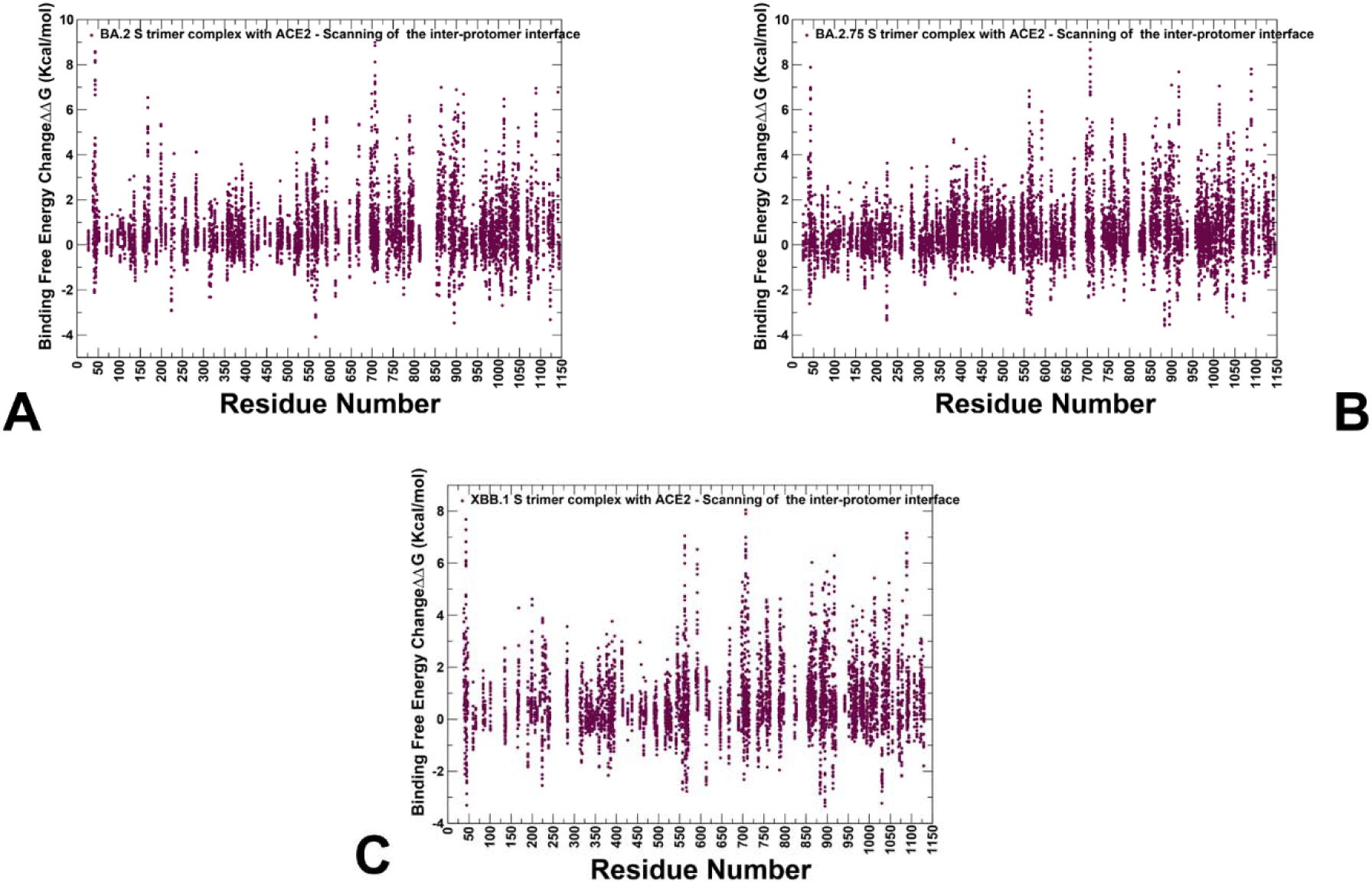
Ensemble-based mutational scanning of the inter-protomer interactions and protein stability for the Omicron BA.2, BA.2.75 and XBB.1 trimer complexes with ACE2. Mutational scanning scatter plots of the inter-protomer interactions is performed for residues identified as part of the protomer-protomer interface if solvent accessibility of a given residue in the complex is at least 5% lower than in the individual protomer. The interface residues are also examined as those on one protomer that have at least one non-hydrogen atom within 5 Å of the other protomer. The mutational scanning distributions are shown for the S Omicron BA.2 2RBD-up trimer complex, pdb id 7XO7 (A), for the S Omicron BA.2.75 1RBD-up trimer complex, pdb id 7YR2 (B) and for the S Omicron XBB.1 1RBD-up trimer complex, pdb id 8IOU (C). The heatmaps show the computed binding free energy changes for 20 single mutations of the interfacial positions. The standard errors of the mean for free energy changes were based on a different number of selected samples from a given trajectory (1,000, 5,000 and 10,000 samples) are within 0.15 kcal/mol.

The analysis showed that the inter-protomer NTD and RBD positions may form small clusters, but the density of these interfacial sites is smaller in BA.2 as compared to BA.2.75 trimer (Figure 7B). The analysis of mutational scanning showed large destabilization changes induced by mutations in the inter-protomer β-sheet segments involving residues 701–705 on one protomer and residues 780–790 from the other protomer (Figure 7A). These inter-protomer hotpots are preserved in all Omicron trimers. Interestingly, this region was identified in structure-guided mutagenesis to facilitate allosteric communication between NTD ligation and proteolytic fusion activation in the S2 region [106]. A spectrum of destabilization changes is seen in the S2 regions (residues 910-1150) (Figure 7A) reflecting the S2 inter-protomer interfaces while also showing some degree of conformational plasticity and energetic tolerance in the S2.

A significantly denser pattern of inter-protomer interface sites was seen in the BA.2.75 trimer complex (Figure 7B), revealing a number of inter-protomer residues in the NTD and RBD regions. These findings indicate that the inter-protomer contacts in BA.2.75 are stronger than in both BA.2 and XBB.1 trimers which is consistent with the increased structural stability of this Omicron variant. This may also have implications for detection and distribution of cryptic pockets. We argue that greater conformational plasticity of the BA.2 trimer as compared to more densely packed BA.2.75 trimer would affect evolution and emergence of dynamic cryptic sites. These results are consistent with the notion that Omicron open trimer structures are packed more tightly with enhanced inter-domain and inter-subunit interactions as compared to the Wu-Hu-1 WT S protein as each of the RBD-down protomers packs more tightly with its neighboring NTD [19].

In this mechanism, Omicron mutations promote a more stable open conformation. Our analysis showed that the inter-protomer stabilization progressively increased from BA.2 to BA.2.75 variant.

Strikingly, mutational scanning of the inter-protomer contacts in the XBB.1 trimer showed a distinct pattern, revealing fewer inter-protomer sites and visibly sparse distribution of these sites in the NTD and RBD (Figure 7C). Nonetheless, the distribution displayed several stability hotspots across all S regions including NTD (residues 38, 43, 47, 135, 168, 199, 200, 225, 232, 235), RBD (residues 396,413), CTD1 (residues 560-563), (residues 1047,1079). These results suggested that XBB.1 trimer may be less densely packed and has smaller inter-protomer interface which may contribute to the reduced stability and greater plasticity of XBB.1 variant. It may be argued that through enhanced plasticity XBB.1 variant may improve virus adaptability and enhance immune evasion potential. This may affect the distribution of cryptic pockets that could be more broadly distributed across both S1 and S2 regions.

### Detection of Cryptic Binding Pockets in the Conformational Ensembles of S Omicron Trimer Complexes with ACE2 : The Effects of Binding and Structural Plasticity in Mediating Networks of Conserved Allosteric Sites

Using the conformational ensembles of the Omicron S RBD-ACE2 complexes an S trimer-ACE2 complexes for BA.2, BA.2.75 and XBB.1 variants we performed a large scale cryptic pocket detection and comparative analysis of the putative binding sites in these systems. In particular, we investigated how variant-induced dynamic changes and ACE2 binding can affect the residue-based propensities for ligand binding (often termed ligandability preferences) and the distribution of cryptic pockets. Through this analysis, we examined the role of Omicro variant evolution and impact of recombinant Omicron sublineages from BA.2 to BA.2.75 and XBB.1 on the stability and population of cryptic binding sites. Here, we particularly focus on identifying conservation patterns and notable divergencies in the distribution of cryptic pockets for studied Omicron variants. We will investigate and validate a hypothesis that functionally important for allostery cryptic pockets formed at the inter-domain and inter-protomer interfaces in the S trimer complexes with ACE2 may be protected through evolution and shared between Omicron variants. There is the increasing interest in identifying druggable sites in the S2 subunit as most recently developed broad-spectrum fusion inhibitors and candidate vaccines can target the conserved elements in the S2 subunit [54].

First, we compared the residue-based pocket propensity profiles (Figure 8A-C) and the predicted RBD cryptic pockets in the RBD-ACE2 complexes (Figure 8D-F). The pocket propensity profiles for the BA.2 RBD (Figure 8A) featured a major dense peak in the RBD core (residues 360-390) and two small peaks (residues 426-432 and 510-515). The pocket detection analysis revealed two highly probable pockets. One pocket is formed by residues Y365, S366, Y369, F377, V382, S383, P384, L387, N388, F392, C432, and F515, while pocket 2 (residues F342, N343, A344, V367, L368, N370, A372, F374, W436, N437, L441, R509) is immediately adjacent to pocket 1 (Figure 8D). In general, the predicted pockets corresponded to a conserved and experimentally validated allosteric site in the RBD core where the essential free fatty acid LA binds [39–41]. According to the cryo-EM structure of the S complex with this small molecule the allosteric pocket is lined up by residues F338, V341, F342, F377, F374, F392, and W436 [39] which is consistent with our predictions. Notably, in the experimental S trimer complex LA binding occurs in a bipartite binding pocket in which the hydrophobic tail of LA molecule is bound to the RBD on one protomer, and the carboxyl headgroup of LA interacts with R408 Q409 of the neighboring RBD [39–41]. Interestingly, our analysis revealed that this cryptic pocket is highly probable and conserved even in the absence of the adjacent protomers. A small pocket on the other side of the BA.2-RBD is minor and much less probable. As a result, the population of ligandable RBD pockets in the BA.2 complex is dominated by the experimentally known LA allosteric site. A similar pocket propensity distribution in the BA.2.75 RBD-ACE2 complex displayed a dominant dense cluster peak for RBD residues 360-390 and smaller peaks for RBD core 430-440 and ACE2 interface positions 485-490 (Figure 8B). The most probable pocket (residues 338, 339, 342, 343, 367, 368, 371, 372, 374, 436, 437, 441) is aligned precisely along the LA allosteric site on the RBD (Figure 8E). The emergence of a single conserved allosteric pocket tin the BA.2.75 RBD may partly reflect structural rigidity of BA.2.75 and structural conservation of the major allosteric site. While the pocket propensity distribution for XBB.1 RBD is generally similar, we noticed the appearance of multiple isolated peaks (residues 367-371, 377-379, 432-434, 513-515) which may be due to the increased conformational plasticity of XBB.1 RBD (Figure 8C). The analysis revealed three equally probable smaller pockets (pocket 1-residues 365, 368, 369, 377, 387, 432, 434, 513, 515; pocket 2 -residues 343, 344, 345, 347, 371, 372, 436, 441, 509; and pocket 3 - residues 363, 365, 386, 387, 390, 391, 392, 395, 515, 524, 525) (Figure 8F). The three RBD cryptic subpockets are adjacent to each and other and combined covered much larger portion of the RBD surface (Figure 6F) Although these subpockets together overlay with the experimental allosteric site, we found a partial fragmentation of the cryptic site leading to formation of several smaller subpockets. Hence, our analysis suggested that variant-induced modulation of conformational plasticity may result in structural variations and expansion of the cryptic RBD site while the location and hydrophobic nature of this allosteric pocket is preserved in all complexes. We argue that this may implications for small molecule binding in the allosteric site likely reducing efficiency of ligands targeting the RBD.

**Figure 8.**
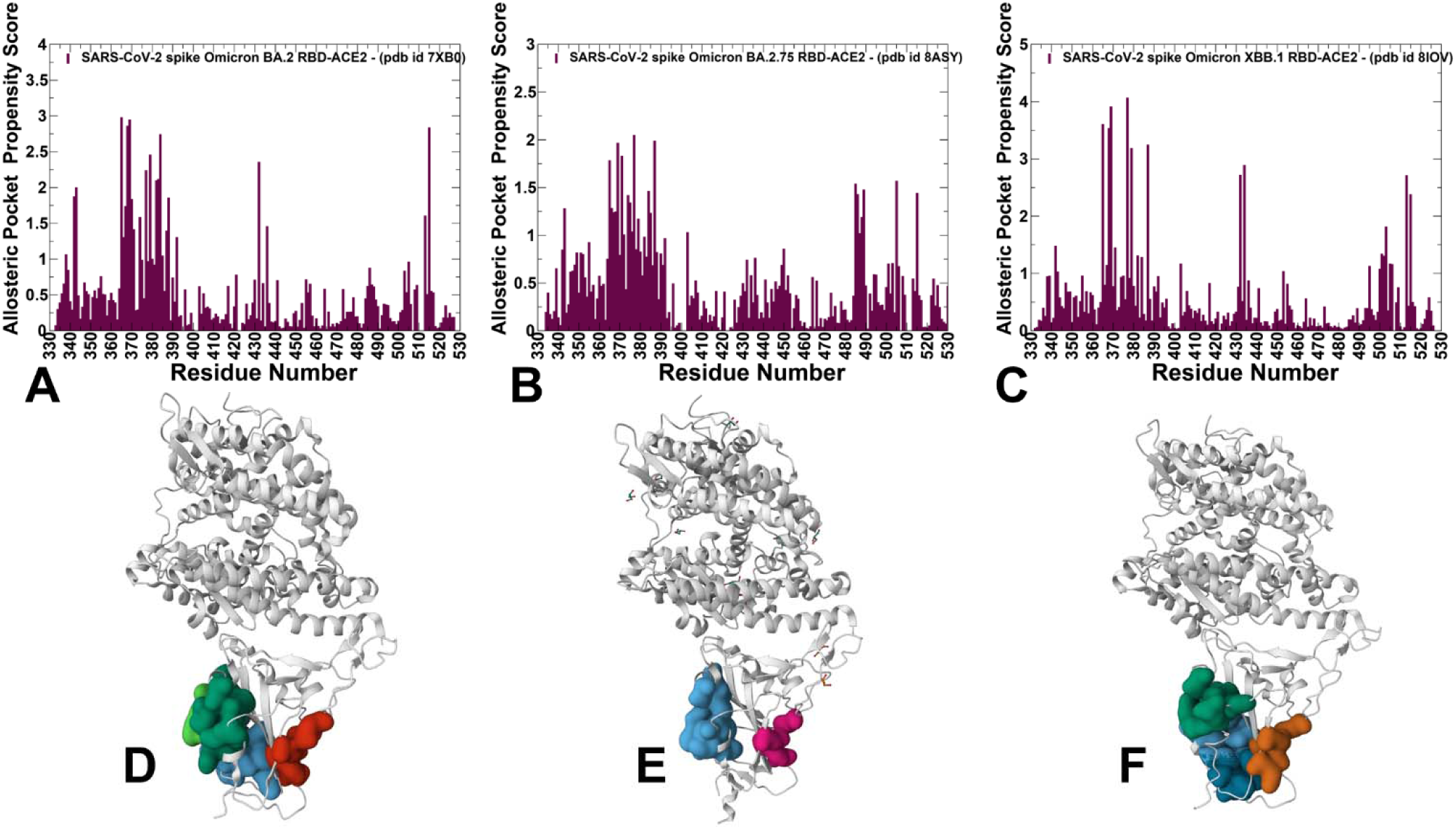
The residue-based pocket propensity distributions of the RBD residues for the BA.2 RBD-ACE2 complex (A), BA.2.75 RBD-ACE2 complex (B) and XBB.1 RBD-ACE2 complex (C). The profiles are shown in maroon-colored filled bars. Structural mapping of top 3 predicted binding pockets on representative RBD conformations in the BA.2 RBD-ACE2 complex (D), BA.2.75 RBD-ACE2 complex (E) and XBB.1 RBD-ACE2 complex (F). The ACE2 chain (chain A) and RBD chain (chain B) are shown in grey ribbons. The predicted pockets are shown in surface. The pockets shown for BA.2 RBD on panel D are : top ranked pocket 1 (B_365, B_366, B_369, B_377, B_382, B_383, B_384, B_387, B_388, B_392, B_432, B_515); pocket 2 (B_342, B_343, B_344, B_367, B_368, B_370, B_372, B_374, B_436, B_437, B_441, B_509) and minor pocket 3 (B_355, B_396, B_464, B_466, B_516). The pockets shown for BA.2.75 RBD on panel E are : top ranked pocket 1 (B_338, B_339, B_342, B_343, B_367, B_368, B_371, B_372, B_374, B_436, B_437, B_441) and minor pocket 2 (B_355, B_396, B_464, B_516). The pockets shown for XBB.1 RBD on panel F are : top ranked pocket 1 (B_365, B_368, B_369, B_377, B_387, B_432, B_434, B_513, B_515); second ranked pocket 2 (B_343, B_344, B_345, B_347, B_371, B_372, B_436, B_441, B_509), third ranked pocket 3 (B_363, B_365, B_386, B_387, B_390, B_391, B_392, B_395, B_515, B_524, B_525) and minor pocket 4 (B_355, B_357, B_396, B_464).

We then used conformational ensembles obtained from molecular simulations of the S trimer-ACE2 complexes to identify the available spectrum of potential cryptic sites. By ranking the pockets using residue-based propensity score for ligand binding we systematically analyzed the distribution and functional significance of these cryptic sites. The reported pocket propensity scores represent ensemble-averaged values obtained from P32Rank calculation based on the equilibrium trajectories (Figure 9). Here again, despite structurally similar structural arrangements of the open S trimer complexes we observed significant differences between pocket propensity profiles. A broad distribution with dense clusters of residues featuring peaks in both S1 and S2 subunits was seen for the BA.2 S-ACE2 complex (Figure 9A). Notably, one could differentiate three major peaks, with one corresponding to residue clusters in the RBD core, another cluster at the CTD1 region (residues 528-591) and pronounced peak in the S2 subunit pointing to CH region (residues 986-1035). Smaller peaks were also observed in the UH region (residues 736-781) and HR1 (residues 910-985). These findings suggested that the BA.2 S trimer could retain a considerable conformational plasticity in the ACE2-bound complex and feature a significant number of cryptic pockets in both S1 and S2 regions. Interestingly, the pocket propensity profile changes in the BA.2 S trimer complex with three ACE2 molecules bound to RBDs in the up position (Figure 9B). In this ensemble, we observed several sharp narrow peaks in the RBD and inter-domain CTD1 regions whereas smaller density was seen in the S2 regions. A different distribution profile was obtained for the BA.2.75 1 RBD-up trimer complex (Figure 9C) in which pronounced sharp peaks are aligned with the NTD and RBD regions while the pocket propensities of residues in the S2 subunit are markedly reduced. Accordingly, this may indicate that diverse cryptic pockets in the BA.2.75 trimer complex may be localized in the NTD and RBD, while the tightly packed S2 subunit may harbor a smaller number of conserved binding pockets. Notably, the pocket propensity profile of the XBB.1 1RBD-up trimer complex (Figure 9D) is similar to BA.2 2RBD-up trimer (Figure 9A) featuring multiple cluster peaks in the NTD, RBD, S1/S2 cleavage region and stalk region of the S2 subunit.

**Figure 9.**
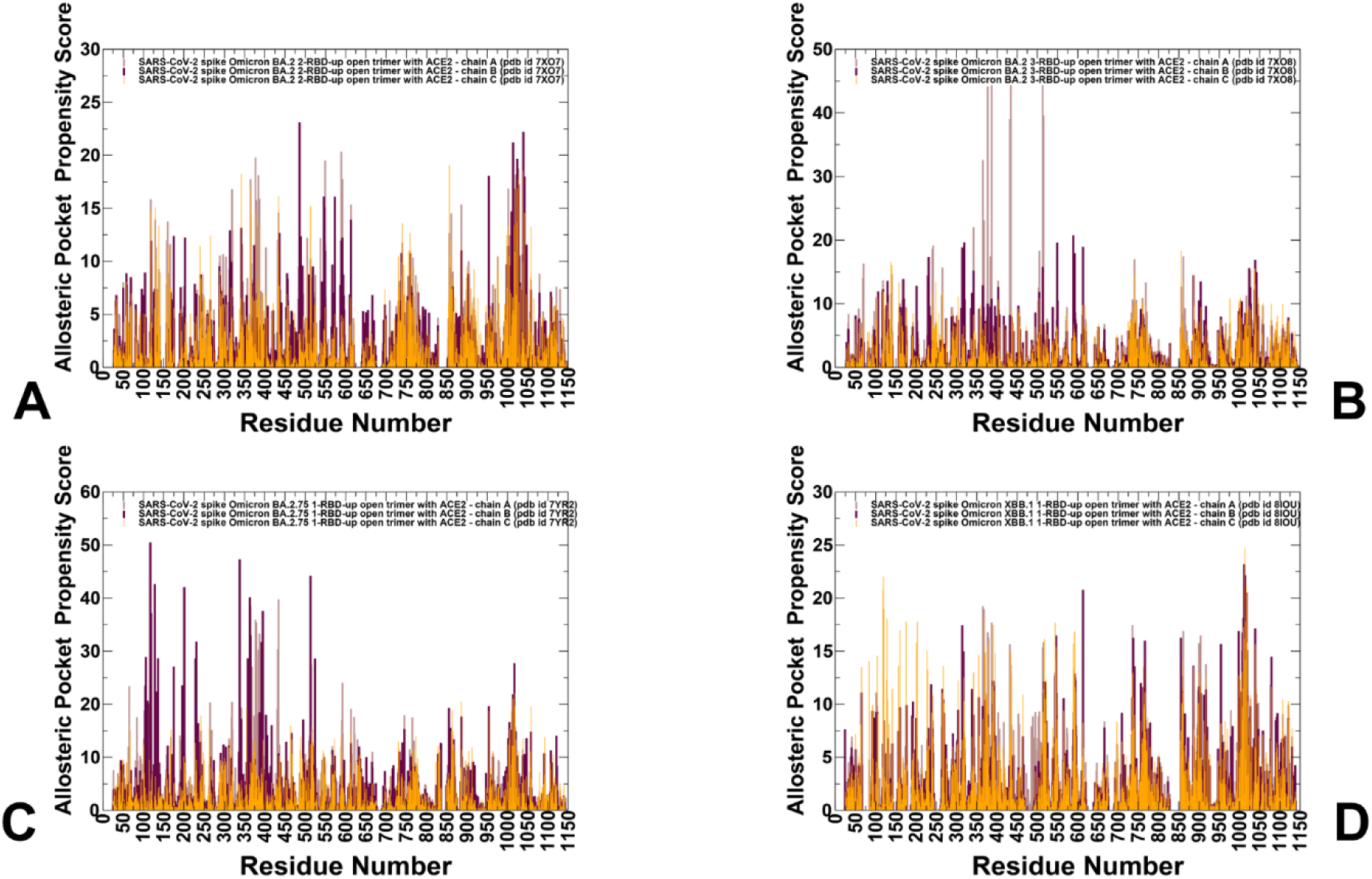
The residue-based pocket propensity distributions for the S Omicron BA.2 2RBD-up trimer complex with ACE2, pdb id 7XO7 (A), Omicron BA.2 3RBD-up trimer complex with ACE2, pdb id 7XO8 (B), Omicron BA.2.75 1RBD-up trimer complex with ACE2, pdb id 7YR2 (C) and Omicron XBB.1 1RBD-up trimer complex with ACE2, pdb id 8IOU (D). The profiles in each structure are shown for the three protomers, protomer A in light brown bars, protomer B in maroon bars, and protomer C in orange bars.

It is interesting to relate our findings with the HDX-MS experiments showing that ACE2 binding may result in stabilization of the CH (residues 986-1035) and CD regions (resides 1035-1068) including S2 stalk segments (regions 899–913, 988–998, 1013–1021) while induce the increased mobility in the S1/S2 cleavage site and HR1 domain [36,37]. According to our predictions, the high pocket propensity regions are generally distributed in the dynamic NTD and RBD regions as well as in S2 subunit. For BA.2 and XBB.1 trimer complexes, we observed a dense broad peak corresponding to HR1 and CH regions (Figure 9A,D), suggesting a potential for emergence of cryptic pockets in these conserved S2 regions. Overall, the results revealed that cryptic binding pockets may emerge along the S trimer architecture in the dynamic S1 regions as well as conserved and stable S2 segments.

By using the distribution-based ranking of the binding pockets, we characterized and classified the probable cryptic pockets in the Omicron complexes. In particular, we focused on the identification of evolutionary conserved and stable sites that may be shared among variants as well as potentially variant-specific unique pockets. The structural analysis of the top ranked binding pockets in the S trimer complexes revealed conserved allosteric pockets shared between Omicron variants, while displaying considerable difference in the composition, size and distribution in the S1 and S2 regions (Figure 10). The distribution of the top five predicted cryptic pockets in the Omicron S trimer complexes showed that they predominantly occupy the NTD and NTD-RBD regions. However, we also detected top ranked binding pockets in the stalk region of the S2 trimer interface, which emerged in all three complexes (Figure 10). Interestingly, among the top cryptic pockets are the inter-domain binding sites seen in the BA.2 and BA.2.75 trimer complexes (Figure 10A,B) but not found in the XBB.1 complex (Figure 10C).

**Figure 10.**
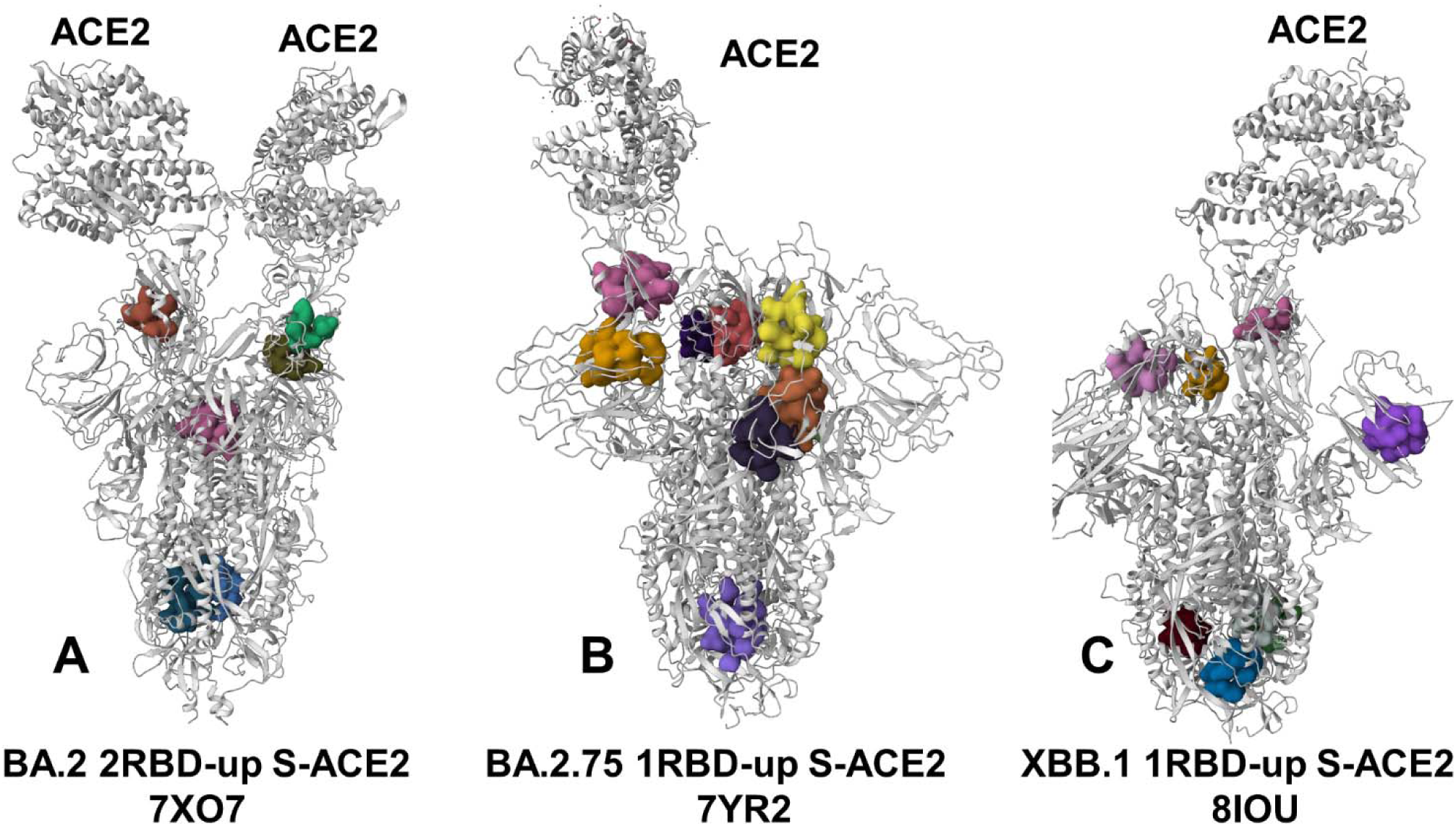
Structural maps of top ranked pockets for the S Omicron BA.2 2RBD-up trimer complex with ACE2, pdb id 7XO7 (A), Omicron BA.2.75 1RBD-up trimer complex with ACE2, pdb id 7YR2 (B) and Omicron XBB.1 1RBD-up trimer complex with ACE2, pdb id 8IOU (C). The structures of Omicron trimer complexes are shown in grey ribbons and predicted binding pockets are shown in surface representation.

To provide a systematic characterization of predicted pockets and identify conserved and variant-specific binding sites, we embarked on a more detailed structure-functional analysis. First, we found that the distribution of the pockets in BA.2 complex is consistent with the conformational adaptability of the S-BA.2 trimer featuring pockets in the NTD-RBD regions, the inter-protomer S1-S2 regions and also in the S2 subunit (Figure 10A, Supplementary Materials, Figure S6, Supplementary Materials, Table S4).

The cryptic pockets formed in the inter-protomer NTD-RBD regions include pocket 1 (NTD:B_115, B_132, B_167, B_168, B_170, B_230, B_231, B_232, RBD: C_357, C_393, C_394, C_518, C_520, C_521, C_523) and pocket 2 (RBD:: A_357, A_359, A_360, A_393, A_394, A_520, A_521, A_523, NTD: C_115, C_130, C_168, C_230, C_231) (Supplementary Materials, Figure S6). A close-up view of the predicted NBD-RBD pocket in the up protomer interacting with ACE2 highlighted the exposed nature of this pocket in the open conformation which is not obstructed by ACE2 binding and interactions with other protomers (Supplementary Materials, Figure S6). Our analysis suggested that targeting this site may allosterically weaken the RBD-ACE2 interactions even in the RBD-up open form. Our analysis also predicted a cryptic pocket formed at the RBD-NTD interface between RBD-up protomer bound to ACE2 and NTD of the adjacent protomer (Supplementary Materials, Figure S5). In this case, RBD residues R357, T393, N394, L518 and T523 from the other side of the RBD-up protomer form a pocket with the NTD residues Q115, E132, T167,F168, P230, I231 and G232 of the neighboring protomer located near the NTD supersite region. Interestingly, both pockets engage RBD residues from the ACE2-interacting up protomer. It is possible that targeting this binding pocket can alter binding strength of the BA.2 trimer with ACE2 and restrict NTD/RBD movements thus limiting conformational adaptability of the RBD-up states. We also found a cryptic binding pocket in the S2 subunit formed by all three protomers in which residues 756, 759, 994, 995, 998 and 1002 from each protomer create a well-defined pocket (Supplementary Materials, Figure S6). This pocket is formed by conserved residues from structurally stable UH region (residues 736-781) and CH region (residues 986-1035). Strikingly, the predicted inter-protomer S2 pocket overlaps with the highly conserved, conformational hinge epitope spanning residues 970–1006 of the SARS-2 spike, at the apex of the S2 domain [46]. This conserved pocket is shared among Omicron variants and is located at the critical functional hinge region involved in modulation S1-S2 opening motions. The experimental studies indicated that S dynamics may impact hinge epitope accessibility [46] and our results showed that this cryptic epitope may be accessible for ligand binding in the S trimer complexes with ACE2.

The recent HDX-MS studies of ACE2-induced allosteric activation of the S protein demonstrated allosteric changes upon ACE2 binding that extend to the hinge region and the top of the central helical bundle of the S2 subunit [38]. The experiments showed that ACE2 binding allosterically perturbs and primes HR1 and regions flanking the S1/S2 cleavage site for cleavage [36–38]. Targeting the predicted conserved cryptic pocket in the hinge apex region may potentially interfere with ACE2-induced destabilization of the HR1 helical structure (residues 930-960) that facilitates the pre-to-post-fusion spike conformational change. This pocket is shared between BA.2 and BA.2.75 complexes but was not stable in the XBB.1 complex suggesting some expansion and thermal breezing of the apex helices preventing three protomers from coming together to form a binding site.

We also found the inter-protomer cryptic pocket formed near S1-S2 hinge region formed by residues functional residues of one protomer (740, 741, 744, 745, 855, 856, 966, 976) and C TD1 residues from the neighboring protomer (548, 549, 571, 572, 573, 589). The importance of this inter-protomer interface residues has been firmly established in experimental studies, showing the quadruple mutant (A570L/T572I/F855Y/N856I) introducing modifications in these positions can shift the equilibrium in the S protein and affect the population of closed and open states [2]. These findings showed that potentially druggable pockets can be formed in the conserved S2 regions of the Omicron trimer complexes. It was also proposed that S2-targeted antibodies and vaccines that are developed to target rigid S2 are likely to be effective for various Omicron variants provided that evolution would continue to favor stabilization of the S2 stalk regions. As a result, design of small molecules for specific targeting of these pockets may potentially affect the ACE2 binding and interfere with S activation.

A considerable redistribution of the cryptic pockets was detected in the BA.2.75 complex (Figure 10B) where most of the predicted sites are in the NTD and RBD regions, while only several minor pockets were found in the peripheral S2 regions (Supplementary Materials, Figure S7). The differences in the distribution of cryptic binding pockets between BA.2 and BA.2.75 complexes are consistent with the increased stability of BA.2.75. We found that BA.2.75 S trimer complex featured well-defined and unobstructed RBD allosteric pocket in all protomers (residues 336, 338, 342, 358, 363, 364, 365, 368, 387, 392, 395, 397, 511, 513, 515, 524) (Supplementary Materials, Figure S7). Given that binding of small molecules in this LA-binding site is known to stabilize the closed S trimer [39–41], our predictions suggest that the RBD allosteric site can be targeted in the ACE2-bound Omicron complex, and potentially induce allosteric destabilization of the RBD-up open state. The analysis also revealed several binding pockets in the NTD : pocket formed by residues 101, 104, 117, 119, 121, 128, 170, 172, 175, 190, 192, 194, 201, 203, 227, 228, 229, 231 ; and pocket formed by residues 100, 101, 239, 240, 242, 248, 249, 250, 263, 264, 265, 65, 66, 81, 84, 94, 95, 96 (Supplementary Materials, Figure S7). Some of these NTD residues include residues 245–264 of the NTD supersite loop. The important revelation of this analysis is detection of several inter-protomer cryptic pockets formed near S1-S2 hinge region. These pockets bring CTD1 residues on one protomer (positions 541, 546, 547, 548, 549, 570, 572, 573, 587, 589, 592) and S2 residues from the adjacent protomer (positions 1000, 740, 741, 744, 745, 855, 856, 966, 976, 977, 978) (Supplementary Materials, Figure S6). A similar pocket was also detected among top ranked sites in the BA.2 S trimer complex. Our previous studies [107–110] and experimental data [2] showed that hinge residue F592 in one protomer can form the inter-protomer cluster with K854, F855, N856, and T859 hinge sites of the adjacent protomer (Supplementary Materials, Figure S7, Supplementary Materials, Table S5). This cryptic site provides a fairly small and deep pocket formed by critically important inter-protomer residues. Indeed, positions S591 and F592 are conserved structurally stable hinge sites of collective motions that can modulate both the inter-protomer and inter-domain changes [107–110]. This binding pocket is located immediately next to the ordered FPPR motif (residues 823-858) that engages in allosteric cross-talk with the RBD regions. Moreover, residues K854, Y855 and I856 are critical sites involved in the inter-protomer contacts and changes in these positions can modulate the shifts between the open and closed sites. The emergence of viable cryptic pockets formed by these regulatory positions can provide a rational strategy for allosteric targeting and modulation of S activity. The pocket detection also revealed the presence of conserved cryptic pocket formed between two protomers in the connector domain CD of S2 which links CH and the C-terminal HR2 (Supplementary Materials, Figure S7, Supplementary Materials, Table S5). The sensitivity of the cryptic pockets formation to the conformational dynamics is more pronounced for the XBB.1 trimer complex (Figure 10C, Supplementary Materials, Figure S8, Supplementary Materials, Table S6). Consistent with the greater flexibility of the XBB.1 complex observed in simulations, we found a heterogeneous distribution of the pocket propensity scores and respectively a more diverse structural allocation of the predicted pockets (Supplementary Materials, Figure S8, Supplementary Materials, Table S6). The predicted cryptic pockets are located in the NTD, RBD, CTD1) and S2 regions. Notably, different pockets are found in the S2. Interestingly, the composition of the pockets indicated an increased number of small pockets as conformational variability may limit formation of large and rigid pockets in the XBB.1 complex. The observed plasticity of the XBB.1 pockets also indicates that these pockets may readily adapt and alter which may be evolutionary beneficial for virus to increase immune evasion and mediate antibody resistance.

We detected pockets in the NTD and RBD regions that are similar to the ones seen in other Omicron variants. The divergences in composition of cryptic pockets for XBB.1 trimer is mainly in S2 regions where the predicted binding sites engage residues from UH, CH and CD functional segments (Supplementary Materials, Figure S8, Supplementary Materials, Table S6). Interestingly, we did not find the inter-protomer CTD1-S2 hinge pocket in the XBB.1 while this pocket is found as highly probable in more stable BA.2 and BA.2.75 trimer structures.

To summarize, our detailed analysis of the cryptic binding pockets in the Omicron complexes revealed a number of conserved and functionally important sites in the NTD, RBD regions as well as near the inter-protomer and inter-domain hinges. Unexpectedly, the results unveiled severa1 conserved cryptic pockets located deeply in the S2 subunit at the intersection of UH, CH and HR1 regions. Importantly, our findings pointed to divergences in the distribution of cryptic sites in a more dynamic XBB.1 trimer complex, resulting in the increased number of smaller pockets in less rigid S2 subunit while preserving the composition of cryptic binding sites in the NTD and RBD regions. Overall, the predicted binding pockets captured the experimentally known allosteric sites in the RBD, NTD and S2 apex hinge regions that emerged as top ranked pockets in the computational analysis.

## Discussion

In this study, we performed a systematic comparative analysis of conformational dynamics, binding energetics, and protein stability in the RBD-ACE2 complexes and full S trimer-ACE2 complexes for BA.2, BA.2.75 and XBB.1 variants. Based on simulations and using conformational ensembles for these systems, we also carried out a comprehensive analysis of evolution of cryptic binding pockets in the Omicron variants. XBB.1 subvariant is a descendant of BA.2 and recombinant of BA.2.10.1 and BA.2.75 sublineages. Conformational dynamics and MSM analysis confirmed that a combination of F486S and F490S mutations in XBB.1 may induce the increased RBD mobility in regions important for antibody binding and thereby enhance immune evasion potential of this Omicron variant. The MSM analysis suggested that BA.2 and XBB.1 RBDs may exhibit a similar degree of conformational plasticity in the complex, while BA.2.75 RBD is characterized by pronounced structural rigidity featuring a single dominant microstate.

Among central results of our study is evidence of progressive stabilization of the RBD regions and also inter-protomer interfaces in BA.2 and BA.2.75 variants. At the same time, XBB.1 trimer may be less densely packed and has smaller inter-protomer interface which may contribute to the reduced stability and greater plasticity of XBB.1 variant. It may be argued that through enhanced plasticity XBB.1 variant may improve virus adaptability and enhance immune evasion potential. The importance of these findings can be appreciated in the context of experimental data showing that Omicron BA.1, BA.2 and BA.2.75 open trimer structures can be characterized by progressively enhanced inter-domain interactions and improved packing of the inter-protomer interfaces, leading to more stable open states of the S protein for these variants [19]. Our analysis of XBB.1 trimer complex showed looser packing and weakened interfacial contacts resulting from the increased conformational flexibility. As appeared, this may alter the composition of functional pockets near inter-protomer hinges and affect the population distribution of closed and open states. Strikingly, cryo-EM studies of XBB.1 ectodomain and the XBB.1 S-ACE2 complex showed two closed states (closed-1 and closed-2) for the unbound XBB.1 trimer [31]. Moreover, the structure of the RBD one-up state, which is stable for BA.2 and BA.2.75 even in the absence of ACE2, was hardly observed in the structures of XBB.1LJS protein [31]. Our analysis is consistent with these data confirming the dynamic nature of the XBB.1 trimer which may alter cryptic sites and remodel conserved inter-protomer hinge sites seen in BA.2 and BA.2.75 structures. The increased conformational mobility and variability of cryptic pockets in XBB.1 can be also related with HDX-MS data showing S2 adaptability in the Omicron S protein which is propagated to HR1 and CH regions [35]. As a result, sequestering some of these S2 pockets in XBB.1 through selective targeting may rigidify specific XBB.1 state and disrupt dynamic changes priming S protein for the fusion events.

One of the important issues addressed in our study is how evolutionary changes in the Omicron variants affect stability and distribution of cryptic sites and what is the functional role of various binding sites in the context of S protein mechanisms and activity. Mutational changes in the Omicron trimers preserved druggable NTD and RBD pockets that are ranked as the most probable across all studied systems. Conformational flexibility of the NTD regions that harbor sites of conformational and mutational divergence still retains several cryptic pockets that are shared among Omicron variants. The consistent emergence of cryptic pockets in the NTD across all Omicron complexes showed that these hidden epitopes may be available for targeting even in the presence of bound ACE2 in open trimers. These results support recent data suggesting that the NTD can serve as an adaptable antigenic surface capable of unlocking cryptic binding pockets which may enable the efficient immune escape [35]. The pocket detection analysis consistently uncovered the allosteric RBD site targeted by LA molecule [39–41].

The results of our study suggested that despite general rigidity of the S2 regions in comparison with more dynamic S1 subunit, there is still an appreciable level of conformational adaptability in the S2, resulting in significant number of dynamic cryptic pockets in the S2. In particular, our data pointed to several conserved pockets in the HR1 and CH regions of the rigid S2 subunit, thereby indicating that an appreciable level of plasticity is present in S2 regions, giving rise to the broader accumulation of dynamic cryptic pockets. There has been a surge of interest in developing broad-spectrum fusion inhibitors, including antibodies, peptides and small molecules as well as vaccines targeting the conserved elements in the S2 subunit such as the fusion peptide, stem helix, and heptad repeats 1 and 2 (HR1-HR2) bundle. These targetable elements emerged as promising targets for the design of small molecules that can modulate various mechanisms of action such as fusion mechanism of action and allostery-based modulators regulating the experimentally established cross-talk between fusion peptide regions and RBD regions. The results of our study revealed several conserved and druggable cryptic pockets in the regulatory hinge regions of CTD1-S2 and S2 that are involved in communication between functional elements of S2 and RBD. Targeted ligand screening in the predicted druggable sites can allow for design of modulators of the S activity and facilitate development of chemical probes of S functions.

## Conclusions

In the current study we performed a systematic comparative analysis of the conformational dynamics, allostery and cryptic binding pockets in the RBD-ACE2 complexes trimers and S trimer complexes with the ACE2 receptor. Multiple microsecond MD simulations and Markov state model (MSM) analysis of the RBD-ACE2 complexes for the Omicron BA.2, BA.2.75 and XBB.1 variants enabled a detailed analysis of conformational states and populations. Using a comparative MSM analysis we showed an exquisite stability signature of the BA.2.75 RBD which could be contrasted with the increased mobility of the XBB.1 RBD. Using conformational ensembles of the SARS-CoV-2 Omicro S trimers, we conducted a systematic binding pocket screening and analysis of functional cryptic pockets in the BA.2, BA.2.75 and XBB.1 complexes with ACE2. The results of this study connected insights from conformational dynamics analysis, comparative mutational scanning of the S-ACE2 binding and the inter-protomer interactions with the evolution of cryptic binding sites. We demonstrated that our approach could reproduce all experimentally known allosteric sites and identify networks of conserved cryptic pockets preserved in different conformational states of the Omicron variants and in the S-ACE2 complexes. The results of our study suggest that despite general rigidity of the S2 regions in comparison with more dynamic S1 subunit, there is still an appreciable level of conformational adaptability in the S2, resulting in significant number of dynamic cryptic pockets. The determined cryptic binding pockets at the inter-protomer regions and in the functional regions of the S2 subunit such as HR1-HR2 bundle and stem helix region, are consistent with the role of the pocket residues in modulating conformational transitions and antibody recognition. Of particular interest is the detection of highly probable pockets in the S2 hinge regions known to be allosterically linked with the S1 movements during activation. The results detailed how mutational and conformational changes in the BA.2 and BA.2.75 spike trimers can modulate the distributions and mediate networks of inter-connected conserved and also variant-specific druggable allosteric pockets. These findings can be important for understanding mechanisms underlying functional roles of druggable cryptic pockets that can be used for both site-specific and allostery-inspired therapeutic intervention strategy targeting distinct conformational states of the SARS-CoV-2 Omicron variants. This may enable engineering of allosteric modulators that could rationally target a complex functional landscape of virus transmissibility.

## Supporting information

Supprting Figures S1-S8 and Tables S1-S6

## Author Contributions

Conceptualization, G.V.; methodology, S.X., P.T., G.V.; software, S.X., P.T., G.V., M.A. and G.G; validation, S.X., P.T., M.A., G.G., G.V.; formal analysis, G.V., M.A. and G.G.; investigation, P.T., G.V.; resources, S.X., P.T., G.V., M.A. and G.G.; data curation, S.X., G.G., M.A., G.V.; writing—original draft preparation, G.V.; writing—review and editing, G.V., M.A. and G.G.; visualization, G.V.; supervision, G.V.; project administration, P.T., G.V.; funding acquisition, P.T., G.V. All authors have read and agreed to the published version of the manuscript.

## Institutional Review Board Statement

Not applicable.

## Informed Consent Statement

Not applicable.

## Data Availability Statement

Data is fully contained within the article. Crystal structures were obtained and downloaded from the Protein Data Bank (http://www.rcsb.org). All simulations were performed using NAMD 2.13 package that was obtained from website https://www.ks.uiuc.edu/Development/Download/. All simulations were performed using the CHARMM36 force field obtained from http://mackerell.umaryland.edu/charmm_ff.shtml. The rendering of protein structures was done with interactive visualization program UCSF ChimeraX (https://www.rbvi.ucsf.edu/chimerax/) and Pymol (https://pymol.org/2/).

## Acknowledgments

The authors acknowledge support from Schmid College of Science and Technology at Chapman University for providing computing resources at the Keck Center for Science and Engineering.

## References

1. Cai, Y.; Zhang, J.; Xiao, T.; Peng, H.; Sterling, S. M.; Walsh, R. M., Jr.; Rawson, S.; Rits-Volloch, S.; Chen, B. Distinct conformational states of SARS-CoV-2 spike protein. Science 2020, 369, 1586–1592. doi: 10.1126/science.abd4251.

2. Henderson, R.; Edwards, R. J.; Mansouri, K.; Janowska, K.; Stalls, V.; Gobeil, S. M. C.; Kopp, M.; Li, D.; Parks, R.; Hsu, A. L., Borgnia, M.J.; Haynes, B.F.; Acharya, P. Controlling the SARS-CoV-2 spike glycoprotein conformation. Nat. Struct. Mol. Biol. 2020, 27, 925–933. doi: 10.1038/s41594-020-0479-4.

3. McCormick, K.D.; Jacobs, J.L.; Mellors, J.W. The emerging plasticity of SARS-CoV-2. Science 2021, 371, 1306–1308. doi: 10.1126/science.abg4493.

4. Ghimire, D.; Han, Y.; Lu, M. Structural Plasticity and Immune Evasion of SARS-CoV-2 Spike Variants. Viruses 2022, 14, 1255. 10.3390/v14061255.

5. Xu, C.; Wang, Y.; Liu, C.; Zhang, C.; Han, W.; Hong, X.; Wang, Y.; Hong, Q.; Wang, S.; Zhao, Q.; Wang, Y.; Yang, Y.; Chen, K.; Zheng, W.; Kong, L.; Wang, F.; Zuo, Q.; Huang, Z.; Cong, Y. Conformational dynamics of SARS-CoV-2 trimeric spike glycoprotein in complex with receptor ACE2 revealed by cryo-EM. Sci. Adv. 2021, 7, eabe5575. doi: 10.1126/sciadv.abe5575.

6. Benton, D. J.; Wrobel, A. G.; Xu, P.; Roustan, C.; Martin, S. R.; Rosenthal, P. B.; Skehel, J. J.; Gamblin, S. J. Receptor binding and priming of the spike protein of SARS-CoV-2 for membrane fusion. Nature 2020, 588, 327–330. doi: 10.1038/s41586-020-2772-0.

7. Turoňová, B.; Sikora, M.; Schürmann, C.; Hagen, W. J. H.; Welsch, S.; Blanc, F. E. C.; von Bülow, S.; Gecht, M.; Bagola, K.; Hörner, C.; van Zandbergen, G.; Landry, J.; de Azevedo, N. T. D.; Mosalaganti, S.; Schwarz, A.; Covino, R.; Mühlebach, M. D.; Hummer, G.; Krijnse Locker, J.; Beck, M. In situ structural analysis of SARS-CoV-2 spike reveals flexibility mediated by three hinges. Science 2020, 370, 203–208. doi: 10.1126/science.abd5223.

8. Lu, M.; Uchil, P. D.; Li, W.; Zheng, D.; Terry, D. S.; Gorman, J.; Shi, W.; Zhang, B.; Zhou, T.; Ding, S.; Gasser, R.; Prevost, J.; Beaudoin-Bussieres, G.; Anand, S. P.; Laumaea, A.; Grover, J. R.; Lihong, L.; Ho, D. D.; Mascola, J.R.; Finzi, A.; Kwong, P. D.; Blanchard, S. C.; Mothes, W. Real-time conformational dynamics of SARS-CoV-2 spikes on virus particles. Cell Host Microbe. 2020, 28, 880–891.e8. doi: 10.1016/j.chom.2020.11.001.

9. Yang, Z.; Han, Y.; Ding, S.; Shi, W.; Zhou, T.; Finzi, A.; Kwong, P.D.; Mothes, W.; Lu, M. SARS-CoV-2 Variants Increase Kinetic Stability of Open Spike Conformations as an Evolutionary Strategy. mBio 2022, 13, e0322721. doi: 10.1128/mbio.03227-21.

10. Díaz-Salinas, M.A.; Li, Q.; Ejemel, M.; Yurkovetskiy, L.; Luban, J.; Shen, K.; Wang, Y.; Munro, J.B. Conformational dynamics and allosteric modulation of the SARS-CoV-2 spike. Elife 2022, 11, e75433. doi: 10.7554/eLife.75433.

11. Hong, Q.; Han, W.; Li, J.; Xu, S.; Wang, Y.; Xu, C.; Li, Z.; Wang, Y.; Zhang, C.; Huang, Z.; Cong, Y. Molecular basis of receptor binding and antibody neutralization of Omicron. Nature 2022. doi: 10.1038/s41586-022-04581-9.

12. Gobeil, S. M.-C.; Henderson, R.; Stalls, V.; Janowska, K.; Huang, X.; May, A.; Speakman, M.; Beaudoin, E.; Manne, K.; Li, D.; Parks, R.; Barr, M.; Deyton, M.; Martin, M.; Mansouri, K.; Edwards, R. J.; Eaton, A.; Montefiori, D. C.; Sempowski, G. D.; Saunders, K. O.; Wiehe, K.; Williams, W.; Korber, B.; Haynes, B. F.; Acharya, P. Structural Diversity of the SARS-CoV-2 Omicron Spike. Mol. Cell 2022, 82, 2050–2068.e6. 10.1016/j.molcel.2022.03.028.

13. Cui, Z.; Liu, P.; Wang, N.; Wang, L.; Fan, K.; Zhu, Q.; Wang, K.; Chen, R.; Feng, R.; Jia, Z.; Yang, M.; Xu, G.; Zhu, B.; Fu, W.; Chu, T.; Feng, L.; Wang, Y.; Pei, X.; Yang, P.; Xie, X.S.; Cao, L.; Cao, Y.; Wang, X. Structural and functional characterizations of infectivity and immune evasion of SARS-CoV-2 Omicron. Cell 2022, 185, 860–871.e13. doi: 10.1016/j.cell.2022.01.019.

14. Zhou, T.; Wang, L.; Misasi, J.; Pegu, A.; Zhang, Y.; Harris, D.R.; Olia, A.S.; Talana, C.A.; Yang, E.S.; Chen, M.; Choe, M.; Shi, W.; Teng, I.T.; Creanga, A.; Jenkins, C.; Leung, K.; Liu, T.; Stancofski, E.D.; Stephens, T.; Zhang, B.; Tsybovsky, Y.; Graham, B.S.; Mascola, J.R.; Sullivan, N.J.; Kwong, P.D. Structural basis for potent antibody neutralization of SARS-CoV-2 variants including B.1.1.529. Science 2022, 376, eabn8897. doi: 10.1126/science.abn8897.

15. Guo, H.; Gao, Y.; Li, T.; Li, T.; Lu, Y.; Zheng, L.; Liu, Y.; Yang, T.; Luo, F.; Song, S.; Wang, W.; Yang, X.; Nguyen, H. C.; Zhang, H.; Huang, A.; Jin, A.; Yang, H.; Rao, Z.; Ji, X. Structures of Omicron Spike Complexes and Implications for Neutralizing Antibody Development. Cell Rep. 2022, 39, 110770. doi: 10.1016/j.celrep.2022.110770.

16. Stalls, V.; Lindenberger, J.; Gobeil, S. M.-C.; Henderson, R.; Parks, R.; Barr, M.; Deyton, M.; Martin, M.; Janowska, K.; Huang, X.; May, A.; Speakman, M.; Beaudoin, E.; Kraft, B.; Lu, X.; Edwards, R. J.; Eaton, A.; Montefiori, D. C.; Williams, W. B.; Saunders, K. O.; Wiehe, K.; Haynes, B. F.; Acharya, P. Cryo-EM Structures of SARS-CoV-2 Omicron BA.2 Spike. Cell Rep. 2022, 39, 111009. doi: 10.1016/j.celrep.2022.111009.

17. Lin, S.; Chen, Z.; Zhang, X.; Wen, A.; Yuan, X.; Yu, C.; Yang, J.; He, B.; Cao, Y.; Lu, G. Characterization of SARS-CoV-2 Omicron Spike RBD Reveals Significantly Decreased Stability, Severe Evasion of Neutralizing-Antibody Recognition but Unaffected Engagement by Decoy ACE2 Modified for Enhanced RBD Binding. Signal Transduct Target Ther. 2022, 7, 6. doi: 10.1038/s41392-022-00914-2.

18. Cerutti, G.; Guo, Y.; Liu, L.; Liu, L.; Zhang, Z.; Luo, Y.; Huang, Y.; Wang, H. H.; Ho, D. D.; Sheng, Z.; Shapiro, L. Cryo-EM Structure of the SARS-CoV-2 Omicron Spike. Cell Rep. 2022, 38, 110428. doi: 10.1016/j.celrep.2022.110428.

19. Ye, G.; Liu, B.; Li, F. Cryo-EM Structure of a SARS-CoV-2 Omicron Spike Protein Ectodomain. Nat Commun. 2022, 13, 1214. doi: 10.1038/s41467-022-28882-9.

20. Saville, J.W.; Mannar, D.; Zhu, X.; Srivastava, S.S.; Berezuk, A.M.; Demers, J.P.; Zhou, S.; Tuttle, K.S.; Sekirov, I.; Kim A.; Li, W.; Dimitrov, D.S.; Subramaniam, S. Structural and biochemical rationale for enhanced spike protein fitness in delta and kappa SARS-CoV-2 variants. Nat. Commun. 2022, 13, 742. doi: 10.1038/s41467-022-28324-6.

21. Li, L.; Liao, H.; Meng, Y.; Li, W.; Han, P.; Liu, K.; Wang, Q.; Li, D.; Zhang, Y.; Wang, L.; Fan, Z.; Zhang, Y.; Wang, Q.; Zhao, X.; Sun, Y.; Huang, N.; Qi, J.; Gao, G.F. Structural basis of human ACE2 higher binding affinity to currently circulating Omicron SARS-CoV-2 sub-variants BA.2 and BA.1.1. Cell 2022, 185, 2952–2960.e10. doi: 10.1016/j.cell.2022.06.023.

22. Xu, Y.; Wu, C.; Cao, X.; Gu, C.; Liu, H.; Jiang, M.; Wang, X.; Yuan, Q.; Wu, K.; Liu, J.; Wang, D.; He, X.; Wang, X.; Deng, S.J.; Xu, H.E.; Yin, W. Structural and biochemical mechanism for increased infectivity and immune evasion of Omicron BA.2 variant compared to BA.1 and their possible mouse origins. Cell Res. 2022, 32, 609–620. doi: 10.1038/s41422-022-00672-4.

23. Zhang, J.; Tang, W.; Gao, H.; Lavine, C. L.; Shi, W.; Peng, H.; Zhu, H.; Anand, K.; Kosikova, M.; Kwon, H. J.; Tong, P.; Gautam, A.; Rits-Volloch, S.; Wang, S.; Mayer, M. L.; Wesemann, D. R.; Seaman, M. S.; Lu, J.; Xiao, T.; Xie, H.; Chen, B. Structural and Functional Characteristics of the SARS-CoV-2 Omicron Subvariant BA.2 Spike Protein. Nat. Struct. Mol. Biol. 2023, 30, 980–990. 10.1038/s41594-023-01023-6.

24. Cao, Y.; Yisimayi, A.; Jian, F.; Song, W.; Xiao, T.; Wang, L.; Du, S.; Wang, J.; Li, Q.; Chen, X.; Yu, Y.; Wang, P.; Zhang, Z.; Liu, P.; An, R.; Hao, X.; Wang, Y.; Wang, J.; Feng, R.; Sun, H.; Zhao, L.; Zhang, W.; Zhao, D.; Zheng, J.; Yu, L.; Li, C.; Zhang, N.; Wang, R.; Niu, X.; Yang, S.; Song, X.; Chai, Y.; Hu, Y.; Shi, Y.; Zheng, L.; Li, Z.; Gu, Q.; Shao, F.; Huang, W.; Jin, R.; Shen, Z.; Wang, Y.; Wang, X.; Xiao, J.; Xie, X. S. BA.2.12.1, BA.4 and BA.5 Escape Antibodies Elicited by Omicron Infection. Nature 2022, 608, 593–602. 10.1038/s41586-022-04980-y.

25. Cao, Y.; Song, W.; Wang, L.; Liu, P.; Yue, C.; Jian, F.; Yu, Y.; Yisimayi, A.; Wang, P.; Wang, Y.; Zhu, Q.; Deng, J.; Fu, W.; Yu, L.; Zhang, N.; Wang, J.; Xiao, T.; An, R.; Wang, J.; Liu, L.; Yang, S.; Niu, X.; Gu, Q.; Shao, F.; Hao, X.; Meng, B.; Gupta, R. K.; Jin, R.; Wang, Y.; Xie, X. S.; Wang, X. Characterization of the Enhanced Infectivity and Antibody Evasion of Omicron BA.2.75. Cell Host Microbe 2022, 30, 1527–1539.e5. doi: 10.1016/j.chom.2022.09.018.

26. Chen, Z.; Li, J.; Zheng, J.; Jin, Y.; Zhang, Y.; Tang, F.; Li, J.; Cheng, H.; Jiang, L.; Wen, H.; Hong, C.; Zeng, X.; Huang, S.; Lu, B.; Li, L.; Wang, Z. Emerging Omicron Subvariants Evade Neutralizing Immunity Elicited by Vaccine or BA.1/BA.2 Infection. J. Med. Virol. 2023, 95, e28539. doi: 10.1002/jmv.28539.

27. Saito, A.; Tamura, T.; Zahradnik, J.; Deguchi, S.; Tabata, K.; Anraku, Y.; Kimura, I.; Ito, J.; Yamasoba, D.; Nasser, H.; Toyoda, M.; Nagata, K.; Uriu, K.; Kosugi, Y.; Fujita, S.; Shofa, M.; Monira Begum, M.; Shimizu, R.; Oda, Y.; Suzuki, R.; Ito, H.; Nao, N.; Wang, L.; Tsuda, M.; Yoshimatsu, K.; Kuramochi, J.; Kita, S.; Sasaki-Tabata, K.; Fukuhara, H.; Maenaka, K.; Yamamoto, Y.; Nagamoto, T.; Asakura, H.; Nagashima, M.; Sadamasu, K.; Yoshimura, K.; Ueno, T.; Schreiber, G.; Takaori-Kondo, A.; Shirakawa, K.; Sawa, H.; Irie, T.; Hashiguchi, T.; Takayama, K.; Matsuno, K.; Tanaka, S.; Ikeda, T.; Fukuhara, T.; Sato, K. Virological Characteristics of the SARS-CoV-2 Omicron BA.2.75 Variant. Cell Host Microbe 2022, 30, 1540–1555.e15. doi: 10.1016/j.chom.2022.10.003.

28. Qu, P.; Evans, J. P.; Zheng, Y.-M.; Carlin, C.; Saif, L. J.; Oltz, E. M.; Xu, K.; Gumina, R. J.; Liu, S.-L. Evasion of Neutralizing Antibody Responses by the SARS-CoV-2 BA.2.75 Variant. Cell Host Microbe 2022, 30, 1518–1526.e4. doi: 10.1016/j.chom.2022.09.015.

29. Cao, Y.; Jian, F.; Wang, J.; Yu, Y.; Song, W.; Yisimayi, A.; Wang, J.; An, R.; Chen, X.; Zhang, N.; Wang, Y.; Wang, P.; Zhao, L.; Sun, H.; Yu, L.; Yang, S.; Niu, X.; Xiao, T.; Gu, Q.; Shao, F.; Hao, X.; Xu, Y.; Jin, R.; Shen, Z.; Wang, Y.; Xie, X. S. Imprinted SARS-CoV-2 Humoral Immunity Induces Convergent Omicron RBD Evolution. Nature 2023, 614, 521–529. doi: 10.1038/s41586-022-05644-7.

30. Wang, Q.; Iketani, S.; Li, Z.; Liu, L.; Guo, Y.; Huang, Y.; Bowen, A. D.; Liu, M.; Wang, M.; Yu, J.; Valdez, R.; Lauring, A. S.; Sheng, Z.; Wang, H. H.; Gordon, A.; Liu, L.; Ho, D. D. Alarming Antibody Evasion Properties of Rising SARS-CoV-2 BQ and XBB Subvariants. Cell 2023, 186, 279–286.e8. 10.1016/j.cell.2022.12.018.

31. Tamura, T.; Ito, J.; Uriu, K.; Zahradnik, J.; Kida, I.; Anraku, Y.; Nasser, H.; Shofa, M.; Oda, Y.; Lytras, S.; Nao, N.; Itakura, Y.; Deguchi, S.; Suzuki, R.; Wang, L.; Begum, M. M.; Kita, S.; Yajima, H.; Sasaki, J.; Sasaki-Tabata, K.; Shimizu, R.; Tsuda, M.; Kosugi, Y.; Fujita, S.; Pan, L.; Sauter, D.; Yoshimatsu, K.; Suzuki, S.; Asakura, H.; Nagashima, M.; Sadamasu, K.; Yoshimura, K.; Yamamoto, Y.; Nagamoto, T.; Schreiber, G.; Maenaka, K.; Ito, H.; Misawa, N.; Kimura, I.; Suganami, M.; Chiba, M.; Yoshimura, R.; Yasuda, K.; Iida, K.; Ohsumi, N.; Strange, A. P.; Takahashi, O.; Ichihara, K.; Shibatani, Y.; Nishiuchi, T.; Kato, M.; Ferdous, Z.; Mouri, H.; Shishido, K.; Sawa, H.; Hashimoto, R.; Watanabe, Y.; Sakamoto, A.; Yasuhara, N.; Suzuki, T.; Kimura, K.; Nakajima, Y.; Nakagawa, S.; Wu, J.; Shirakawa, K.; Takaori-Kondo, A.; Nagata, K.; Kazuma, Y.; Nomura, R.; Horisawa, Y.; Tashiro, Y.; Kawai, Y.; Irie, T.; Kawabata, R.; Motozono, C.; Toyoda, M.; Ueno, T.; Hashiguchi, T.; Ikeda, T.; Fukuhara, T.; Saito, A.; Tanaka, S.; Matsuno, K.; Takayama, K.; Sato, K. Virological Characteristics of the SARS-CoV-2 XBB Variant Derived from Recombination of Two Omicron Subvariants. Nat Commun. 2023, 14, 2800. doi: 10.1038/s41467-023-38435-3.

32. Yue, C.; Song, W.; Wang, L.; Jian, F.; Chen, X.; Gao, F.; Shen, Z.; Wang, Y.; Wang, X.; Cao, Y. ACE2 Binding and Antibody Evasion in Enhanced Transmissibility of XBB.1.5. Lancet Infect. Dis. 2023, 23, 278–280. 10.1016/s1473-3099(23)00010-5.

33. Hoffmann, M.; Arora, P.; Nehlmeier, I.; Kempf, A.; Cossmann, A.; Schulz, S. R.; Morillas Ramos, G.; Manthey, L. A.; Jäck, H.-M.; Behrens, G. M. N.; Pöhlmann, S. Profound Neutralization Evasion and Augmented Host Cell Entry Are Hallmarks of the Fast-Spreading SARS-CoV-2 Lineage XBB.1.5. Cell Mol Immunol. 2023, 1–4. doi: 10.1038/s41423-023-00988-0.

34. Costello, S. M.; Shoemaker, S. R.; Hobbs, H. T.; Nguyen, A. W.; Hsieh, C.-L.; Maynard, J. A.; McLellan, J. S.; Pak, J. E.; Marqusee, S. The SARS-CoV-2 Spike Reversibly Samples an Open-Trimer Conformation Exposing Novel Epitopes. Nat Struct Mol Biol. 2022, 29, 229–238. doi: 10.1038/s41594-022-00735-5.

35. Calvaresi, V.; Wrobel, A. G.; Toporowska, J.; Hammerschmid, D.; Doores, K. J.; Bradshaw, R. T.; Parsons, R. B.; Benton, D. J.; Roustan, C.; Reading, E.; Malim, M. H.; Gamblin, S. J.; Politis, A. Structural Dynamics in the Evolution of SARS-CoV-2 Spike Glycoprotein. Nat Commun. 2023, 14, 1421. doi: 10.1038/s41467-023-36745-0.

36. Braet, S. M.; Buckley, T. S.; Venkatakrishnan, V.; Dam, K.-M. A.; Bjorkman, P. J.; Anand, G. S. Timeline of Changes in Spike Conformational Dynamics in Emergent SARS-CoV-2 Variants Reveal Progressive Stabilization of Trimer Stalk with Altered NTD Dynamics. Elife 2023, 12, e82584. doi: 10.7554/eLife.82584.

37. Raghuvamsi, P. V.; Tulsian, N. K.; Samsudin, F.; Qian, X.; Purushotorman, K.; Yue, G.; Kozma, M. M.; Hwa, W. Y.; Lescar, J.; Bond, P. J.; MacAry, P. A.; Anand, G. S. SARS-CoV-2 S Protein:ACE2 Interaction Reveals Novel Allosteric Targets. Elife 2021, 10, e63646. doi: 10.7554/eLife.63646.

38. Chen, C.; Zhu, R.; Hodge, E. A.; Díaz-Salinas, M. A.; Nguyen, A.; Munro, J. B.; Lee, K. K. hACE2-Induced Allosteric Activation in SARS-CoV versus SARS-CoV-2 Spike Assemblies Revealed by Structural Dynamics. ACS Infect Dis. 2023, 9, 1180–1189. doi: 10.1021/acsinfecdis.3c00010.

39. Toelzer, C.; Gupta, K.; Yadav, S. K. N.; Borucu, U.; Davidson, A. D.; Kavanagh Williamson, M.; Shoemark, D. K.; Garzoni, F.; Staufer, O.; Milligan, R.; Capin, J.; Mulholland, A. J.; Spatz, J.; Fitzgerald, D.; Berger, I.; Schaffitzel, C. Free Fatty Acid Binding Pocket in the Locked Structure of SARS-CoV-2 Spike Protein. Science 2020, 370, 725–730. doi: 10.1126/science.abd3255.

40. Toelzer, C.; Gupta, K.; Berger, I.; Schaffitzel, C. Cryo-EM Reveals Binding of Linoleic Acid to SARS-CoV-2 Spike Glycoprotein, Suggesting an Antiviral Treatment Strategy. Acta Crystallogr D Struct Biol. 2023, 79, 111–121. doi: 10.1107/S2059798323000049.

41. Toelzer, C.; Gupta, K.; Yadav, S. K. N.; Hodgson, L.; Williamson, M. K.; Buzas, D.; Borucu, U.; Powers, K.; Stenner, R.; Vasileiou, K.; Garzoni, F.; Fitzgerald, D.; Payré, C.; Gautam, G.; Lambeau, G.; Davidson, A. D.; Verkade, P.; Frank, M.; Berger, I.; Schaffitzel, C. The Free Fatty Acid–Binding Pocket Is a Conserved Hallmark in Pathogenic β-Coronavirus Spike Proteins from SARS-CoV to Omicron. Sci Adv. 2022, 8, eadc9179. doi: 10.1126/sciadv.adc9179.

42. Hao, A.; Song, W.; Li, C.; Zhang, X.; Tu, C.; Wang, X.; Wang, P.; Wu, Y.; Ying, T.; Sun, L. Defining a Highly Conserved Cryptic Epitope for Antibody Recognition of SARS-CoV-2 Variants. Signal Transduct Target Ther. 2023, 8, 269. doi: 10.1038/s41392-023-01534-0.

43. Bangaru, S.; Ozorowski, G.; Turner, H. L.; Antanasijevic, A.; Huang, D.; Wang, X.; Torres, J. L.; Diedrich, J. K.; Tian, J.-H.; Portnoff, A. D.; Patel, N.; Massare, M. J.; Yates, J. R., III; Nemazee, D.; Paulson, J. C.; Glenn, G.; Smith, G.; Ward, A. B. Structural Analysis of Full-Length SARS-CoV-2 Spike Protein from an Advanced Vaccine Candidate. Science 2020, 370, 1089–1094. doi: 10.1126/science.abe1502.

44. Rosa, A.; Pye, V. E.; Graham, C.; Muir, L.; Seow, J.; Ng, K. W.; Cook, N. J.; Rees-Spear, C.; Parker, E.; dos Santos, M. S.; Rosadas, C.; Susana, A.; Rhys, H.; Nans, A.; Masino, L.; Roustan, C.; Christodoulou, E.; Ulferts, R.; Wrobel, A. G.; Short, C.-E.; Fertleman, M.; Sanders, R. W.; Heaney, J.; Spyer, M.; Kjær, S.; Riddell, A.; Malim, M. H.; Beale, R.; MacRae, J. I.; Taylor, G. P.; Nastouli, E.; van Gils, M. J.; Rosenthal, P. B.; Pizzato, M.; McClure, M. O.; Tedder, R. S.; Kassiotis, G.; McCoy, L. E.; Doores, K. J.; Cherepanov, P. SARS-CoV-2 Can Recruit a Heme Metabolite to Evade Antibody Immunity. Sci Adv. 2021, 7, eabg7607. doi: 10.1126/sciadv.abg7607.

45. Altomare, C. G.; Adelsberg, D. C.; Carreno, J. M.; Sapse, I. A.; Amanat, F.; Ellebedy, A. H.; Simon, V.; Krammer, F.; Bajic, G. Structure of a Vaccine-Induced, Germline-Encoded Human Antibody Defines a Neutralizing Epitope on the SARS-CoV-2 Spike N-Terminal Domain. mBio 2022, 13, e0358021. doi: 10.1128/mbio.03580-21.

46. Silva, R. P.; Huang, Y.; Nguyen, A. W.; Hsieh, C.-L.; Olaluwoye, O. S.; Kaoud, T. S.; Wilen, R. E.; Qerqez, A. N.; Park, J.-G.; Khalil, A. M.; Azouz, L. R.; Le, K. C.; Bohanon, A. L.; DiVenere, A. M.; Liu, Y.; Lee, A. G.; Amengor, D. A.; Shoemaker, S. R.; Costello, S. M.; Padlan, E. A.; Marqusee, S.; Martinez-Sobrido, L.; Dalby, K. N.; D’Arcy, S.; McLellan, J. S.; Maynard, J. A. Identification of a Conserved S2 Epitope Present on Spike Proteins from All Highly Pathogenic Coronaviruses. Elife 2023, 12, e83710. doi: 10.7554/eLife.83710.

47. Zimmerman, M. I.; Porter, J. R.; Ward, M. D.; Singh, S.; Vithani, N.; Meller, A.; Mallimadugula, U. L.; Kuhn, C. E.; Borowsky, J. H.; Wiewiora, R. P., Hurley, M.F.D.; Harbison, A.M.; Fogarty, C.A.; Coffland, J.E.; Fadda, E.; Voelz, V.A.; Chodera, J.D.; Bowman, G.R. SARS-CoV-2 simulations go exascale to predict dramatic spike opening and cryptic pockets across the proteome. Nat. Chem. 2021, 13, 651–659. doi: 10.1038/s41557-021-00707-0.

48. Mori, T.; Jung, J.; Kobayashi, C.; Dokainish, H. M.; Re, S.; Sugita, Y. Elucidation of Interactions Regulating Conformational Stability and Dynamics of SARS-CoV-2 S-Protein. Biophys J. 2021, 120, 1060–1071. doi: 10.1016/j.bpj.2021.01.012.

49. Zuzic, L.; Samsudin, F.; Shivgan, A. T.; Raghuvamsi, P. V.; Marzinek, J. K.; Boags, A.; Pedebos, C.; Tulsian, N. K.; Warwicker, J.; MacAry, P.; Crispin, M.; Khalid, S.; Anand, G. S.; Bond, P. J. Uncovering Cryptic Pockets in the SARS-CoV-2 Spike Glycoprotein. Structure 2022, 30, 1062–1074.e4. doi: 10.1016/j.str.2022.05.006.

50. Ghoula, M.; Naceri, S.; Sitruk, S.; Flatters, D.; Moroy, G.; Camproux, A. C. Identifying Promising Druggable Binding Sites and Their Flexibility to Target the Receptor-Binding Domain of SARS-CoV-2 Spike Protein. Comput Struct Biotechnol J. 2023, 21, 2339–2351. doi: 10.1016/j.csbj.2023.03.029.

51. Verkhivker, G.; Agajanian, S.; Kassab, R.; Krishnan, K. Probing Mechanisms of Binding and Allostery in the SARS-CoV-2 Spike Omicron Variant Complexes with the Host Receptor: Revealing Functional Roles of the Binding Hotspots in Mediating Epistatic Effects and Communication with Allosteric Pockets. Int. J. Mol. Sci. 2022, 23, 11542, doi: 10.3390/ijms231911542.

52. Wang, Q.; Wang, L.; Zhang, Y.; Zhang, X.; Zhang, L.; Shang, W.; Bai, F. Probing the Allosteric Inhibition Mechanism of a Spike Protein Using Molecular Dynamics Simulations and Active Compound Identifications. J. Med. Chem. 2021, 65, 2827–2835, doi:10.1021/acs.jmedchem.1c00320.

53. Li, B.; Wang, L.; Ge, H.; Zhang, X.; Ren, P.; Guo, Y.; Chen, W.; Li, J.; Zhu, W.; Chen, W.; Zhu, L.; Bai, F. Identification of Potential Binding Sites of Sialic Acids on the RBD Domain of SARS-CoV-2 Spike Protein. Front. Chem. 2021, 9, 659764. doi: 10.3389/fchem.2021.659764.

54. Guo, L.; Lin, S.; Chen, Z.; Cao, Y.; He, B.; Lu, G. Targetable Elements in SARS-CoV-2 S2 Subunit for the Design of Pan-Coronavirus Fusion Inhibitors and Vaccines. Signal Transduct Target Ther. 2023, 8, 197. doi: 10.1038/s41392-023-01472-x.

55. Ma, J.; Su, D.; Sun, Y.; Huang, X.; Liang, Y.; Fang, L.; Ma, Y.; Li, W.; Liang, P.; Zheng, S. Cryo-Electron Microscopy Structure of S-Trimer, a Subunit Vaccine Candidate for COVID-19. J Virol. 2021, 95, e00194–21. doi: 10.1128/JVI.00194-21.

56. Huo, J.; Dijokaite-Guraliuc, A.; Liu, C.; Zhou, D.; Ginn, H. M.; Das, R.; Supasa, P.; Selvaraj, M.; Nutalai, R.; Tuekprakhon, A.; Duyvesteyn, H. M. E.; Mentzer, A. J.; Skelly, D.; Ritter, T. G.; Amini, A.; Bibi, S.; Adele, S.; Johnson, S. A.; Paterson, N. G.; Williams, M. A.; Hall, D. R.; Plowright, M.; Newman, T. A. H.; Hornsby, H.; de Silva, T. I.; Temperton, N.; Klenerman, P.; Barnes, E.; Dunachie, S. J.; Pollard, A. J.; Lambe, T.; Goulder, P.; Fry, E. E.; Mongkolsapaya, J.; Ren, J.; Stuart, D. I.; Screaton, G. R. A Delicate Balance between Antibody Evasion and ACE2 Affinity for Omicron BA.2.75. Cell Rep. 2023, 42, 111903. doi: 10.1016/j.celrep.2022.111903.

57. Rose, P. W.; Prlic, A.; Altunkaya, A.; Bi, C.; Bradley, A. R.; Christie, C. H.; Costanzo, L. D.; Duarte, J. M.; Dutta, S.; Feng, Z.; Green, R. K.; Goodsell, D. S.; Hudson, B.; Kalro, T.; Lowe, R.; Peisach, E.; Randle, C.; Rose, A. S.; Shao, C.; Tao, Y. P.; Valasatava, Y.; Voigt, M.; Westbrook, J. D.; Woo, J.; Yang, H.; Young, J. Y.; Zardecki, C.; Berman, H. M.; Burley, S. K. The RCSB protein data bank: integrative view of protein, gene and 3D structural information. Nucleic Acids Res. 2017, 45, D271–D281. doi: 10.1093/nar/gkw1000.

58. Hekkelman, M.L.; Te Beek, T.A.; Pettifer, S.R.; Thorne, D.; Attwood, T.K.; Vriend, G. WIWS: A protein structure bioinformatics web service collection. Nucleic Acids Res. 2010, 38, W719–W723. doi: 10.1093/nar/gkq453.

59. Fernandez-Fuentes, N.; Zhai, J.; Fiser, A. ArchPRED: A template based loop structure prediction server. Nucleic Acids Res. 2006, 34, W173–W176. doi: 10.1093/nar/gkl113.

60. Krivov, G.G.; Shapovalov, M.V.; Dunbrack, R.L., Jr. Improved prediction of protein side-chain conformations with SCWRL4. Proteins 2009, 77, 778–795. doi: 10.1002/prot.22488.

61. Søndergaard C. R.; Olsson M. H.; Rostkowski M.; Jensen J. H. Improved treatment of ligands and coupling effects in empirical calculation and rationalization of pKa values. J. Chem. Theory Comput. 2011, 7, 2284–2295. 10.1021/ct200133y.

62. Olsson M. H.; Søndergaard C. R.; Rostkowski M.; Jensen J. H. PROPKA3: consistent treatment of internal and surface residues in empirical pKa predictions. J. Chem. Theory Comput. 2011, 7, 525–537. 10.1021/ct100578z.

63. Bhattacharya, D.; Nowotny, J.; Cao, R.; Cheng, J. 3Drefine: An Interactive Web Server for Efficient Protein Structure Refinement. Nucleic Acids Res. 2016, 44, W406–W409. doi: 10.1093/nar/gkw336.

64. Huang, J.; Rauscher, S.; Nawrocki, G.; Ran, T.; Feig, M.; de Groot, B.L.; Grubmüller, H.; MacKerell, A.D. Jr. CHARMM36m: an improved force field for folded and intrinsically disordered proteins. Nat. Methods 2017, 14, 71–73. doi: 10.1038/nmeth.4067.

65. Jorgensen, W. L.; Chandrasekhar, J.; Madura, J. D.; Impey, R. W.; Klein, M. L. Comparison of Simple Potential Functions for Simulating Liquid Water. J. Chem. Phys. 1983, 79, 926– 935. 10.1063/1.445869.

66. Ryckaert, J.-P.; Ciccotti, G.; Berendsen, H. J. C. Numerical Integration of the Cartesian Equations of Motion of a System with Constraints: Molecular Dynamics of n-Alkanes. J. Comput. Phys. 1977, 23, 327–341. 10.1016/0021-9991(77)90098-5.

67. Di Pierro, M.; Elber, R.; Leimkuhler, B. A Stochastic Algorithm for the Isobaric-Isothermal Ensemble with Ewald Summations for All Long Range Forces. J. Chem. Theory Comput. 2015, 11, 5624–5637. 10.1021/acs.jctc.5b00648.

68. Eastman, P.; Swails, J.; Chodera, J. D.; McGibbon, R. T.; Zhao, Y.; Beauchamp, K. A.; Wang, L.-P.; Simmonett, A. C.; Harrigan, M. P.; Stern, C. D.; Wiewiora, R. P.; Brooks, B. R.; Pande, V. S. OpenMM 7: Rapid Development of High Performance Algorithms for Molecular Dynamics. PLoS Comput. Biol. 2017, 13, e1005659. doi: 10.1371/journal.pcbi.1005659.

69. Childers, M. C.; Daggett, V. Validating Molecular Dynamics Simulations against Experimental Observables in Light of Underlying Conformational Ensembles. J. Phys. Chem. B. 2018, 122, 6673–6689. doi: 10.1021/acs.jpcb.8b02144.

70. McGibbon, R. T.; Beauchamp, K. A.; Harrigan, M. P.; Klein, C.; Swails, J. M.; Hernández, C. X.; Schwantes, C. R.; Wang, L.-P.; Lane, T. J.; Pande, V. S. MDTraj: A Modern Open Library for the Analysis of Molecular Dynamics Trajectories. Biophys. J. 2015, 109, 1528– 1532. doi: 10.1016/j.bpj.2015.08.015.

71. Haque, I. S.; Beauchamp, K. A.; Pande, V. S. A Fast 3 x N Matrix Multiply Routine for Calculation of Protein RMSD, bioRxiv 2014, 10.1101/008631.

72. Theobald D.L. Rapid Calculation of RMSDs Using a Quaternion-Based Characteristic Polynomial. Acta Crystallogr. A. 2005, 61, 478–480.

73. Liu, P.; Agrafiotis, D. K.; Theobald, D. L. Fast Determination of the Optimal Rotational Matrix for Macromolecular Superpositions. J. Comput. Chem. 2010, 31, 1561–1563. doi: 10.1002/jcc.21439.

74. Naritomi, Y.; Fuchigami, S. Slow Dynamics in Protein Fluctuations Revealed by Time-Structure Based Independent Component Analysis: The Case of Domain Motions. J. Chem. Phys. 2011, 134, 065101. doi: 10.1063/1.3554380.

75. Schwantes, C. R.; Pande, V. S. Improvements in Markov State Model Construction Reveal Many Non-Native Interactions in the Folding of NTL9. J. Chem. Theory Comput. 2013, 9, 2000–2009. doi: 10.1021/ct300878a.

76. M. Sultan, M.; Pande, V. S. tICA-Metadynamics: Accelerating Metadynamics by Using Kinetically Selected Collective Variables. J. Chem. Theory Comput. 2017, 13, 2440–2447. doi: 10.1021/acs.jctc.7b00182.

77. Trozzi, F.; Wang, X.; Tao, P. UMAP as a Dimensionality Reduction Tool for Molecular Dynamics Simulations of Biomacromolecules: A Comparison Study. J. Phys. Chem. B. 2021, 125, 5022–5034. doi: 10.1021/acs.jpcb.1c02081.

78. Scherer, M. K.; Trendelkamp-Schroer, B.; Paul, F.; Pérez-Hernández, G.; Hoffmann, M.; Plattner, N.; Wehmeyer, C.; Prinz, J.-H.; Noé, F. PyEMMA 2: A Software Package for Estimation, Validation, and Analysis of Markov Models. J. Chem. Theory Comput. 2015, 11, 5525–5542. doi: 10.1021/acs.jctc.5b00743.

79. Wu, H.; Paul, F.; Wehmeyer, C.; Noe, F., Multiensemble Markov models of molecular thermodynamics and kinetics. Proc. Natl. Acad. Sci. U.S.A. 2016, 113 (23), E3221–E3230, doi: 10.1073/pnas.1525092113.

80. Suárez, E.; Adelman, J. L.; Zuckerman, D. M. Accurate Estimation of Protein Folding and Unfolding Times: Beyond Markov State Models. J. Chem. Theory Comput. 2016, 12, 3473–3481. doi: 10.1021/acs.jctc.6b00339.

81. Bowman, G. R.; Bolin, E. R.; Hart, K. M.; Maguire, B. C.; Marqusee, S., Discovery of multiple hidden allosteric sites by combining Markov state models and experiments. Proc. Natl. Acad. Sci. U.S.A. 2015, 112 (9), 2734–2739, doi: 10.1073/pnas.1417811112.

82. Bowman, G. R.; Noé, F. Software for Building Markov State Models. Adv. Exp. Med. Biol. 2014, 797, 139. doi: 10.1007/978-94-007-7606-7_11.

83. Bowman, G. R. A Tutorial on Building Markov State Models with MSMBuilder and Coarse-Graining Them with BACE. Methods Mol Biol. 2014, 1084, 141–58. doi: 10.1007/978-1-62703-658-0_8.

84. Trendelkamp-Schroer, B.; Wu, H.; Paul, F.; Noé, F. Estimation and Uncertainty of Reversible Markov Models. J. Chem. Phys. 2015, 143, 174101. doi: 10.1063/1.4934536.

85. Bowman, G. R.; Huang, X.; Pande, V. S. Using Generalized Ensemble Simulations and Markov State Models to Identify Conformational States. Methods 2009, 49, 197–201. doi: 10.1016/j.ymeth.2009.04.013.

86. Sacquin-Mora, S.; Lavery, R., Investigating the local flexibility of functional residues in hemoproteins. Biophys. J. 2006, 90, 2706–2717.

87. Sacquin-Mora, S.; Laforet, E.; Lavery, R., Locating the active sites of enzymes using mechanical properties. Proteins 2007, 67, 350–359.

88. Sacquin-Mora S. Bridging Enzymatic Structure Function via Mechanics: A Coarse-Grain Approach. Methods Enzymol. 2016, 578, 227–248. doi: 10.1016/bs.mie.2016.05.022.

89. Ermak, D.L.; McCammon, J.A. Brownian dynamics with hydrodynamic interactions. J. Chem. Phys. 1978, 69, 1352–1360.

90. Pastor, R.W.; Venable, R.; Karplus, M. Brownian dynamics simulation of a lipid chain in a membrane bilayer. J. Chem. Phys. 1988, 89, 1112–1127.

91. Rotkiewicz, P.; Skolnick, J. Fast procedure for reconstruction of full-atom protein models from reduced representations. J. Comput. Chem. 2008, 29, 1460–1465.

92. Lombardi, L. E.; Marti, M. A.; Capece, L. CG2AA: backmapping protein coarse-grained structures. Bioinformatics 2016, 32, 1235–1237.

93. Dehouck, Y.; Kwasigroch, J. M.; Rooman, M.; Gilis, D. BeAtMuSiC: Prediction of changes in protein-protein binding affinity on mutations. Nucleic Acids Res. 2013, 41, W333–W339. doi: 10.1093/nar/gkt450.

94. Dehouck, Y.; Gilis, D.; Rooman, M. A new generation of statistical potentials for proteins. Biophys. J. 2006, 90, 4010–4017. doi: 10.1529/biophysj.105.079434.

95. Dehouck, Y.; Grosfils, A.; Folch, B.; Gilis, D.; Bogaerts, P.; Rooman, M. Fast and accurate predictions of protein stability changes upon mutations using statistical potentials and neural networks: PoPMuSiC-2.0. Bioinformatics 2009, 25, 2537–2543. doi: 10.1093/bioinformatics/btp445.

96. Krivák, R.; Hoksza, D. P2Rank: Machine Learning Based Tool for Rapid and Accurate Prediction of Ligand Binding Sites from Protein Structure. J. Cheminform. 2018, 10, 39. doi: 10.1186/s13321-018-0285-8.

97. Jakubec, D.; Skoda, P.; Krivak, R.; Novotny, M.; Hoksza, D. PrankWeb 3: Accelerated Ligand-Binding Site Predictions for Experimental and Modelled Protein Structures. Nucleic Acids Res. 2022, 50, W593–W597. doi: 10.1093/nar/gkac389.

98. Xiao, S., Tian, H., & Tao, P. PASSer2.0: Accurate Prediction of Protein Allosteric Sites Through Automated Machine Learning. Front. Mol. Biosci. 2022, 9, 879251. doi: 10.3389/fmolb.2022.879251.

99. Tian, H.; Xiao, S.; Jiang, X.; Tao, P. PASSer: Fast and Accurate Prediction of Protein Allosteric Sites. Nucleic Acids Res. 2023, 51, W427–W431. doi: 10.1093/nar/gkad303. doi: 10.1093/nar/gkad303.

100. Tian, H.; Xiao, S.; Jiang, X.; Tao, P. PASSerRank: Prediction of Allosteric Sites with Learning to Rank. J. Comput. Chem. 2023 Aug 10. doi: 10.1002/jcc.27193.

101. Verkhivker, G.; Alshahrani, M.; Gupta, G. Balancing Functional Tradeoffs between Protein Stability and ACE2 Binding in the SARS-CoV-2 Omicron BA.2, BA.2.75 and XBB Lineages: Dynamics-Based Network Models Reveal Epistatic Effects Modulating Compensatory Dynamic and Energetic Changes. Viruses 2023, 15, 1143. doi: 10.3390/v15051143.

102. Starr, T. N.; Greaney, A. J.; Hilton, S. K.; Ellis, D.; Crawford, K. H. D.; Dingens, A. S.; Navarro, M. J.; Bowen, J. E.; Tortorici, M. A.; Walls, A. C.; King, N. P.; Veesler, D.; Bloom, J. D. Deep Mutational Scanning of SARS-CoV-2 Receptor Binding Domain Reveals Constraints on Folding and ACE2 Binding. Cell 2020, 182, 1295–1310.e20. doi: 10.1016/j.cell.2020.08.012.

103. Starr, T. N.; Greaney, A. J.; Stewart, C. M.; Walls, A. C.; Hannon, W. W.; Veesler, D.; Bloom, J. D. Deep Mutational Scans for ACE2 Binding, RBD Expression, and Antibody Escape in the SARS-CoV-2 Omicron BA.1 and BA.2 Receptor-Binding Domains. PLoS Pathog. 2022, 18, e1010951. doi: 10.1371/journal.ppat.1010951.

104. Wang, Q.; Iketani, S.; Li, Z.; Guo, Y.; Yeh, A. Y.; Liu, M.; Yu, J.; Sheng, Z.; Huang, Y.; Liu, L.; Ho, D. D. Antigenic Characterization of the SARS-CoV-2 Omicron Subvariant BA.2.75. Cell Host Microbe 2022, 30, 1512–1517.e4. doi: 10.1016/j.chom.2022.09.002.

105. Zhao, Z.; Xie, Y.; Bai, B.; Luo, C.; Zhou, J.; Li, W.; Meng, Y.; Li, L.; Li, D.; Li, X.; Li, X.; Wang, X.; Sun, J.; Xu, Z.; Sun, Y.; Zhang, W.; Fan, Z.; Zhao, X.; Wu, L.; Ma, J.; Li, O. Y.; Shang, G.; Chai, Y.; Liu, K.; Wang, P.; Gao, G. F.; Qi, J. Structural Basis for Receptor Binding and Broader Interspecies Receptor Recognition of Currently Circulating Omicron Sub-Variants. Nat Commun. 2023,14, 4405. doi: 10.1038/s41467-023-39942-z.

106. Qing, E.; Li, P.; Cooper, L.; Schulz, S.; Jäck, H.-M.; Rong, L.; Perlman, S.; Gallagher, T. Inter-Domain Communication in SARS-CoV-2 Spike Proteins Controls Protease-Triggered Cell Entry. Cell Rep. 2022, 39, 110786. doi: 10.1016/j.celrep.2022.110786.

107. Verkhivker, G. M.; Di Paola, L. Dynamic Network Modeling of Allosteric Interactions and Communication Pathways in the SARS-CoV-2 Spike Trimer Mutants: Differential Modulation of Conformational Landscapes and Signal Transmission via Cascades of Regulatory Switches. J. Phys. Chem. B. 2021, 125, 850–873, doi: 10.1021/acs.jpcb.0c10637.

108. Verkhivker, G.M.; Di Paola, L. Integrated Biophysical Modeling of the SARS-CoV-2 Spike Protein Binding and Allosteric Interactions with Antibodies. J. Phys. Chem. B. 2021, 125, 4596–4619, DOI: 10.1021/acs.jpcb.1c00395.

109. Verkhivker, G.M.; Agajanian, S.; Oztas, D.Y.; Gupta, G. Dynamic Profiling of Binding and Allosteric Propensities of the SARS-CoV-2 Spike Protein with Different Classes of Antibodies: Mutational and Perturbation-Based Scanning Reveals the Allosteric Duality of Functionally Adaptable Hotspots. J. Chem. Theory Comput. 2021, 17, 4578–4598, doi: 10.1021/acs.jctc.1c00372.

110. Verkhivker, G.M.; Agajanian, S.; Oztas, D.Y.; Gupta, G. Comparative Perturbation-Based Modeling of the SARS-CoV-2 Spike Protein Binding with Host Receptor and Neutralizing Antibodies: Structurally Adaptable Allosteric Communication Hotspots Define Spike Sites Targeted by Global Circulating Mutations. Biochemistry 2021, 60, 1459–1484, doi: 10.1021/acs.biochem.1c00139.

