## Supplementary material for "Examining Functional Linkages Between Conformational Dynamics, Protein Stability and Evolution of Cryptic Binding Pockets in the SARS-CoV-2 Omicron Spike Complexes with the ACE2 Host Receptor: Recombinant Omicron Variants Mediate Variability of Conserved Allosteric Sites and Binding Epitopes": Supprting Figures S1-S8 and Tables S1-S6: SUPPLEMENTARY_MATERIALS.docx

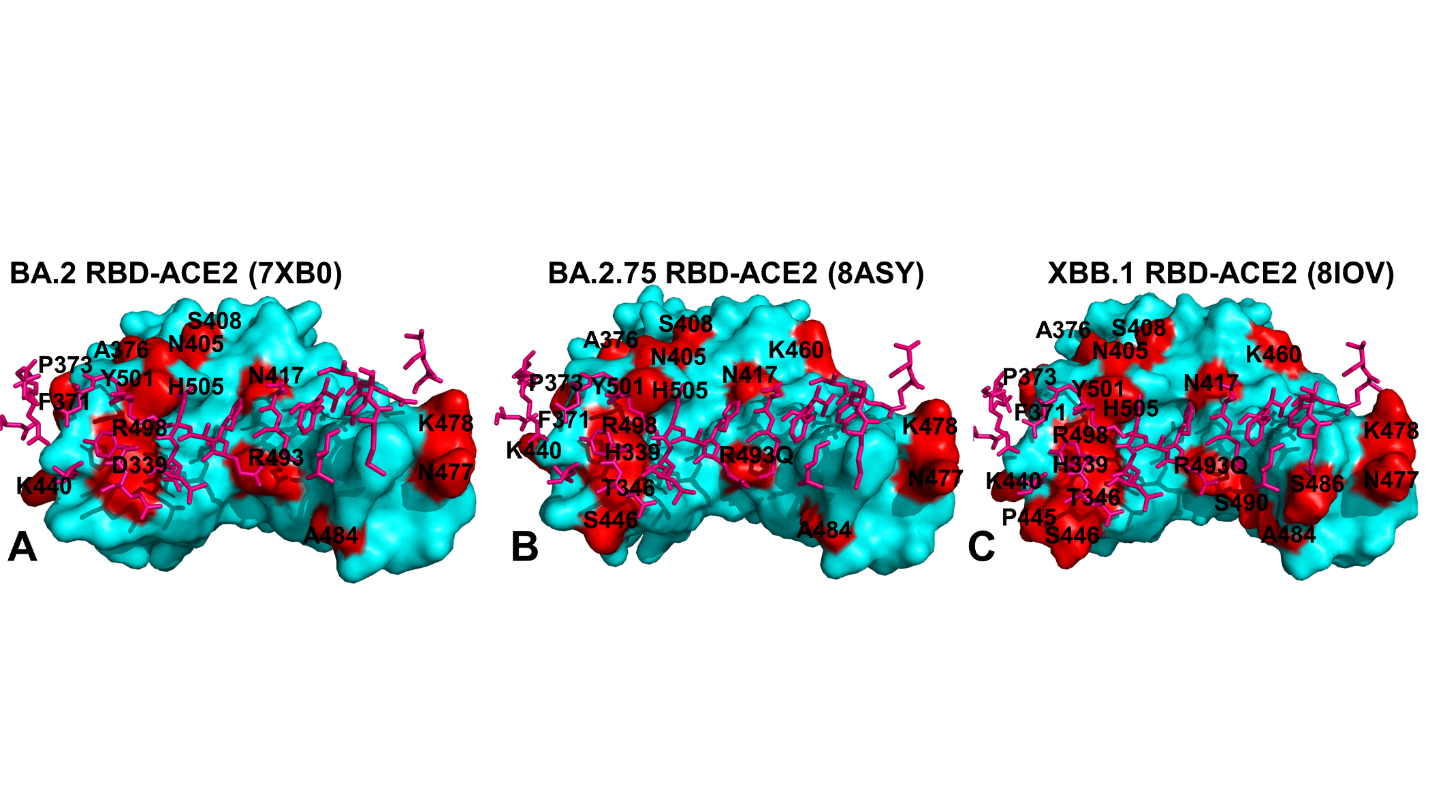

**Figure S1.** Structural organization of the SARS-CoV-2-RBD Omicron BA.2, BA.2.75, and XBB.1. binding interfaces in the complexes with human ACE2 enzyme. (A) The RBD-BA.2 is shown in cyan surface from the top view. The ACE2 binding residues are shown in pink sticks. The Omicron RBD BA.2 sites (G339D, S371F, S373P, S375F, T376A, D405N, R408S, K417N, N440K, S477N, T478K, E484A, Q493R, Q498R, N501Y, Y505H) are shown in red surface and annotated. (B) The RBD-BA.2.75 is shown from the top view. The ACE2 binding residues are shown in pink sticks. The Omicron RBD BA.2.75 sites (G339H, S371F, S373P, S375F, T376A, D405N, R408S, K417N, N440K, G446N, N460K, S477N, T478K, E484A, R493Q, Q498R, N501Y, Y505H) are shown in red surface and annotated. (C) The RBD-XBB.1 is shown from the top view and the ACE2 binding residues are shown in pink-colored sticks. The Omicron RBD XBB.1 sites (G339H, R346T, L368I, S371F, S373P, S375F, T376A, D405N, R408S, K417N, N440K, V445P, G446S,N460K, S477N, T478K, E484A, F486S, F490S, R493Q, Q498R, N501Y, Y505H) are shown in red-colored surface and annotated.

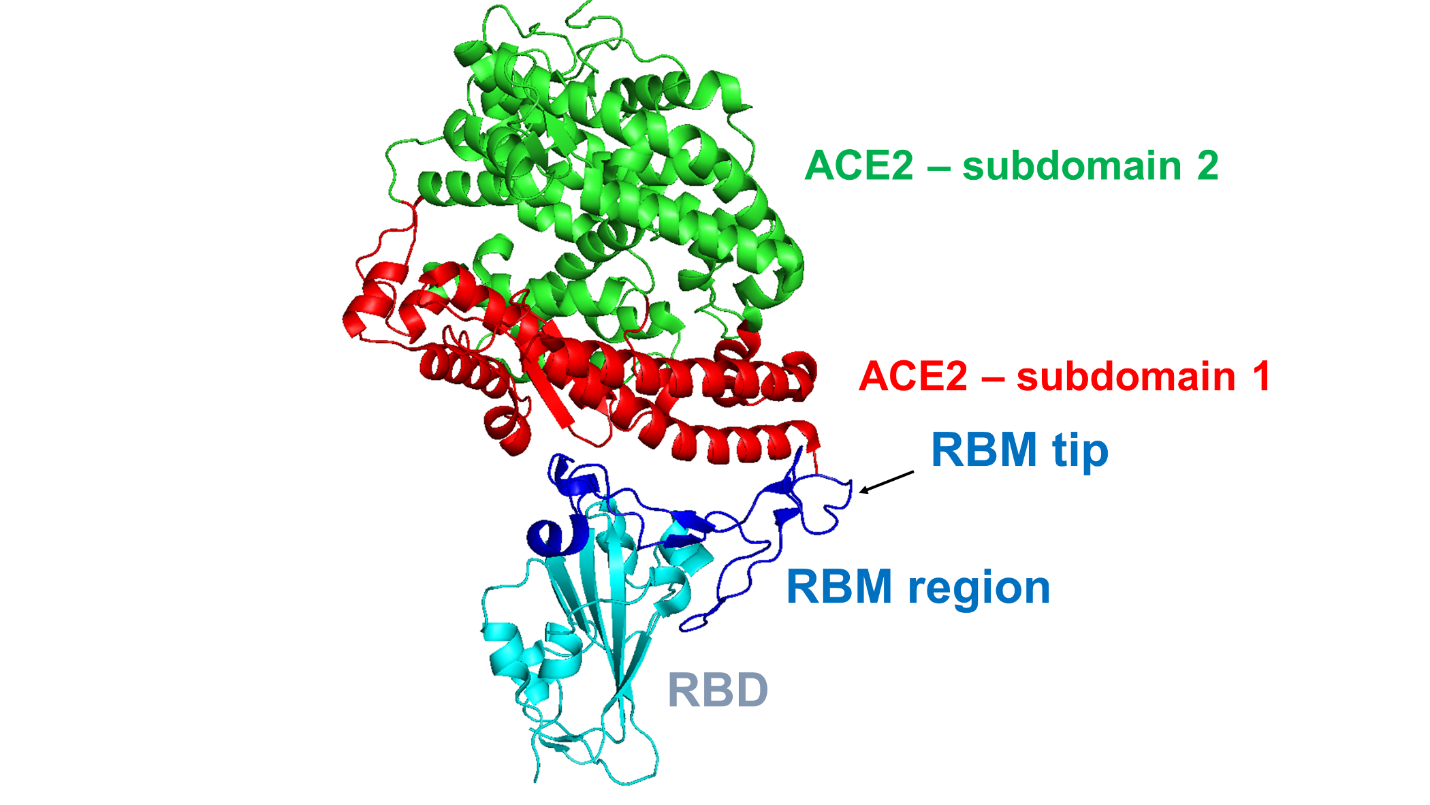

**Figure S2.** A general overview of the secondary structure elements and binding interface in the SARS-CoV-2 RBD complex with human ACE2. The RBD is shown in cyan and secondary structure elements are annotated. The RBM region is in blue ribbons and annotated. The subdomains I and II of human ACE2 are shown in red and green ribbons respectively.

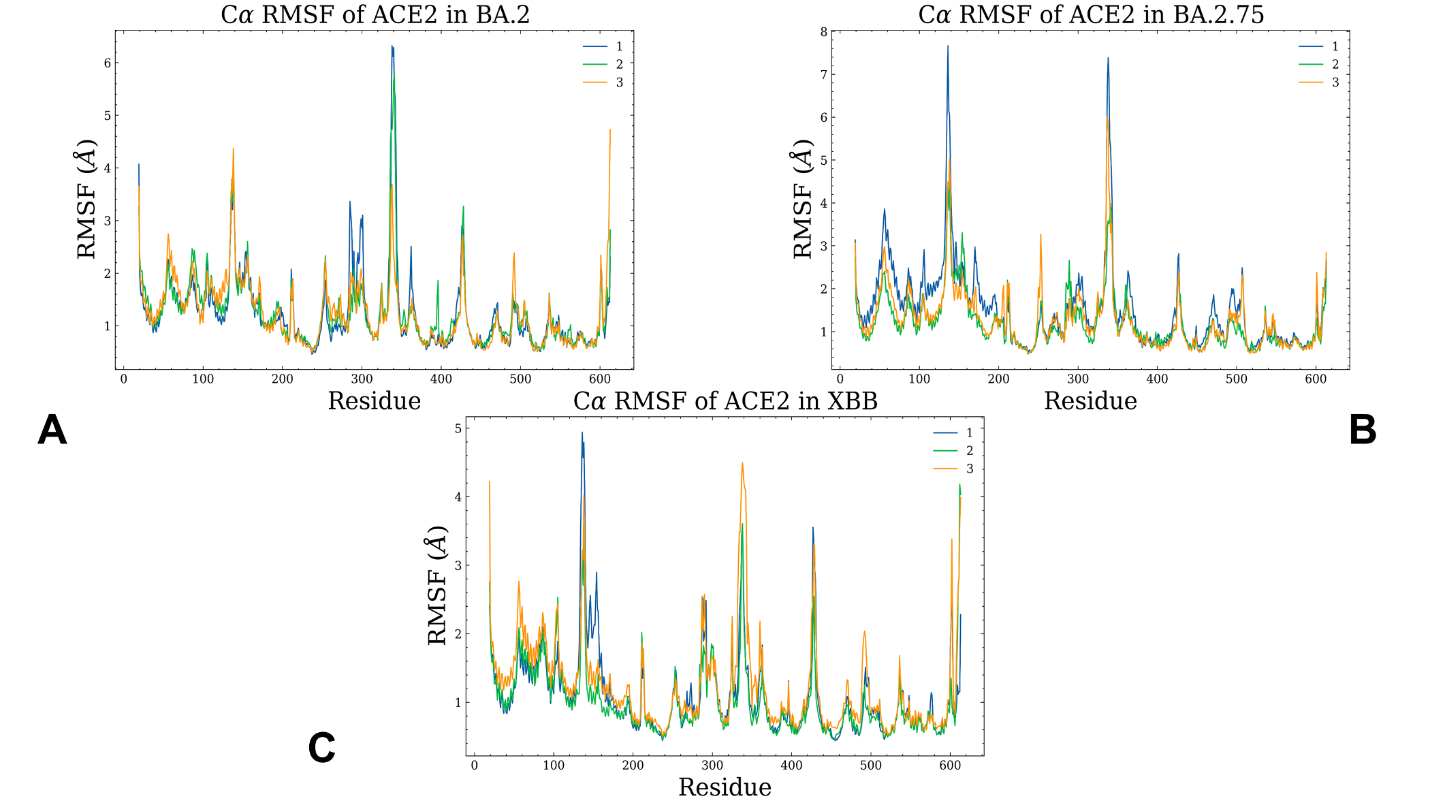

**Figure S3.** Conformational dynamics profiles of the ACE2 residues obtained from three independent MD simulations of the Omicron RBD BA.2, BA.2.75 and XBB.1 complexes with ACE2. The RMSF profiles for the ACE2 backbone residues obtained from three independent microsecond MD simulations of the Omicron BA.2 RBD-ACE2 (A), Omicron BA.2.75 RBD-ACE2 (B), Omicron XBB.1 RBD-ACE2 (C).

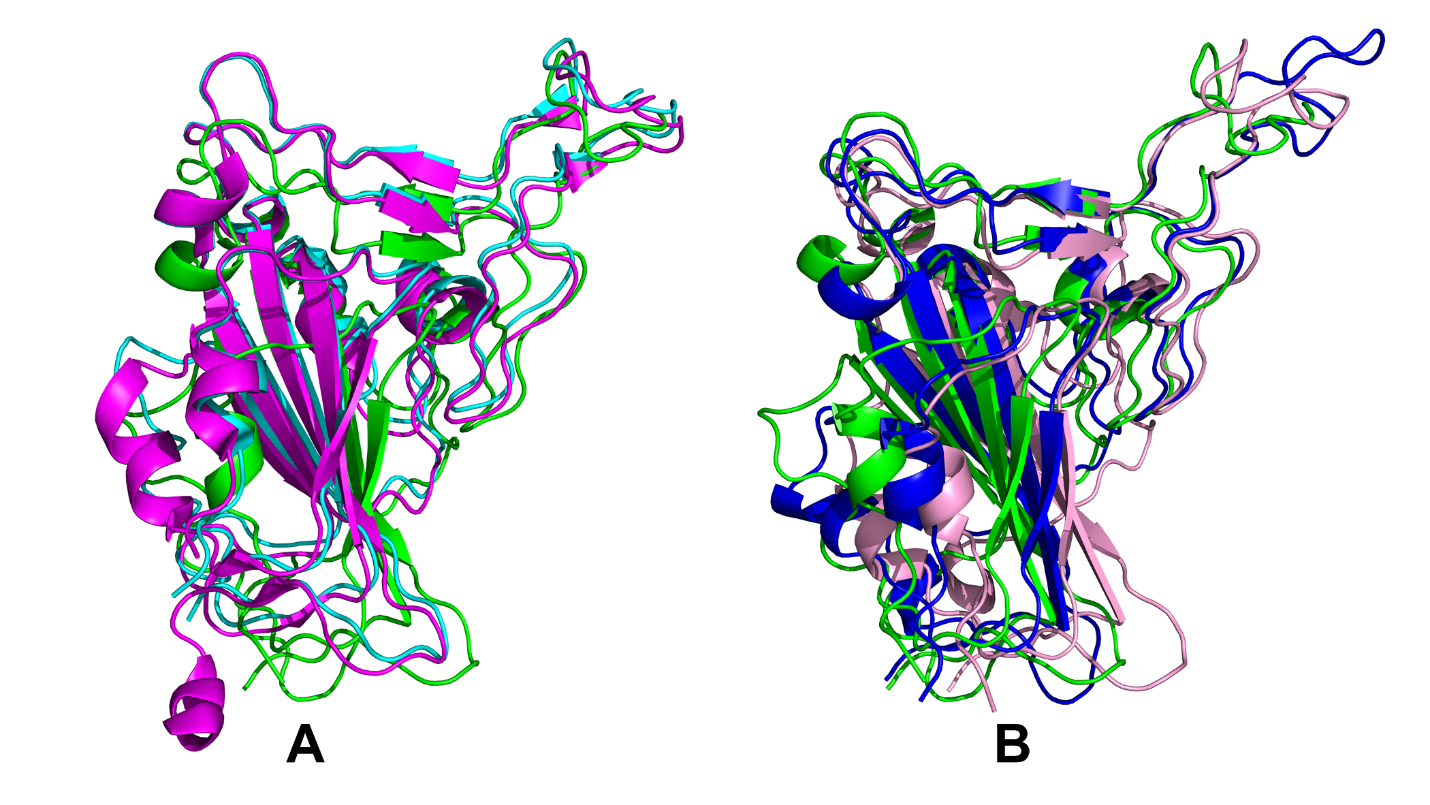

**Figure S4.** Structural analysis of the macrostates obtained from atomistic simulations of Omicron BA.2, BA.2.75 ad XBB.1 complexes with ACE2. (A) Structural alignment of the crystal structures of the Omicron BA.2 RBD (in cyan), BA.2.75 RBD (in magenta) and RBD conformation of the macrostate 9. (B) Structural alignment of RBD conformations for macrostate 9 (in green), macrostate 8 (in pink) and macrostate 1 (in blue). The RBD structures are shown in ribbons. For clarity of the presentation the ACE2 molecules that are very similar in all structures and identified macrostates are omitted.

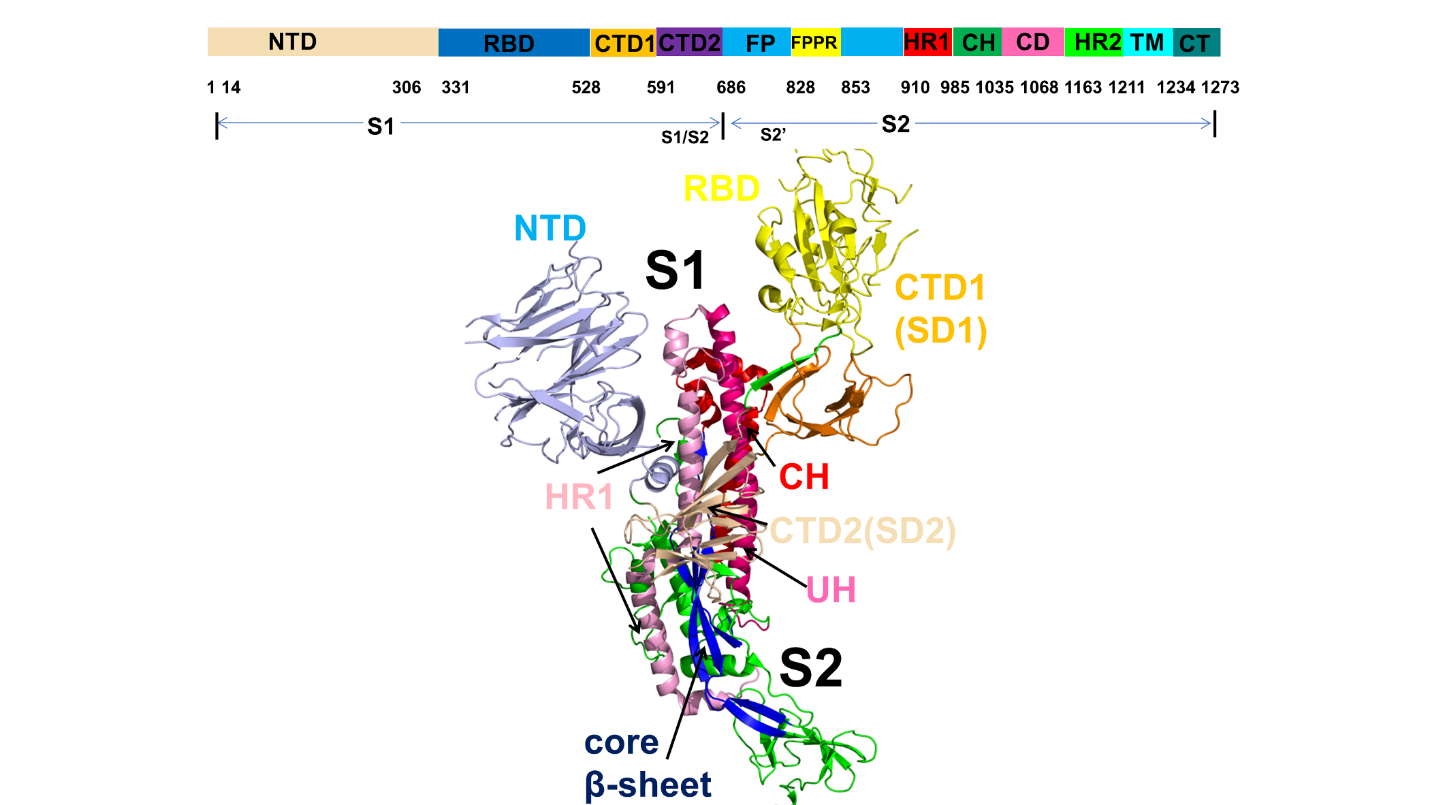

**Figure S5.** A schematic representation of domain organization and residue range for the full-length SARS-CoV-2 S protein. The subunits S1 and S2 include NTD RBD, C-terminal domain 1(CTD1), C-terminal domain 2 (CTD2), S1/S2 cleavage site (S1/S2), S2’ cleavage site (S2’), fusion peptide (FP), fusion peptide proximal region (FPPR), heptad repeat 1 (HR1), central helix region (CH), connector domain (CD), heptad repeat 2 (HR2), transmembrane domain (TM), and cytoplasmic tail (CT).The subunits S1 regions are annotated as follows : NTD (residues 14-306) in light blue; RBD (residues 331-528) in yellow; CTD1 (residues 528-591) in orange; CTD2 (residues 592-686) in wheat color ; upstream helix (UH) (residues 736-781) in red; HR1 (residues 910-985) in pink; CH (residues 986-1035) in hot pink; core β-sheet (residues 711-736, 1045-1076) (in blue).

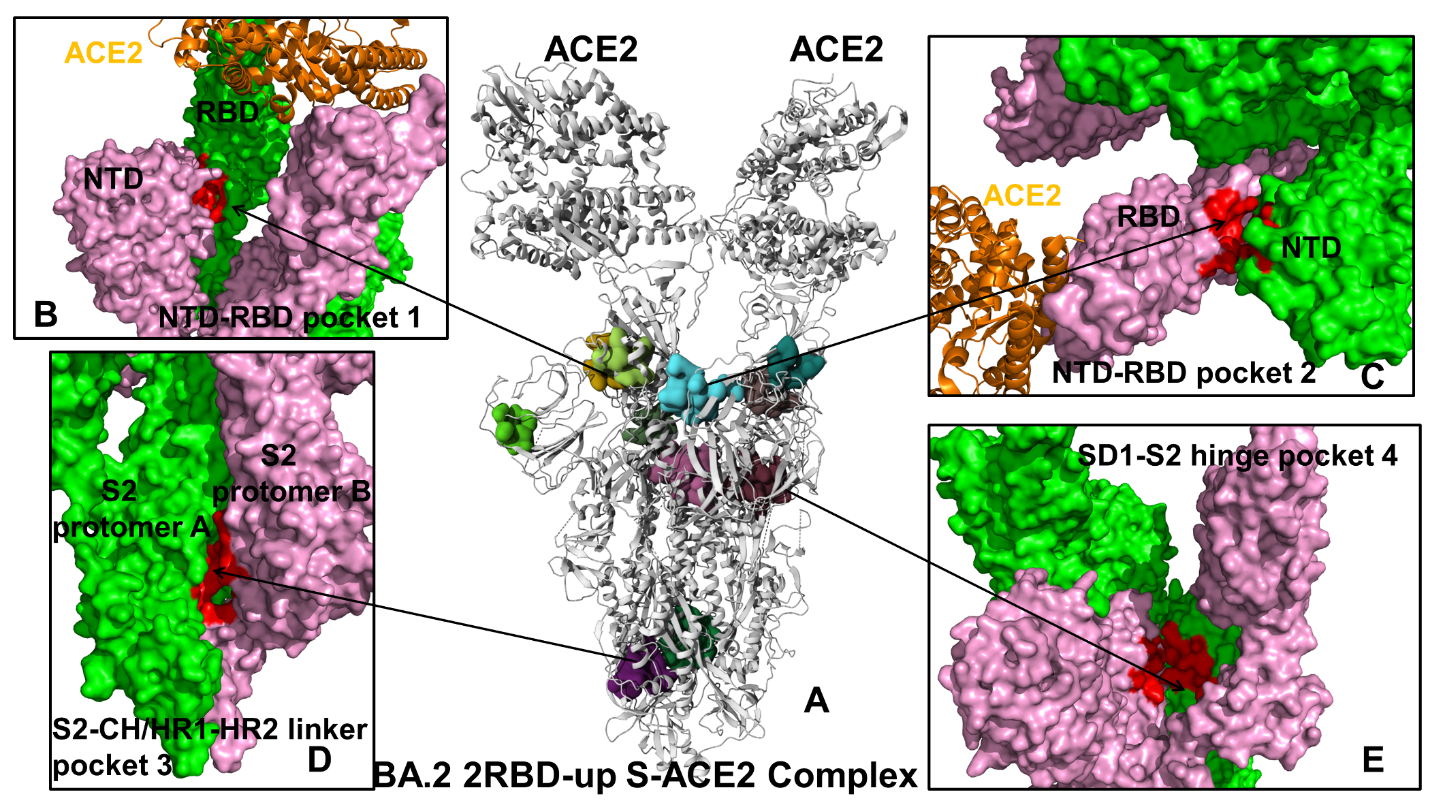

**Figure S6.** Structural map and residue-based close-ups of the top ranked cryptic sites for the ensemble of the S-BA.2 2RBD-up trimer complex with ACE2 . (A) The BA.2 2RBD-up trimer complex (pdb id 7XO7) is shown in ribbons and the top ranked allosteric sites are in surface. (B) Pocket 1 (NTD-RBD inter-protomer pocket, protomer A RBD residues A_357, A_359, A_360, A_393, A_394, A_520, A_521, A_523, and protomer C NTD residues C_115, C_130, C_168, C_230, C_231). (C) Pocket 2 ( NTD-RBD interprotomer pocket, protomer B NTD residues B_115, B_132, B_167, B_168, B_170, B_230, B_231, B_232, and protomer C RBD residues C_357, C_393, C_394, C_518, C_520, C_521, C_523). (D) Pocket 3 (inter-protomer pocket in S2, protomer A residues A_1034, A_1035, A_1036, A_886, A_904, A_908, and protomer B residues B_1038, B_1039, B_1040, B_1046, B_1047, B_1107, B_1108, B_909). (E) Pocket 4 (inter-protomer pocket in S2, protomer A residues A_1002, A_970, A_995, A_998, A_999, and protomer B residues B_1001, B_1002, B_756, B_759, B_994, B_995, B_998, C_1002, C_756, C_970, C_994, C_995, C_998, C_999). The predicted pockets are shown in red surface.

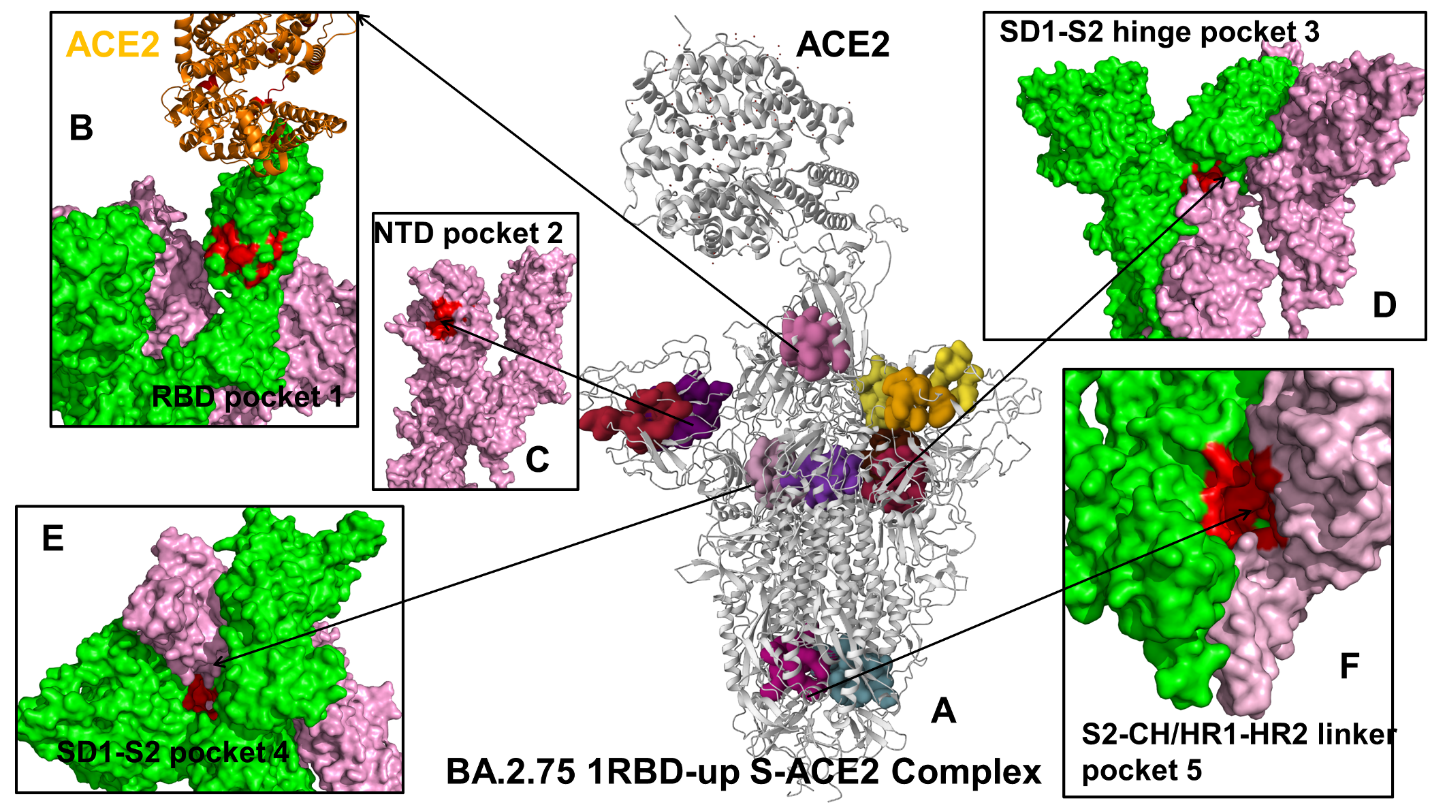
**Figure S7.** Structural map and residue-based close-ups of the top ranked cryptic sites for the ensemble of the S-BA.2.75 1RBD-up trimer complex with ACE2 . (A) The BA.2.75 1RBD-up trimer complex (pdb id 7YR2) is shown in ribbons and the top ranked allosteric sites are in surface. (B) Pocket 1 (RBD pocket residues on RBD-up protomer D: D_336, D_338, D_342, D_364, D_365, D_367, D_368, D_369, D_371, D_377, D_379, D_382, D_383, D_384, D_387, D_432, D_434, D_513, D_515). (C) Pocket 2 ( NTD pocket residues on RBD-up protomer D: D_100, D_101, D_239, D_240, D_242, D_248, D_249, D_250, D_263, D_264, D_265, D_65, D_66, D_81, D_84, D_94, D_95, D_96). (D) Pocket 3 (inter-protomer SD1-S2 hinge pocket. Protomer C residues C_541, C_546, C_547, C_548, C_549, C_570, C_572, C_573, C_587, C_589, C_592, and protomer D residues D_1000, D_740, D_741, D_744, D_745, D_855, D_856, D_966, D_976, D_977, D_978). (E) Pocket 4 (inter-protomer SD1-S2 pocket. Protomer C residues C_974, C_976, C_979 and protomer E residues E_391, E_517, E_518, E_519, E_520, E_522, E_544, E_545, E_546, E_565, E_567, E_571, E_573). (F) Pocket 5 (inter-protomer pocket in S2 CH/HR1-HR2. Protomer C residues C_1035, C_1036, C_1038, C_884, C_885, C_886, C_887, C_896, C_904, C_907, C_908, and protomer E residues E_1038, E_1040, E_1047, E_1107, E_1108, E_712, E_909, E_910). On panels (B-F) the S protein protomers are shown in green and pink surfaces. For clarity only two chains out of three protomers are highlighted. The predicted pockets are shown in red surface.

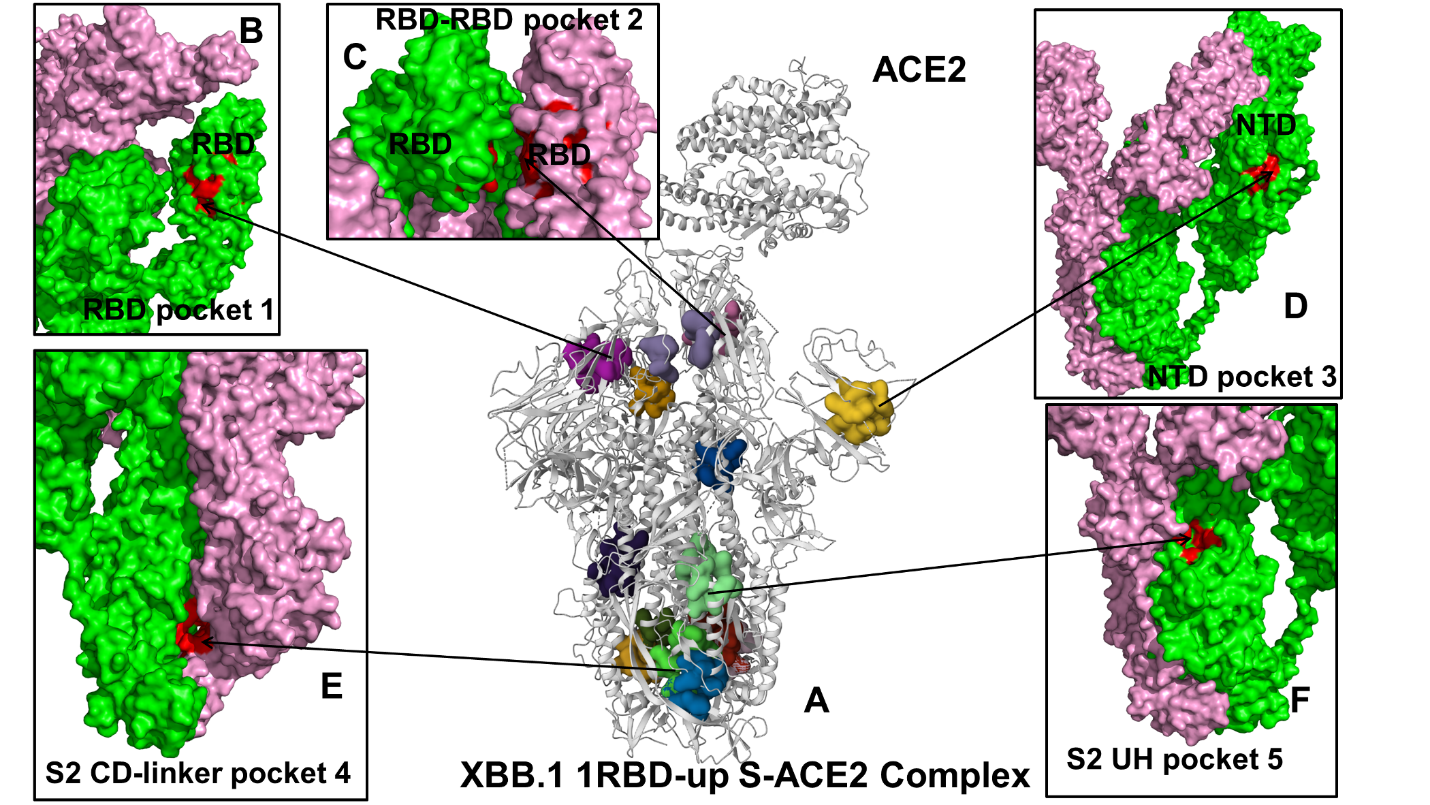

**Figure S8.** Structural map and residue-based close-ups of the top ranked cryptic sites for the ensemble of the S-XBB.1 1RBD-up trimer complex with ACE2 . (A) The XBB.1 1RBD-up trimer complex (pdb id 8IOU) is shown in ribbons and the top ranked allosteric sites are in surface. (B) Pocket 1 (RBD pocket residues on protomer C : C_338, C_365, C_368, C_369, C_377, C_387, C_432, C_434, C_513). (C) Pocket 2 (RBD-RBD pocket. RBD residues on protomer B B_369, B_372, B_373, B_374, B_377, and RBD residues on protomer C C_420, C_421, C_460). (D) Pocket 3 (NTD pocket on RBD-up protomer A : A_101, A_102, A_103, A_104, A_121, A_175, A_176, A_190, A_192, A_94, A_96, A_99). (E) Pocket 4 ( inter-protomer S2 pocket in CD region of S2. Protomer A residues A_1034, A_1035, A_1036, A_886, and protomer B residues B_1038, B_1040, B_1047, B_1107, B_1108, B_909). (F) Pocket 5 ( S2 pocket in UH region. Protomer A residues A_1056, A_1058, A_730, A_731, A_733, A_775, A_778, A_782, A_863, A_865, A_867, A_870). The predicted pockets are shown in red surface.

**Table S1.** The list of the intermolecular contacts in the structure of SARS-CoV-2 BA.2 RBD-ACE2 complex (pdb id 7XB0).

| **ACE2 Residue** | **ACE2 Residue Number** | **ACE2 Chain ID** | **RBD Residue** | **RBD Residue Number** | **RBD Chain ID** |
| --- | --- | --- | --- | --- | --- |
| ASP | 355 | A | GLY | 502 | B |
| TYR | 83 | A | ASN | 487 | B |
| GLN | 24 | A | ASN | 477 | B |
| ASP | 355 | A | THR | 500 | B |
| ASP | 30 | A | PHE | 456 | B |
| GLN | 42 | A | TYR | 449 | B |
| ASP | 38 | A | TYR | 501 | B |
| GLY | 354 | A | TYR | 501 | B |
| TYR | 83 | A | TYR | 489 | B |
| LEU | 45 | A | VAL | 445 | B |
| LEU | 45 | A | THR | 500 | B |
| LYS | 353 | A | TYR | 495 | B |
| GLN | 24 | A | PHE | 486 | B |
| HIS | 34 | A | LEU | 455 | B |
| LYS | 353 | A | GLY | 502 | B |
| LYS | 353 | A | THR | 500 | B |
| SER | 19 | A | ALA | 475 | B |
| LYS | 31 | A | LEU | 455 | B |
| LEU | 79 | A | PHE | 486 | B |
| GLN | 24 | A | TYR | 489 | B |
| TYR | 41 | A | ARG | 498 | B |
| ASN | 330 | A | THR | 500 | B |
| THR | 20 | A | ALA | 475 | B |
| LYS | 353 | A | VAL | 503 | B |
| ARG | 357 | A | THR | 500 | B |
| LYS | 353 | A | HIS | 505 | B |
| TYR | 41 | A | TYR | 501 | B |
| TYR | 83 | A | PHE | 486 | B |
| GLU | 35 | A | ARG | 493 | B |
| ASP | 38 | A | ARG | 498 | B |
| LEU | 79 | A | GLY | 485 | B |
| GLY | 354 | A | GLY | 502 | B |
| GLN | 24 | A | ALA | 475 | B |
| GLY | 354 | A | THR | 500 | B |
| ASP | 38 | A | GLY | 496 | B |
| PHE | 28 | A | TYR | 489 | B |
| ALA | 25 | A | ASN | 487 | B |
| THR | 27 | A | TYR | 489 | B |
| THR | 27 | A | PHE | 456 | B |
| SER | 19 | A | GLY | 476 | B |
| HIS | 34 | A | TYR | 453 | B |
| GLY | 354 | A | VAL | 503 | B |
| ASP | 30 | A | ASN | 417 | B |
| MET | 82 | A | PHE | 486 | B |
| GLU | 37 | A | HIS | 505 | B |
| GLY | 354 | A | HIS | 505 | B |
| LEU | 45 | A | ARG | 498 | B |
| ASP | 38 | A | TYR | 449 | B |
| THR | 27 | A | ALA | 475 | B |
| ASP | 355 | A | TYR | 501 | B |
| LYS | 353 | A | ARG | 403 | B |
| HIS | 34 | A | ARG | 493 | B |
| GLN | 325 | A | GLN | 506 | B |
| TYR | 41 | A | THR | 500 | B |
| HIS | 34 | A | ARG | 403 | B |
| GLN | 24 | A | GLY | 476 | B |
| GLY | 352 | A | GLY | 502 | B |
| GLN | 24 | A | ASN | 487 | B |
| GLN | 42 | A | ARG | 498 | B |
| LYS | 31 | A | ARG | 493 | B |
| ASP | 38 | A | TYR | 495 | B |
| SER | 19 | A | ASN | 477 | B |
| LYS | 353 | A | TYR | 501 | B |
| ASP | 30 | A | LEU | 455 | B |
| LYS | 31 | A | TYR | 489 | B |
| LYS | 31 | A | PHE | 456 | B |
| LYS | 31 | A | GLY | 485 | B |
| HIS | 34 | A | ASN | 417 | B |
| THR | 27 | A | TYR | 473 | B |
| THR | 20 | A | ASN | 477 | B |
| GLY | 326 | A | THR | 500 | B |
| GLN | 42 | A | GLY | 446 | B |
| GLY | 352 | A | TYR | 501 | B |
| ASN | 330 | A | PRO | 499 | B |

**Table S2.** The list of the intermolecular contacts in the structure of SARS-CoV-2 BA.2.75 RBD-ACE2 complex (pdb id 8ASY).

| **ACE2 Residue** | **ACE2 Residue Number** | **ACE2 Chain ID** | **RBD Residue** | **RBD Residue Number** | **RBD Chain ID** |
| --- | --- | --- | --- | --- | --- |
| ARG | 357 | A | THR | 500 | B |
| LYS | 31 | A | PHE | 456 | B |
| GLU | 35 | A | GLN | 493 | B |
| ASP | 38 | A | TYR | 449 | B |
| ASP | 355 | A | GLY | 502 | B |
| LYS | 353 | A | GLY | 496 | B |
| GLU | 37 | A | HIS | 505 | B |
| LEU | 45 | A | THR | 500 | B |
| THR | 27 | A | ALA | 475 | B |
| ASN | 330 | A | THR | 500 | B |
| TYR | 83 | A | TYR | 489 | B |
| ASP | 30 | A | PHE | 456 | B |
| THR | 27 | A | TYR | 489 | B |
| ASP | 355 | A | THR | 500 | B |
| LYS | 353 | A | TYR | 495 | B |
| HIS | 34 | A | GLN | 493 | B |
| PHE | 28 | A | TYR | 489 | B |
| PHE | 28 | A | ASN | 487 | B |
| ASP | 38 | A | GLY | 496 | B |
| SER | 19 | A | ASN | 477 | B |
| LYS | 353 | A | HIS | 505 | B |
| ASP | 38 | A | TYR | 495 | B |
| LYS | 353 | A | GLY | 502 | B |
| SER | 19 | A | ALA | 475 | B |
| THR | 27 | A | PHE | 456 | B |
| GLY | 354 | A | TYR | 501 | B |
| GLN | 24 | A | ASN | 477 | B |
| LEU | 79 | A | PHE | 486 | B |
| GLN | 42 | A | ARG | 498 | B |
| TYR | 83 | A | ASN | 487 | B |
| LYS | 353 | A | TYR | 501 | B |
| LYS | 31 | A | LEU | 455 | B |
| GLY | 354 | A | THR | 500 | B |
| LYS | 353 | A | THR | 500 | B |
| TYR | 83 | A | PHE | 486 | B |
| GLN | 24 | A | TYR | 489 | B |
| LEU | 45 | A | VAL | 445 | B |
| ASP | 30 | A | LEU | 455 | B |
| LYS | 353 | A | ARG | 498 | B |
| HIS | 34 | A | ASN | 417 | B |
| ASP | 38 | A | TYR | 501 | B |
| GLY | 354 | A | HIS | 505 | B |
| GLY | 354 | A | GLY | 502 | B |
| GLN | 42 | A | TYR | 449 | B |
| TYR | 41 | A | TYR | 501 | B |
| MET | 82 | A | PHE | 486 | B |
| HIS | 34 | A | TYR | 453 | B |
| GLY | 352 | A | TYR | 501 | B |
| ALA | 25 | A | ASN | 487 | B |
| TYR | 41 | A | THR | 500 | B |
| SER | 19 | A | GLY | 476 | B |
| LYS | 31 | A | TYR | 489 | B |
| LYS | 31 | A | GLN | 493 | B |
| GLY | 354 | A | VAL | 503 | B |
| ASP | 355 | A | TYR | 501 | B |
| ASP | 38 | A | ARG | 498 | B |
| TYR | 41 | A | ARG | 498 | B |
| LEU | 45 | A | ARG | 498 | B |
| GLN | 24 | A | ALA | 475 | B |
| GLN | 24 | A | GLY | 476 | B |
| GLN | 42 | A | SER | 446 | B |
| HIS | 34 | A | LEU | 455 | B |
| THR | 27 | A | TYR | 473 | B |
| GLN | 24 | A | ASN | 487 | B |
| LYS | 31 | A | PHE | 490 | B |
| GLN | 325 | A | VAL | 503 | B |
| GLN | 24 | A | PHE | 486 | B |

**Table S3.** The list of the intermolecular contacts in the structure of SARS-CoV-2 XBB.1 RBD-ACE2 complex (pdb id 8IOV).

| ACE2 Residue | ACE2 Residue Number | ACE2 Chain ID | RBD Residue | RBD Residue Number | RBD Chain ID |
| --- | --- | --- | --- | --- | --- |
| LEU | 79 | A | GLY | 485 | B |
| ALA | 25 | A | ASN | 487 | B |
| HIS | 34 | A | ARG | 403 | B |
| LEU | 45 | A | THR | 500 | B |
| GLN | 24 | A | ALA | 475 | B |
| TYR | 83 | A | ASN | 487 | B |
| ASN | 330 | A | THR | 500 | B |
| ASP | 30 | A | ASN | 417 | B |
| ASP | 38 | A | SER | 494 | B |
| TYR | 41 | A | ARG | 498 | B |
| GLN | 42 | A | ARG | 498 | B |
| TYR | 83 | A | TYR | 489 | B |
| GLY | 354 | A | GLY | 502 | B |
| LYS | 31 | A | SER | 490 | B |
| HIS | 34 | A | TYR | 453 | B |
| LYS | 353 | A | GLY | 504 | B |
| TYR | 41 | A | THR | 500 | B |
| THR | 20 | A | ASN | 477 | B |
| LEU | 79 | A | SER | 486 | B |
| THR | 27 | A | ALA | 475 | B |
| LYS | 353 | A | GLY | 502 | B |
| ALA | 36 | A | ARG | 493 | B |
| MET | 82 | A | ASN | 487 | B |
| HIS | 34 | A | ASN | 417 | B |
| GLY | 326 | A | THR | 500 | B |
| GLN | 24 | A | TYR | 489 | B |
| ASP | 38 | A | TYR | 495 | B |
| GLY | 354 | A | TYR | 501 | B |
| LYS | 31 | A | TYR | 489 | B |
| GLY | 354 | A | HIS | 505 | B |
| GLN | 24 | A | GLY | 476 | B |
| PHE | 28 | A | TYR | 489 | B |
| ASP | 355 | A | GLY | 502 | B |
| TYR | 83 | A | SER | 486 | B |
| ASP | 38 | A | ARG | 498 | B |
| ASN | 330 | A | PRO | 499 | B |
| GLN | 42 | A | TYR | 449 | B |
| LYS | 353 | A | TYR | 501 | B |
| HIS | 34 | A | SER | 494 | B |
| LEU | 45 | A | PRO | 445 | B |
| LYS | 353 | A | HIS | 505 | B |
| LYS | 31 | A | LEU | 455 | B |
| LYS | 353 | A | VAL | 503 | B |
| GLN | 24 | A | ASN | 477 | B |
| ASP | 38 | A | TYR | 501 | B |
| LEU | 45 | A | ARG | 498 | B |
| ASP | 355 | A | TYR | 501 | B |
| LYS | 31 | A | ALA | 484 | B |
| LEU | 39 | A | ARG | 493 | B |
| LYS | 353 | A | TYR | 495 | B |
| HIS | 34 | A | ARG | 493 | B |
| MET | 82 | A | SER | 486 | B |
| LYS | 353 | A | GLN | 506 | B |
| LYS | 31 | A | PHE | 456 | B |
| GLY | 352 | A | TYR | 501 | B |
| ASP | 30 | A | LEU | 455 | B |
| LYS | 31 | A | CYS | 488 | B |
| GLY | 354 | A | THR | 500 | B |
| SER | 19 | A | GLY | 476 | B |
| GLN | 325 | A | GLN | 506 | B |
| PHE | 28 | A | PHE | 456 | B |
| ASP | 38 | A | TYR | 449 | B |
| HIS | 34 | A | GLU | 406 | B |
| ASP | 30 | A | PHE | 456 | B |
| THR | 27 | A | TYR | 473 | B |
| TYR | 41 | A | TYR | 501 | B |
| GLN | 42 | A | SER | 446 | B |
| THR | 27 | A | PHE | 456 | B |
| SER | 19 | A | ASN | 477 | B |
| GLN | 24 | A | ASN | 487 | B |
| ASP | 38 | A | ARG | 493 | B |
| ARG | 357 | A | THR | 500 | B |
| HIS | 34 | A | LEU | 455 | B |
| GLU | 35 | A | ARG | 493 | B |
| GLU | 37 | A | HIS | 505 | B |
| THR | 27 | A | TYR | 489 | B |
| ASP | 355 | A | THR | 500 | B |
| SER | 19 | A | ALA | 475 | B |
| ASP | 38 | A | GLY | 496 | B |
| GLY | 354 | A | VAL | 503 | B |
