## Supplementary figures and images for "Examining Functional Linkages Between Conformational Dynamics, Protein Stability and Evolution of Cryptic Binding Pockets in the SARS-CoV-2 Omicron Spike Complexes with the ACE2 Host Receptor: Recombinant Omicron Variants Mediate Variability of Conserved Allosteric Sites and Binding Epitopes"

### FigureS1.tif

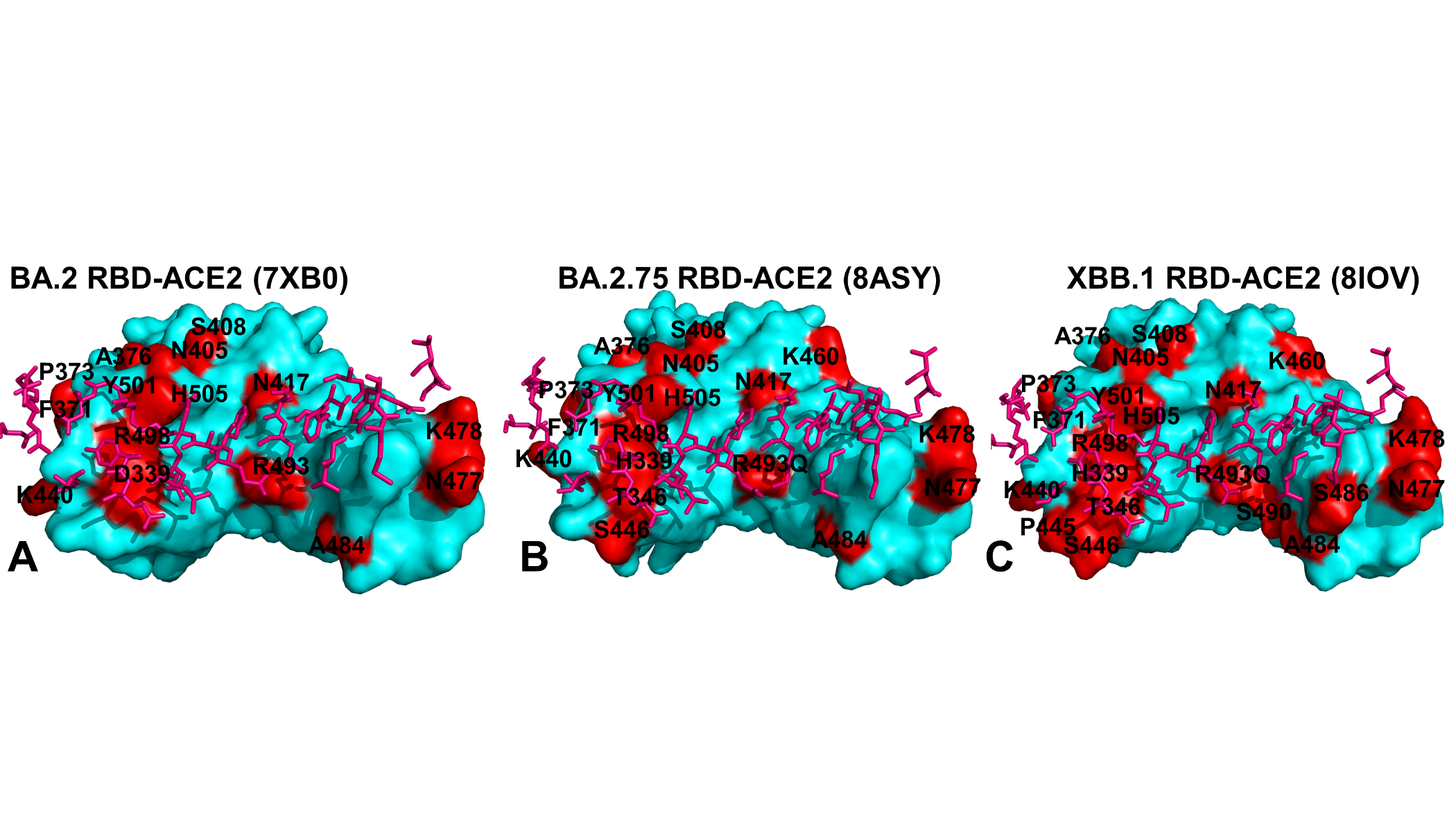

### FigureS2.tif

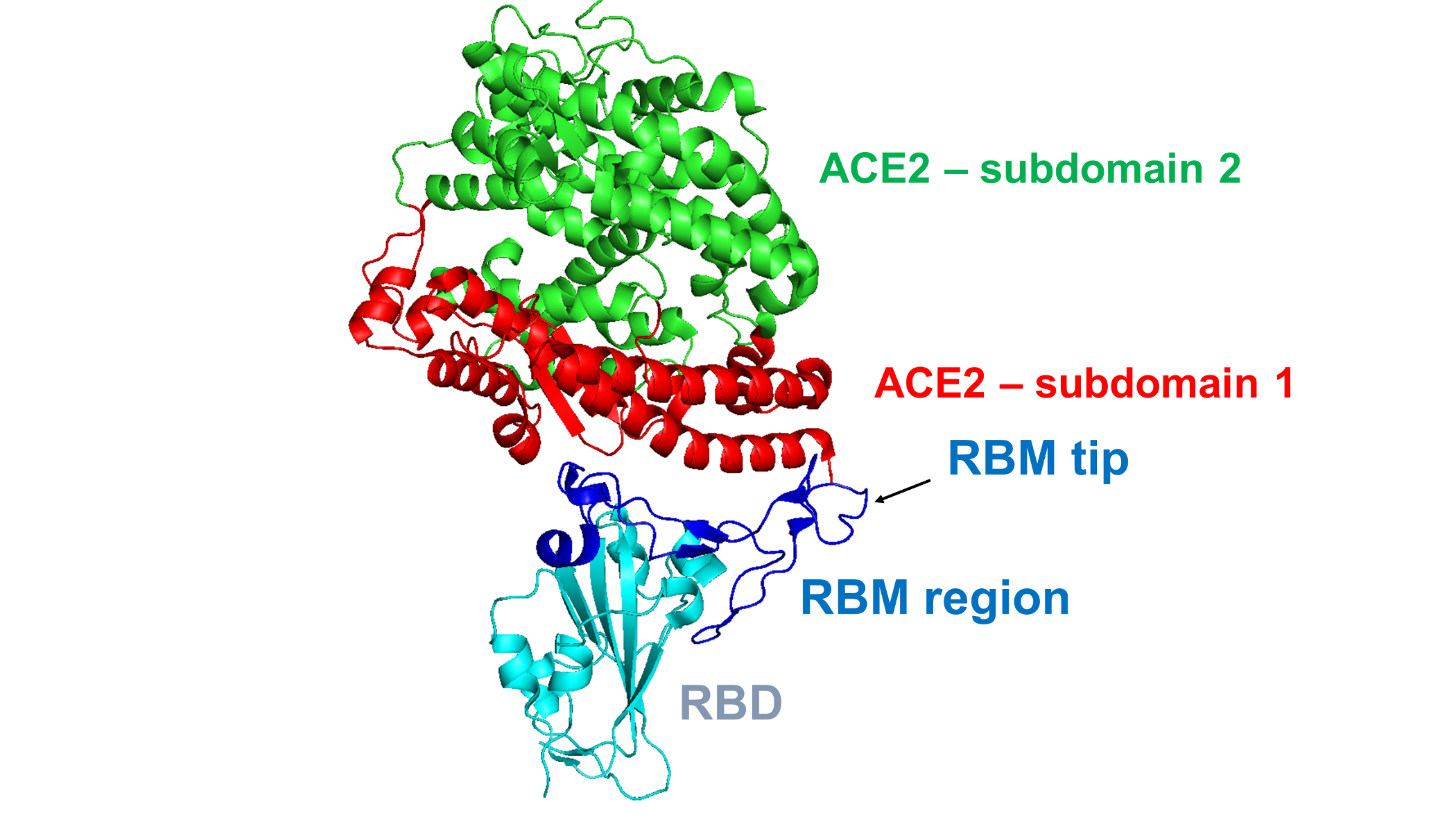

### FigureS3.tif

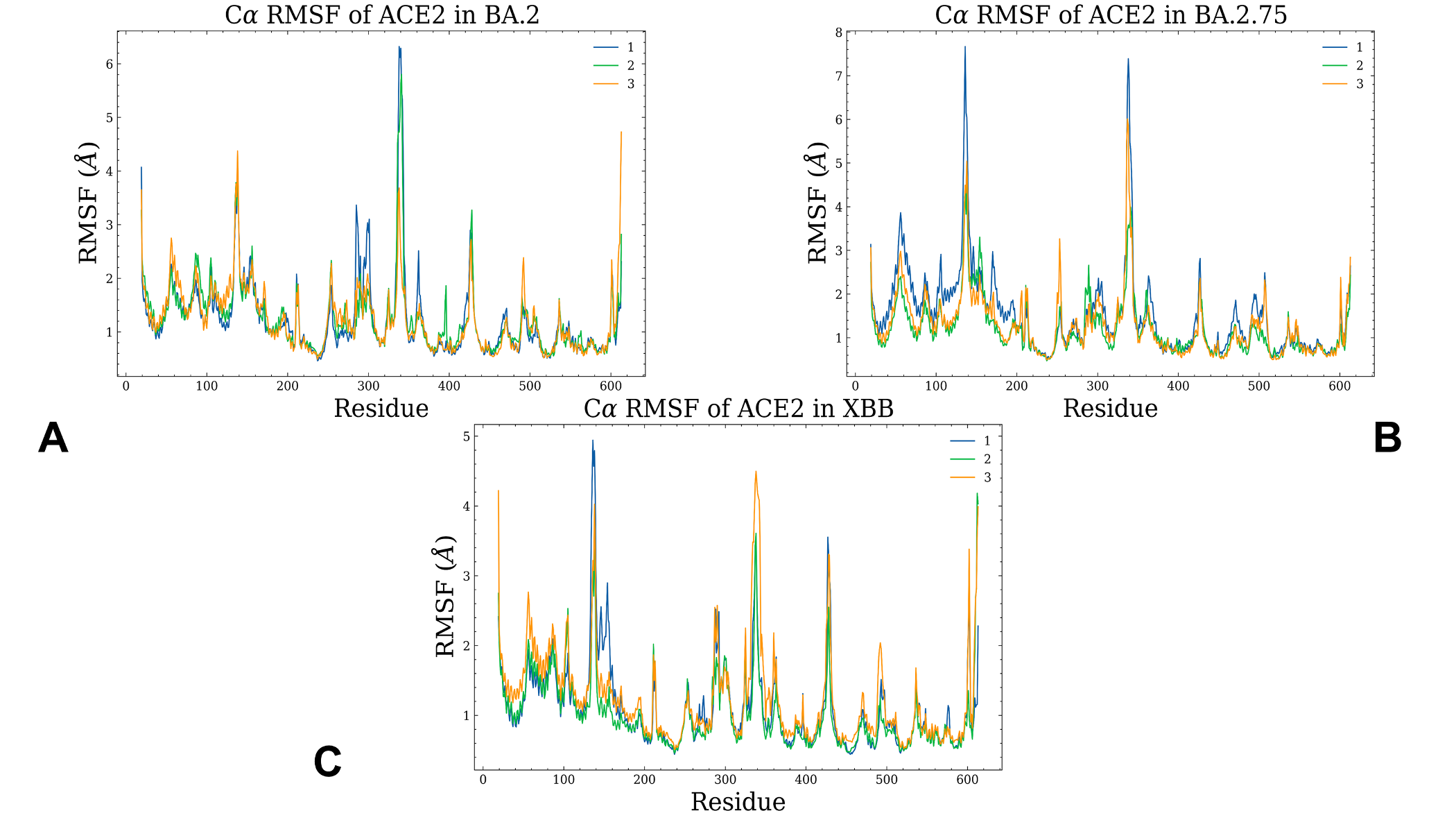

### FigureS4.tif

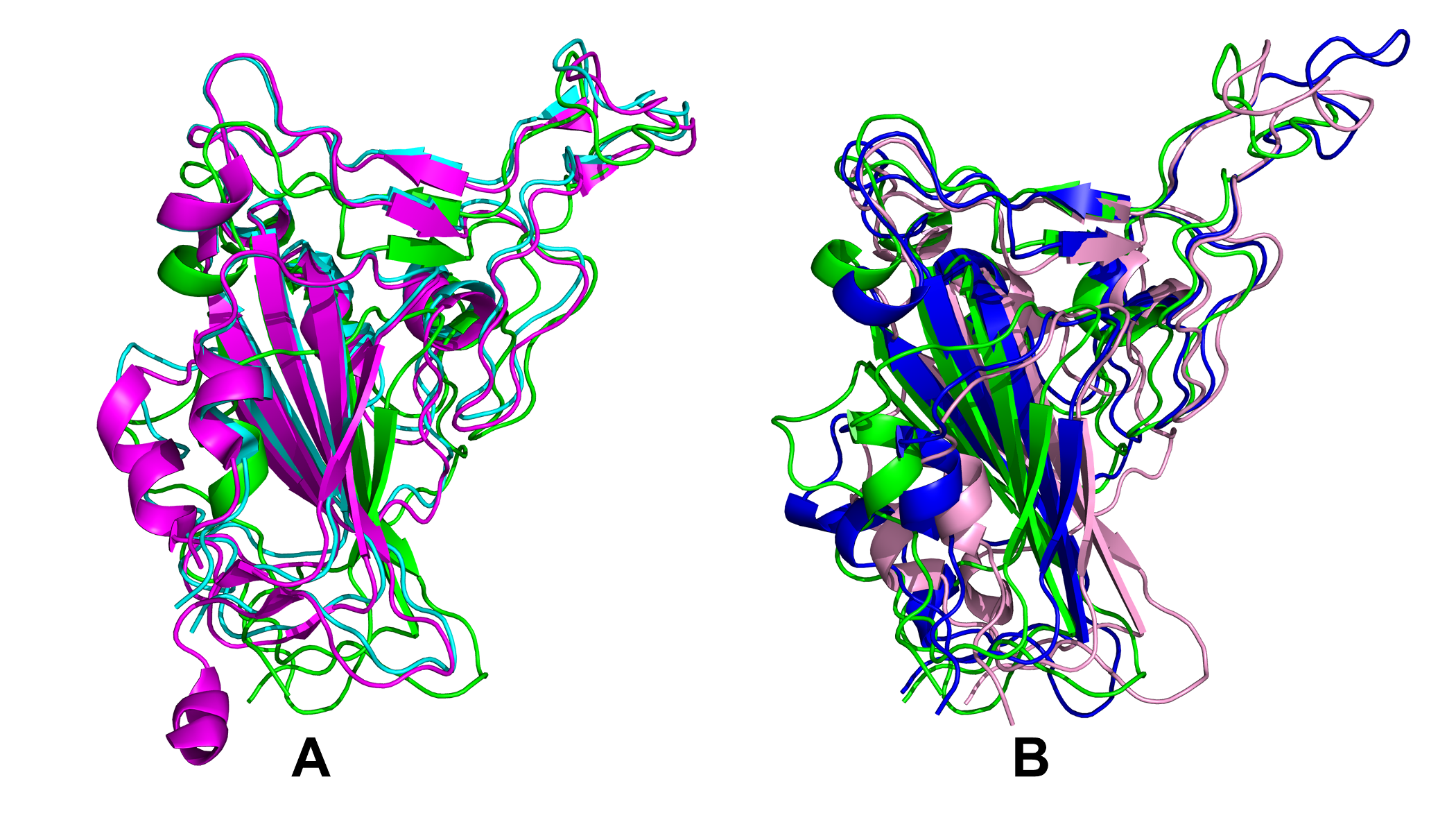

### FigureS5.tif

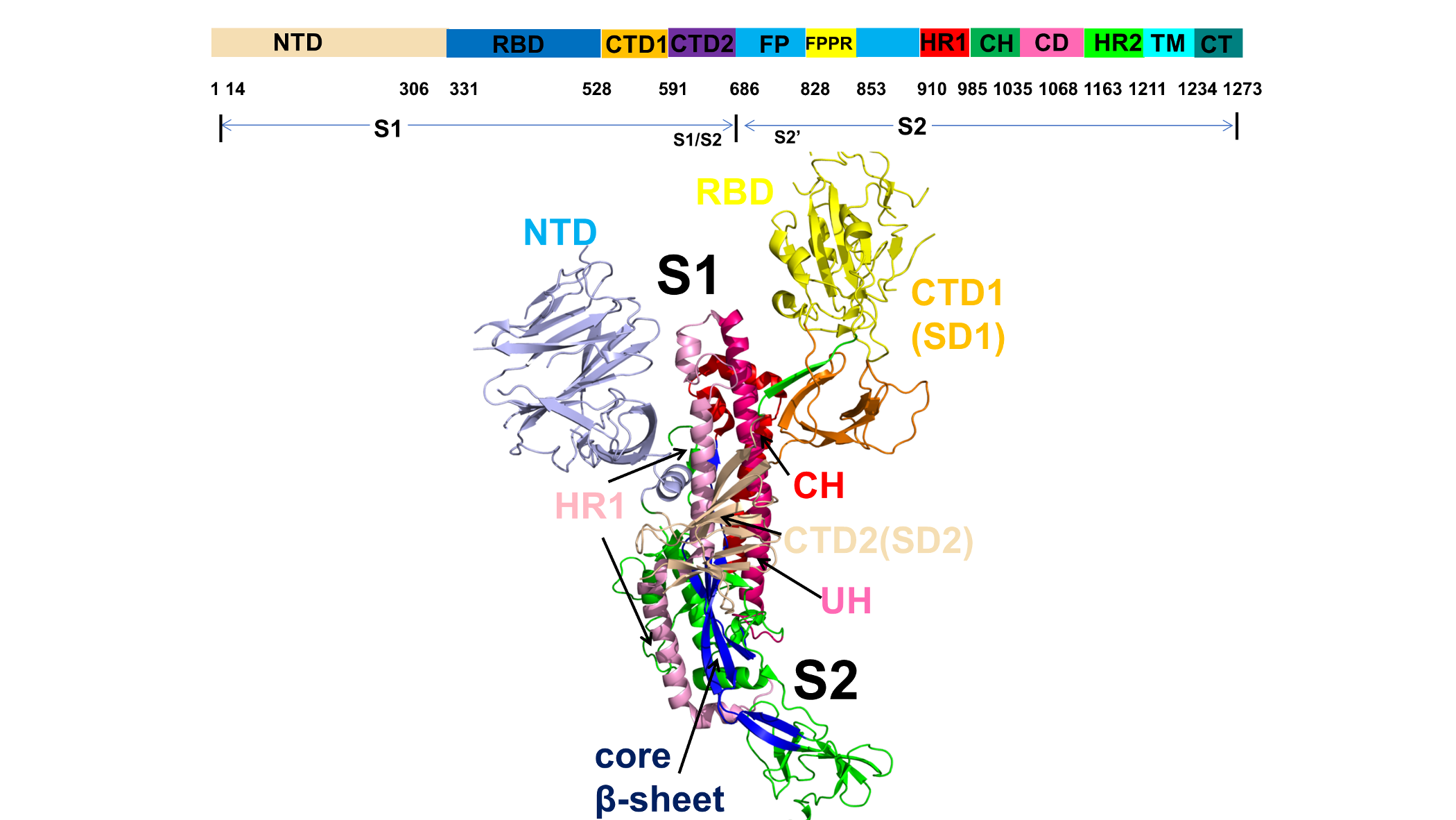

### FigureS6.tif

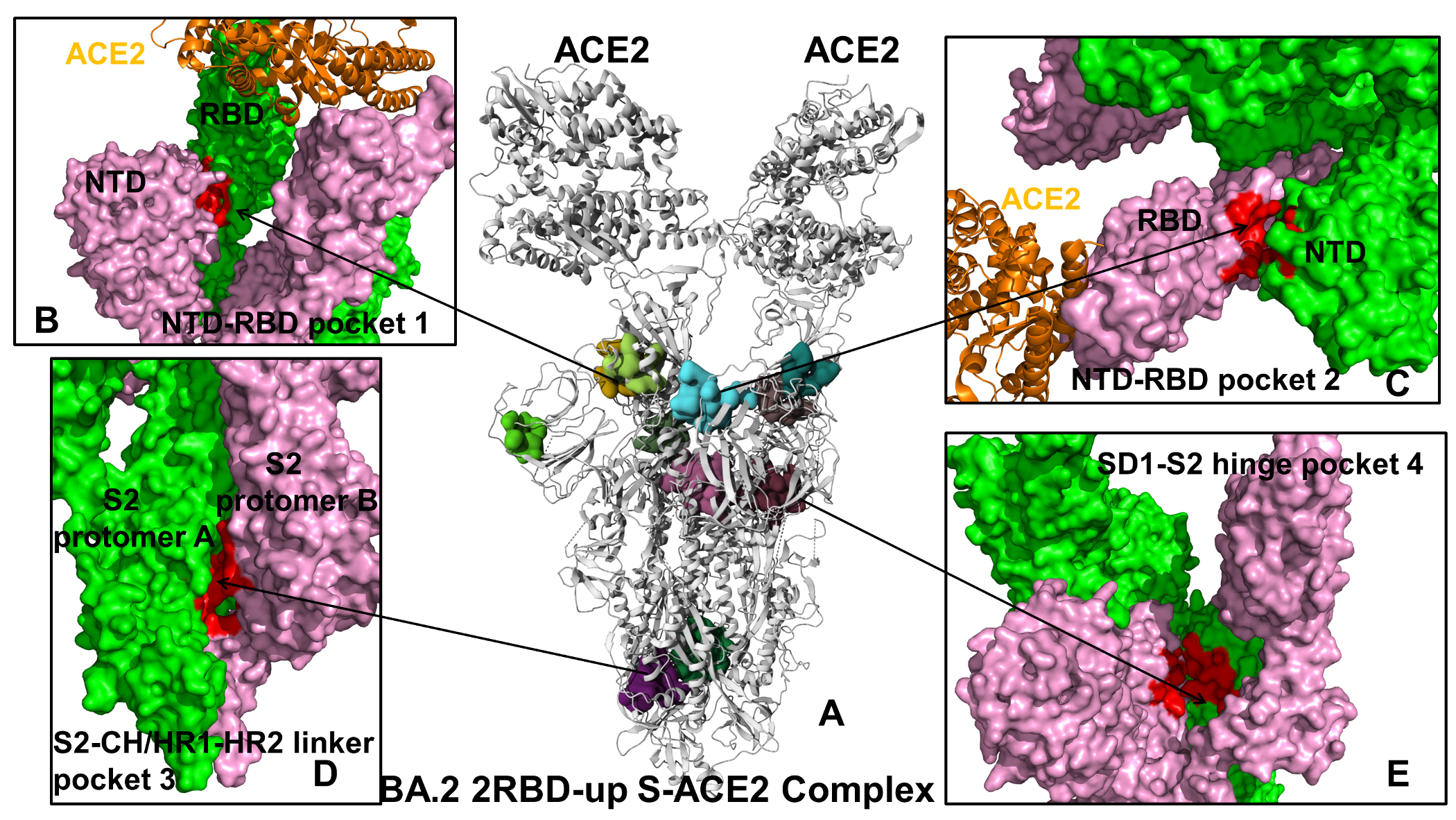

### FigureS7.tif

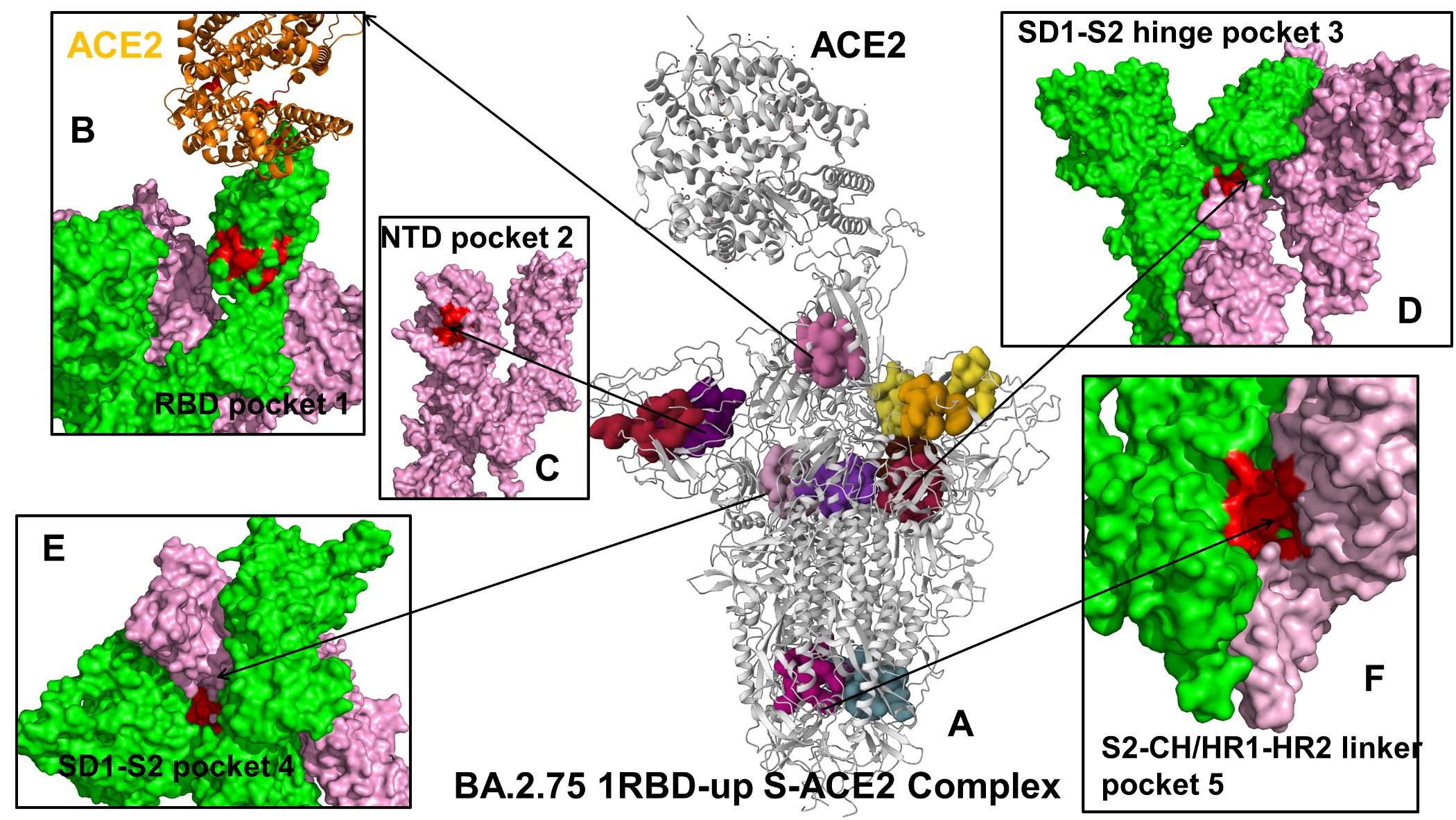

### FigureS8.tif

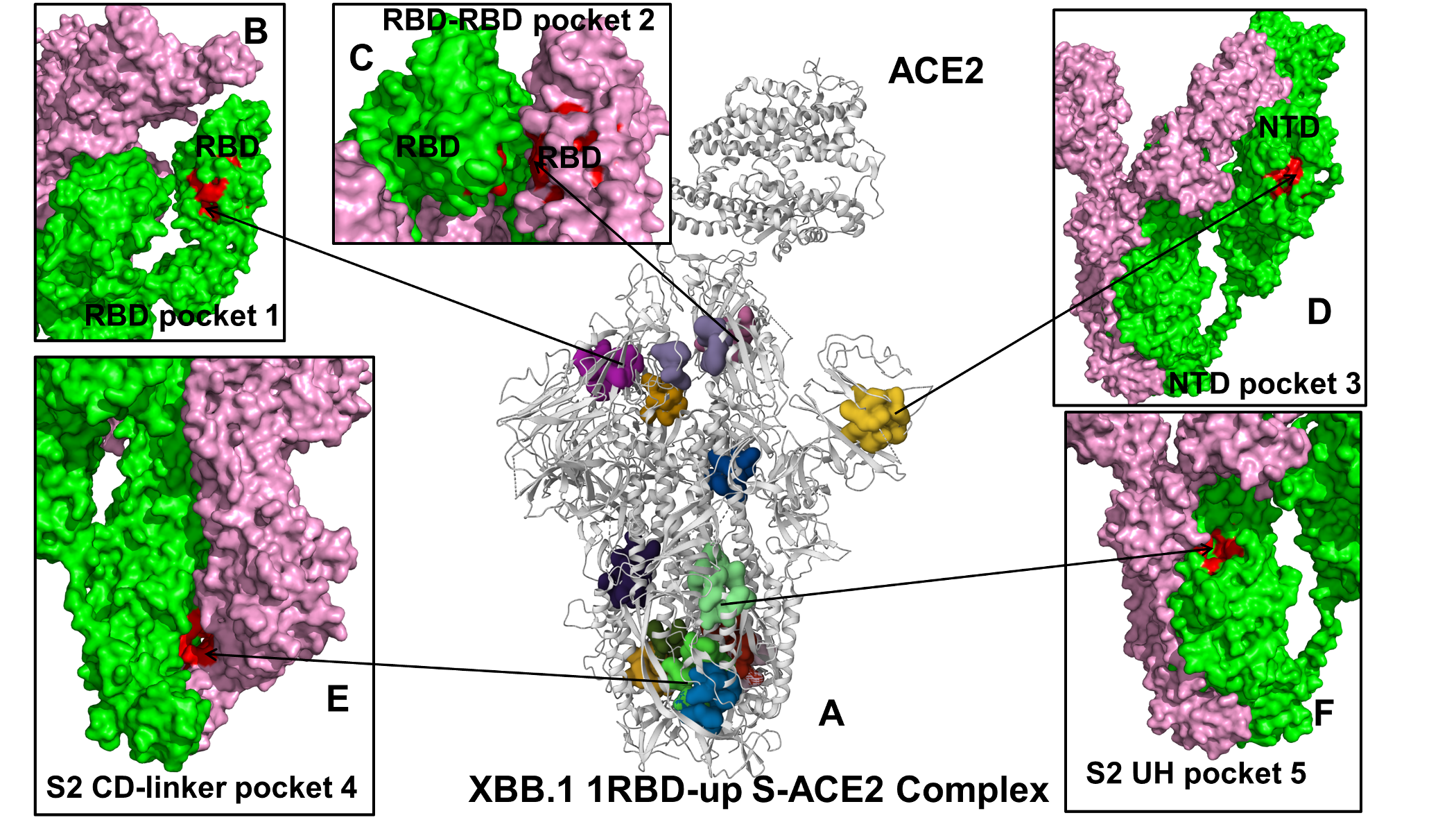
